# Host identity and fungal source pools shape mycorrhizal assembly under carbon removal interventions in large-scale afforestation

**DOI:** 10.64898/2026.10.26.761856

**Authors:** Isabelle J. Thomson, Oliver Lindsay, Heather Allen, Colin Averill, Thomas D. Brekke, Charles Nicholls, Brian Steidinger, Elena Vanguelova, Bonnie Waring, Laura M. Suz, Martin I. Bidartondo

## Abstract

1. Large-scale afforestation is central to global carbon dioxide removal (CDR) commitments. Soil inoculation and enhanced rock weathering (ERW) may increase tree biomass carbon stocks during woodland establishment. Both can alter soil fungal communities, but whether they change the root-colonising mycorrhizal fungi that support tree nutrition and growth is largely unknown.
2. In a fully-factorial field trial on former upland pasture in Wales (11.5 ha; ∼25,000 saplings), we tested the effects of soil inoculation (addition of soil from nearby mature, donor forests) and ERW (basalt amendment) on root-associated ectomycorrhizal (EcM) and arbuscular mycorrhizal (AM) fungi, and tree growth in mixed-broadleaf and conifer plantations, two and four years after planting. To identify key colonisation sources, we also characterised mycorrhizal fungi in roots and soils of surrounding hedgerows, source tree nursery, non-forested control plots, donor woodlands and soil inoculum.
3. Soil inoculation caused a subtle shift in conifer EcM community composition (3% of variation) but had no detectable effect in broadleaves. Basalt amendment raised soil pH but did not change mycorrhizal composition, diversity, or colonisation.
4. Root-associated EcM communities were distinct from those detected in plantation soils and shared far more taxa with the tree nursery than with the donor inoculum (41-48% vs. 2-9% of plantation EcM taxa) whereas AM communities largely reflected fungi present in pre-existing pasture.
5. Host identity was the dominant factor structuring mycorrhizal communities, overshadowing both treatments, and was the strongest predictor of aboveground biomass.
6. *Synthesis and applications*: This field-scale evaluation shows early mycorrhizal assembly during woodland establishment is strongly structured by tree species identity and pre-existing fungal source pools, limiting the effect of inoculation at planting. Mycorrhizal management should therefore target the nursery stage and be tailored to individual tree species. Basalt amendment for ERW is unlikely to harm mycorrhizal communities during establishment. Because soil fungal communities did not reflect root symbiosis, our findings also stress that monitoring of belowground outcomes in CDR-focused planting should include root sampling rather than relying on soil DNA alone.

## 1 Introduction

Large-scale afforestation and reforestation are central to national (Forster et al., 2021) and global (Rockström et al., 2017) commitments on carbon dioxide (CO_2_) removal (CDR) for climate change mitigation. As forests mature, they can become net carbon sinks, removing atmospheric CO_2_ through photosynthesis and forming long-term carbon stores in their biomass, litter and soils (Kafy et al., 2023; Psistaki et al., 2024). However, newly established woodlands can remain net carbon sources for decades after planting, as tree nursery and planting operations, soil disturbance and seedling mortality can drive initial carbon losses (Banin et al., 2022; Bárcena et al., 2014; Hong et al., 2023; Laganiere et al., 2010). Whether newly planted woodlands contribute meaningfully to short-term climate targets depends on how quickly they transition from carbon source to sink.

Mycorrhizal fungi facilitate host acquisition of soil nutrients, particularly nitrogen (N), phosphorus (P), and water in exchange for photosynthetically fixed carbon, influencing tree establishment, growth and resilience (Augé et al., 2014; Brundrett et al., 1996; Hawkins et al., 2023; Jung et al., 2012; Tourville et al., 2024), and potentially the rate at which developing forests transition from carbon sources to carbon sinks. Arbuscular mycorrhizas (AM) and ectomycorrhizas (EcM) dominate terrestrial ecosystems and differ in their physiological and ecological traits (Tedersoo & Bahram, 2019). Root-associated ectomycorrhizal community composition is associated with tree growth and biomass carbon stocks across European forests (Anthony et al., 2022), and with soil carbon stocks at the landscape scale (Barber et al., 2026). The success of tree plantation for CDR can therefore be influenced by the assembly and functioning of mycorrhizal fungi.

Soil inoculation and enhanced rock weathering (ERW) have both been proposed as strategies for accelerating CDR in afforestation and reforestation (Averill et al., 2022; Gaucher et al., 2025). However, little is known about how soil inoculation and enhanced rock weathering affect mycorrhizal community assembly during woodland establishment.

Soil inoculation is intended to address microbial legacies of previous land use. Afforestation sites are often characterised by depleted soil microbial diversity and mycorrhizal inoculum, reflecting land-use legacies such as long-term agriculture (King et al., 2023; Riedo et al., 2021; Yuan et al., 2025). Natural colonisation and diversity of mycorrhizal fungi in planted seedlings can also decline with distance from an established woodland (Dickie & Reich, 2005; Sara Grove et al., 2019; Teste et al., 2009). Thus, simply planting trees may not rapidly recover the microbial legacies of previous land use (Azevedo et al., 2026). Whole soil inoculation aims to introduce intact microbial communities from nearby mature woodlands (Duell et al., 2023; Gomes et al., 2025; Leff et al., 2015; Wubs et al., 2016; Zang et al., 2024). In planted trees this can increase mycorrhizal colonisation, fungal richness, alter fungal community composition (Lance et al., 2019; Wang et al., 2023) and increase seedling growth (McMahen et al., 2022; Taniguchi et al., 2021), but its effects vary among host species, inoculum sources and environmental conditions (Lance et al., 2019; Parasquive et al., 2024; St-Denis et al., 2017). Some field experiments have found changes in fungal community composition without corresponding improvements in seedling performance (Bashian-Victoroff et al., 2026; Wasyliw et al., 2023), while others have found little or no effect on either (Grove et al., 2019; Rodriguez-Ramos et al., 2022). Consequently, we still have a limited understanding of what determines whether fungi introduced in donor soil successfully establish on planted trees.

Enhanced rock weathering is a CDR strategy in which silicate rocks, such as basalt, are applied to soils, where weathering converts gaseous CO_2_ to bicarbonate that can be exported to the oceans, generating long-term CDR. Research has focused on agricultural systems (Beerling et al., 2020; Clarkson et al., 2024), although ERW may also enhance forest productivity and CDR (Gaucher et al., 2025). Basalt can also alter soil pH, mineral nutrients availability and water retention (Anda et al., 2015; Conceição et al., 2022; Goll et al., 2021; Lewis et al., 2021; Truog, 1947), all of which are ecological filters of both mycorrhizal community composition (Davison et al., 2021; Gosling et al., 2013; van der Linde et al., 2018) and colonisation (Carrino-Kyker et al., 2016; Gryndler et al., 2006; Liu et al., 2021). However, little is known about how ERW affects mycorrhizal community assembly during woodland establishment.

Both whole soil inoculation and ERW therefore have the potential to modify mycorrhizal communities, with implications for CDR and afforestation success. We investigated these effects in a fully-factorial field experiment comprising a plantation of ∼25,000 EcM and AM trees on 11.5 ha of former upland pasture (>100 years) in Wales (Fig. 1). At this site, both strategies have been shown to alter soil fungal communities and tree growth (Waring et al., 2026). However, plant hosts selectively recruit fungi from the available soil fungal pool, meaning bulk soil communities do not necessarily reflect those colonising roots, and the two may respond differently to changing soil conditions (Goldmann et al., 2016; Frew et al., 2025; Fu et al., 2026). We therefore characterised root-associated EcM and AM communities, and AM colonisation of the plantation. We integrated these data with plantation aboveground biomass, used here as an early-stage proxy for carbon sequestration outcomes. Finally, to assess the contribution of soil inoculum to plantation communities relative to other potential sources of colonisation, we compared plantation root and soil mycorrhizal communities with donor woodland inoculum sources, surrounding hedgerows, the source tree nursery and non-forested pasture controls.

**Figure 1.**
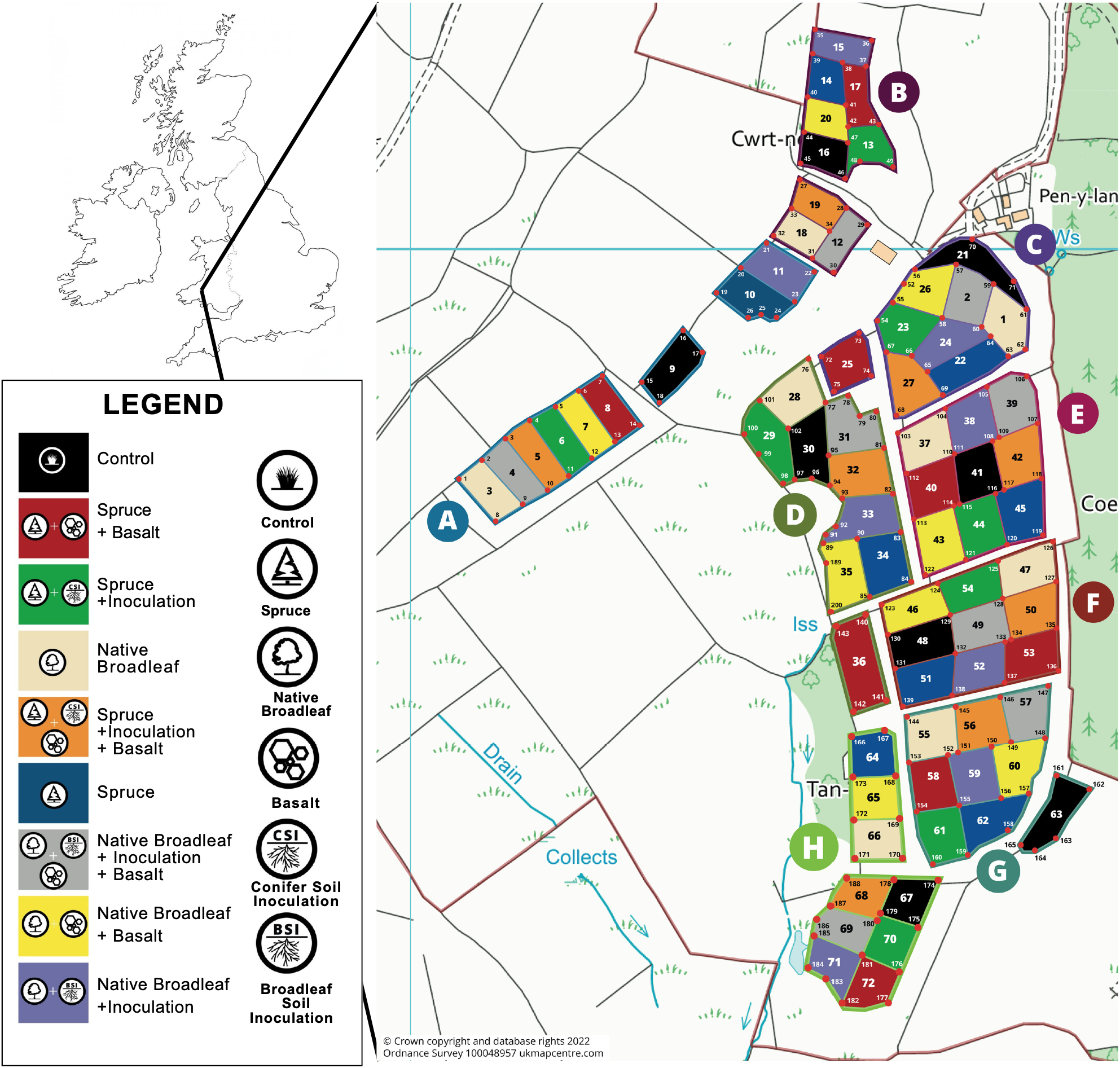
Map of The Carbon Community project experimental field trial, Glandwr Forest, Cynghordy, Wales. This includes the 72 plots and their corresponding treatment (s) (11.52 ha; ∼25,000 trees). Letters indicate blocks, and numbers indicate plot. Adapted from Waring et al., (2026).

We hypothesised: (H1) Soil inoculation would alter root-associated mycorrhizal community composition, especially EcM; (H2) inoculated EcM communities contain taxa attributable to donor woodland, whereas AM communities retain a pasture legacy; (H3) basalt amendment would alter mycorrhizal community composition through changes to soil chemistry; and (H4) that variation in EcM community composition would be associated with plantation tree biomass.

## 2 Materials and Methods

### 2.1 Experimental site and design

The study was conducted at The Carbon Community experimental afforestation site in Glandwr Forest, Cynghordy, Wales (52.0636°N, 3.7449°W), established in 2020. The site has a mean annual temperature of 9.6°C, and precipitation of 1,566 mm (Met Office, 2026). The plantations are established on Manod series soils. Prior to afforestation, the land had been managed as upland sheep pasture for at least 100 years.

The experiment comprises a fully-factorial field trial covering 11.52 ha, divided into eight experimental blocks (Fig. 1; 1.44 ha each). Each block contains nine plots (approximately 40 × 40 m), for a total of 72 plots. A timeline of site establishment, treatment application and sampling is provided in Fig. 2. Approximately 3,200 two-year-old saplings per block were planted by hand into mounds between 7 May and 11 June 2021, supplied by Tilhill Forestry Maelor Nursery (tilhill.com; Wrexham, Wales).

**Figure 2.**
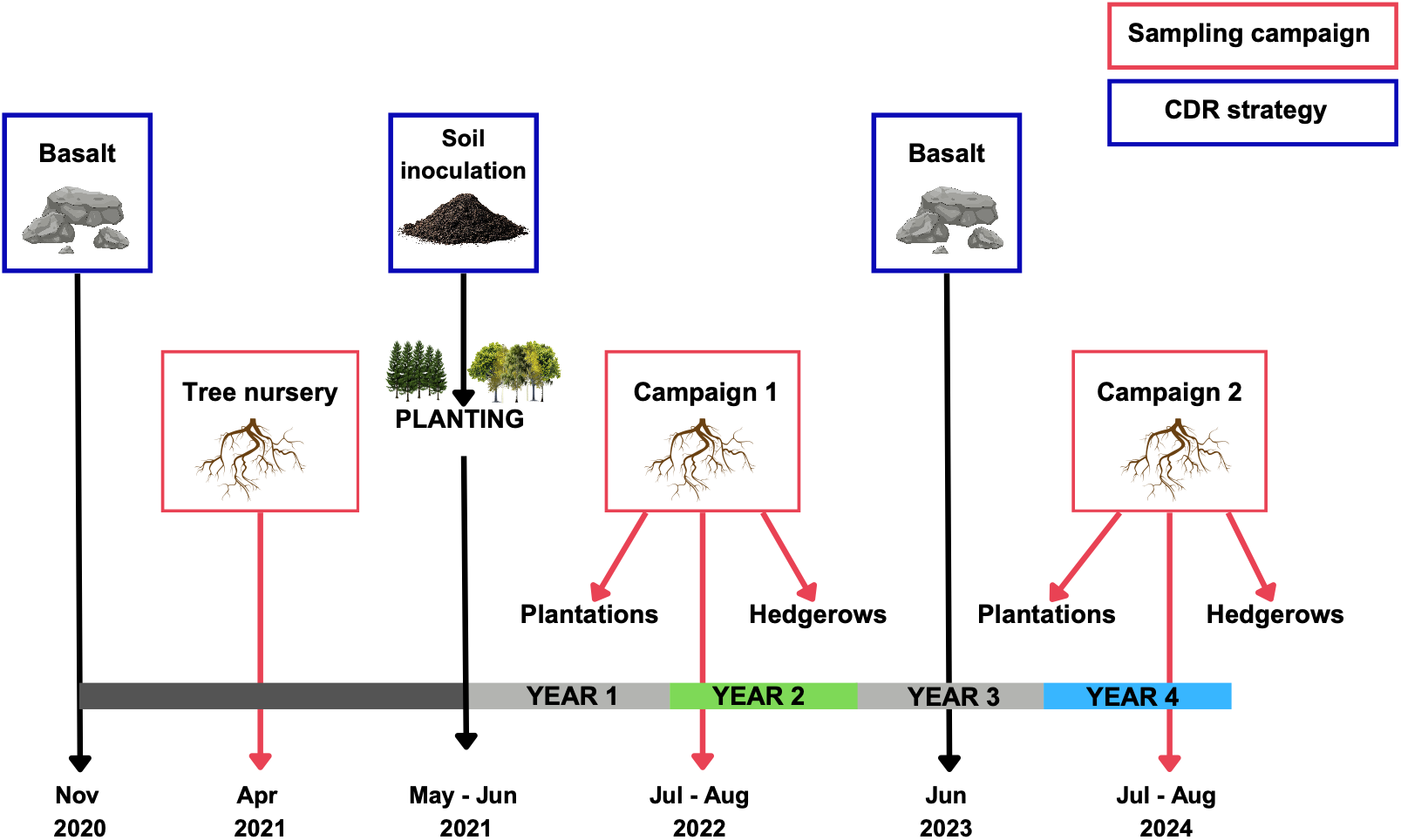
Experimental timeline for The Carbon Community project, illustrating the timing of treatment applications (dark blue) and sampling campaigns (red) throughout the study. Root-associated fungal communities were sampled from the tree nursery, plantation plots, and adjacent hedgerows.

The experiment followed a factorial design with plantation type, and basalt and soil inoculation treatments, as the main experimental factors. Plantation type consisted of either conifer monoculture - non-native EcM Sitka spruce (*Picea sitchensis;* hereafter, spruce) plantation or a mixed broadleaf plantation with both native EcM and AM tree species (Table S1) - alder (*Alnus glutinosa, 22%*), downy birch (*Betula pubescens, 35%*), wild cherry (*Prunus avium, 19%*), sessile oak (*Quercus petraea, 13%*), rowan (*Sorbus aucuparia, 6%*), and aspen (*Populus tremula, 5%*), in compliance with the UK Forestry Standard.

Within each block, plots were assigned to one of four treatments for each plantation type: (i) untreated control (no inputs), (ii) soil inoculation, (iii) basalt amendment, or (iv) combined soil inoculation and basalt amendment. In addition, each block contained a non-forested pasture control, resulting in nine plots per block.

### 2.2 Treatment design

Basalt was applied in November 2020 prior to planting (at a rate of 40 t ha^-1^; 2-4 mm particle size) and reapplied in June 2023 (24 t ha^-1^; 0-2mm; Fig. 2). Soil inoculation consisted of applying 200 ml of fresh woodland soil directly to the roots of each tree during planting. Inoculum for the broadleaf plantation was collected from a nearby native broadleaf woodland in Rhandirmwyn (ca. 7 km from the experimental site). Inoculum for the spruce plantation was collected from a mature spruce plantation near Carmarthen, approximately 48 km from the experimental site. Soil was collected from 10-15 cm beneath the litter layer and between 3-120 hours before planting.

### 2.3 Root sampling

Mycorrhizal roots were sampled in experiment year 2 and year 4, after treatment applications. Within each plot, target trees nearest the plot centre were selected and four soil cores (2.5 × 30 cm) were removed 20-25 cm from the stem base.

For broadleaf plots, sampling of target trees reflected the planting ratio of EcM and AM tree species, with two EcM and one AM tree sampled per plot. Sampling groups are represented on (Table S1). Two spruce trees (EcM) were sampled per plot in all eight blocks, matching the per-plot EcM sampling effort in the broadleaf plantation.

To identify potential sources of mycorrhizal fungi colonising plantation roots, we sampled roots from the source Maelor Nursery prior to planting (nursery-grown seedlings), woodland inoculum sources (sites from which donor soil was obtained), surrounding hedgerows and non-forested pasture controls. Further detail is provided in S1.

### 2.4 Mycorrhizal assessment

Roots were washed from soil cores (for the plantation samples, the four cores taken per stem were pooled), and twelve roots were randomly selected from each sample using a grid-based approach and processed according to mycorrhizal type. For EcM hosts, one colonised root tip (mycorrhiza) was selected, host identity verified by root morphology, and morphological traits associated with soil exploration type were recorded (Agerer, 2001) before DNA extraction (Extract-N-Amp Plant PCR Kit; Sigma-Aldrich). For AM plantation hosts and pasture controls (mostly grasses), each root was divided into two 1 cm segments, one was pooled per sampled for DNA extraction (DNeasy 96 Plant Kit; QIAGEN) and the other was stained for colonisation assessment following methods from Kowal et al. (2020). Nursery seedlings were sampled by morphotype (EcM hosts) or pooled by tree species (AM hosts) (For further details, see S1.7; Sample totals Table S2).

### 2.5 Molecular analysis of mycorrhizal fungi

EcM fungi were identified by Sanger sequencing of the ITS region (primers ITS1F/ITS4; Gardes & Bruns, 1993; White et al., 1990) from individual ectomycorrhizas (Table S4). Sequences were processed with the MycorrhizaTracer pipeline (Brekke et al., 2026), which uses VSEARCH (Rognes et al., 2016) to match sequences against UNITE v10 (Abarenkov et al., 2024). Sequences matching ≥97% to a reference were assigned the corresponding SH (species hypothesis); the rest were clustered into *de novo* OTUs at 97% similarity. Taxonomy was assigned by BLASTN (Sayers et al., 2025), with assignments made to species (≥97% identity), genus (94.0-96.9%), family (91.0-93.9%), and order or higher (<91.0%; Deslippe et al., 2011). Low-quality sequences were removed and non-EcM taxa were excluded using FUNGuild (vGuilds_v1.1; Nguyen et al., 2016), giving one EcM taxon assigned to each mycorrhiza.

AM fungi were identified from pooled root DNA by PacBio long-read sequencing of the partial SSU and full ITS region (primers NS1B1ngs/ITS4ngsUni; Tedersoo & Anslan, 2019; Tedersoo & Lindahl, 2016; Table S4), allowing detection of multiple AM taxa per sample. Reads were processed with nf-core/ampliseq (Straub et al., 2020), clustered into 97% OTUs and assigned taxonomy against UNITE v10, retaining only Glomeromycota and Endogonales as AM fungi (Hoysted et al., 2018). For further details, see S1.8-9.

### 2.6 Arbuscular mycorrhizal colonisation assessment

Stained, mounted root sections were scanned using an Epredia 3DHISTECH Pannoramic MIDI II Rx brightfield digital slide scanner (37x objective; 0.27 μm pixel^−1^). Individual root images were uploaded onto QuPath (v0.6.0) for colonisation assessment. A detailed description of the workflow and annotation scheme is in S4. Arbuscular mycorrhizal colonisation was assessed for year 2 root samples only. Year 4 roots were not assessed for colonisation, as older roots were too woody to clear and stain reliably.

### 2.7 Soil mycorrhizal fungal communities

Raw sequence data from DNA extractions of the plantation soils, soil inoculum, and pasture control soils were obtained from Waring et al. (2026). Soils had been collected from the rooting zone of trees in year 2. Total DNA was extracted and the ITS region (primers ITS9munngs and ITS4ng; Table S4) was amplified and sequenced using circular consensus sequencing on a PacBio platform. To identify mycorrhizal fungi in these soils, sequences were processed using the nf-core/ampliseq pipeline, as previously described.

### 2.8 Plantation pH and biomass measurements

Soil pH was measured in year 4, as described in S1.11. Plantation biomass data were retrieved as described by Waring et al. (2026) (S1.12). Because individual biomass measurements could not be matched to the trees sampled for mycorrhizal analyses, biomass was averaged for each tree species within each plot before analysis.

### 2.9 Community assessment and statistical analyses

All analyses were performed on R (v4.3.3) (R Core Team, 2020), package version and additaionl information are listed in S1.13. Mycorrhizal communities were assembled at plantation level (broadleaf vs spruce) and tree-species level, with each community defined by the combination of plantation type or tree species, block, treatment, and year of sampling. After excluding EcM communities with fewer than five successful DNA sequences, and uncolonised AM roots, analyses included 127 EcM and 62 AM communities at the plantation level, and 179 EcM and 62 AM communities at the tree species level across both sampling years (Table S3). EcM data were converted to relative and analysed using Bray-Curtis dissimilarity; AM read counts were converted into presence-absence to limit PCR bias and analysed using Jaccard (AM) dissimilarity.

Treatment effects on community composition (H1 & H3) were tested with PERMANOVA (vegan; Oksanen et al., 2013), with basalt, inoculum, sampling year and host species as predictors, and permutations restricted to reflect the split-plot sampling design (S1.14). Multivariate dispersion was tested with PERMDISP (Anderson & Walsh, 2013), and variance partitioned among predictors (*varpart)*, with unique fractions tested by distance-based redundancy analysis (db-RDA) following the same permutation schemes (S1.14). Treatment effects were also tested on principal coordinates analysis (PCoA) axis scores using linear mixed-effects models (lme4; Bates et al., 2015). Indicator taxa were identified with indicspecies (De Cáceres & Legendre, 2009) and alpha diversity was estimated as Hill numbers (iNEXT; Hsieh et al., 2016). Treatment effects on AM colonisation in cherry and rowan were tested using a beta-binomial GLM (glmmTMB), weighted by the number of root regions assessed.

The contribution of each potential colonisation source to plantation communities (H2) was measured as asymmetric overlap: the proportion of taxa in a focal community (A) also detected in a comparison community (B): 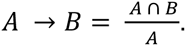 Overlaps were visualised with *eulerr* (Larsson, 2024).

Relationships between EcM community composition and plantation biomass (H4) were tested separately for each year using linear mixed-effects models of square-root-transformed biomass, with PCoA axes 1 and 2, plantation type, host species, and treatment as fixed effects, and plot and block as random effects. Because host identity strongly structured composition, a second set of models used PCoA scores centred within each tree species, so that composition coefficients reflected only within-host variation, independent of host identity (S1.13). Models were selected by AIC and likelihood ratio tests. EcM taxa associated with biomass were identified using per-OTU binomial GLMs (presence ∼ biomass) with Benjamini– Hochberg correction.

## 3 Results

### 3.1 Composition of mycorrhizal fungal communities

Of the 4,607 ectomycorrhizas collected across the plantation plots and potential sources of colonisation, 3,682 yielded a successful EcM fungal identification (ID), representing, 203 EcM OTUs. After quality filtering, 203 EcM OTUs remained, of which 130 matched a UNITE SH and 73 OTUs were identified as *de novo* (Tables S2 & S5). The spruce plantation yielded 53 EcM OTUs, comprising 52% Basidiomycota and 48% Ascomycota, with Ascomycota accounting for 70% of ectomycorrhizas and *Trichophaea* (57%), *Amphinema* (16%) and *Hebeloma* (11%) the dominant genera. Broadleaf roots hosted 93 EcM OTUs, predominantly Basidiomycota (74% of OTUs, 77% of ectomycorrhizas), with *Thelephora* (37%), *Hebeloma* (25%) and *Peziza* (17%) the most abundant genera.

Of the 971 fungal OTUs sequenced from 843 AM root sampled, 63 were AM fungi, all occurring in the broadleaf plots and 40 also found in the pasture controls (Table S6). Glomeromycetes dominated AM reads (77%), with *Dominikia* (23%) and *Glomus* (16%) the most common general. Year-specific richness and communities from surrounding hedgerows, woodland soil inoculum sources, tree nursery and soils are described in S3.1, and OTUs are given in S5.

### 3.2 (H1) Soil inoculation shifted spruce but not broadleaf EcM communities

Soil inoculation had a small but significant effect on spruce EcM fungal community composition (PERMANOVA n= 64, pseudo-(p) F_1,60_ = 2.213, R^2^= 0.032, *p* = 0.009), with no difference in dispersion (PERMDISP; *p* = 0.202). Inoculation accounted for 2.1% of explained variance in EcM communities (db-RDA; *p* = 0.015; Fig. 3). Sampling year explained more variation (PERMANOVA n = 64, p-F_1,60_= 7.137, R^2^ = 0.102, *p* = 0.001), accompanied by a reduction in among-plot dispersion (PERMDISP *p* = 0.001; mean distance to median 0.49 vs 0.33), indicating EcM communities converged over time. There was no year x treatment interaction (PERMANOVA n = 64, p-F_1,60_ = 0.536, R^2^ = 0.008, *p* = 0.800; Table S10). Communities separated by year along the PCo1 and PCoA2, and by inoculum along the PCoA2 and PCoA3 (Fig. 4). *Trichophaea* sp. 2 (SH1093534.10FU) was significantly associated with inoculated plots (*p* = 0.042), and *Hebeloma hiemale* (SH1375983.10FU) with non-inoculated plots (*p* = 0.008; Table S20), but inoculation did not affect EcM alpha diversity (*p* = 0.861; Table S7).

**Figure 3.**
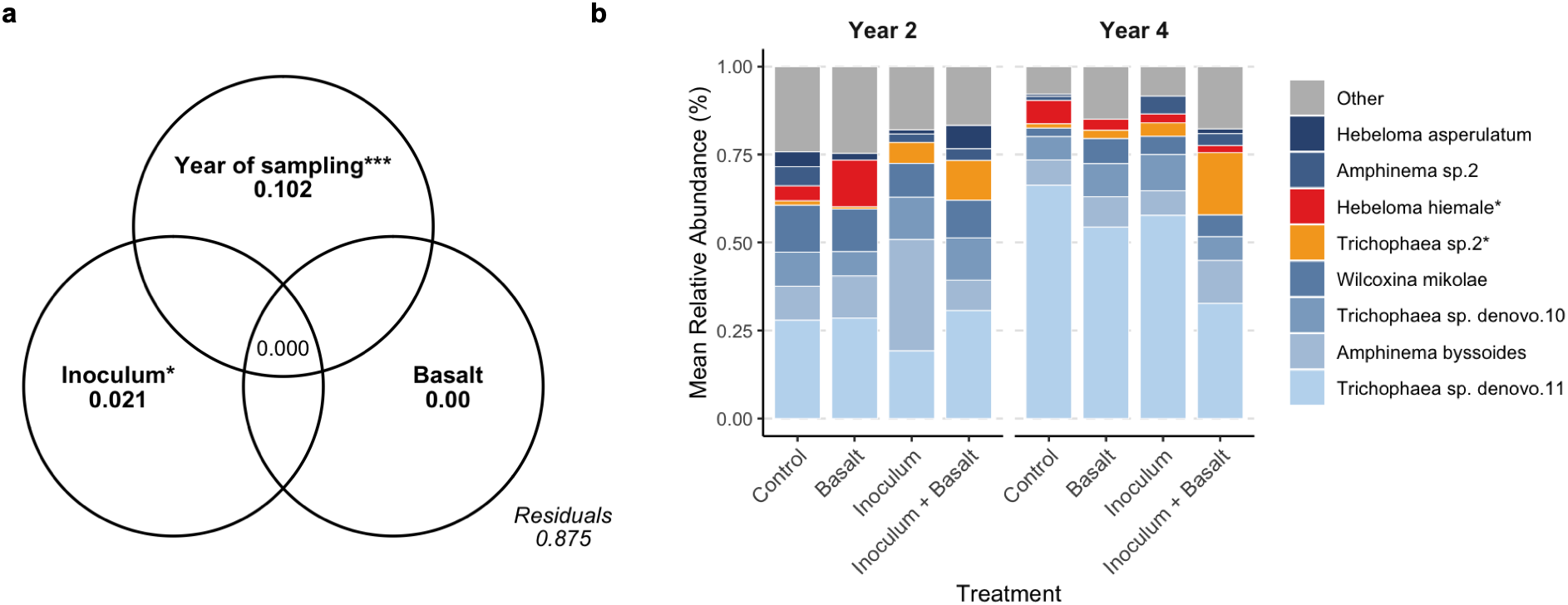
Ectomycorrhizal (EcM) fungal communities in Sitka spruce respond to soil inoculation (Inoculum). (a) Partitioning of variance in EcM fungal community composition explained by basalt, soil inoculation (Inoculum), and year of sampling. Values represent adjusted R^2^. Block effect was included in model but is not shown (R^2^ = 0.02). Asterisks indicate variables that significantly explain variation in community composition (distance-based RDA, *p*<0.05, n = 64). Values <0.001 are not shown. (b) Mean relative abundance of EcM taxa across treatments, panels separate sampling years. Bars show eight most abundant EcM taxa, and indicator species of inoculum are marked by an asterisk (*). Remaining taxa are grouped as “Other”.

**Figure 4.**
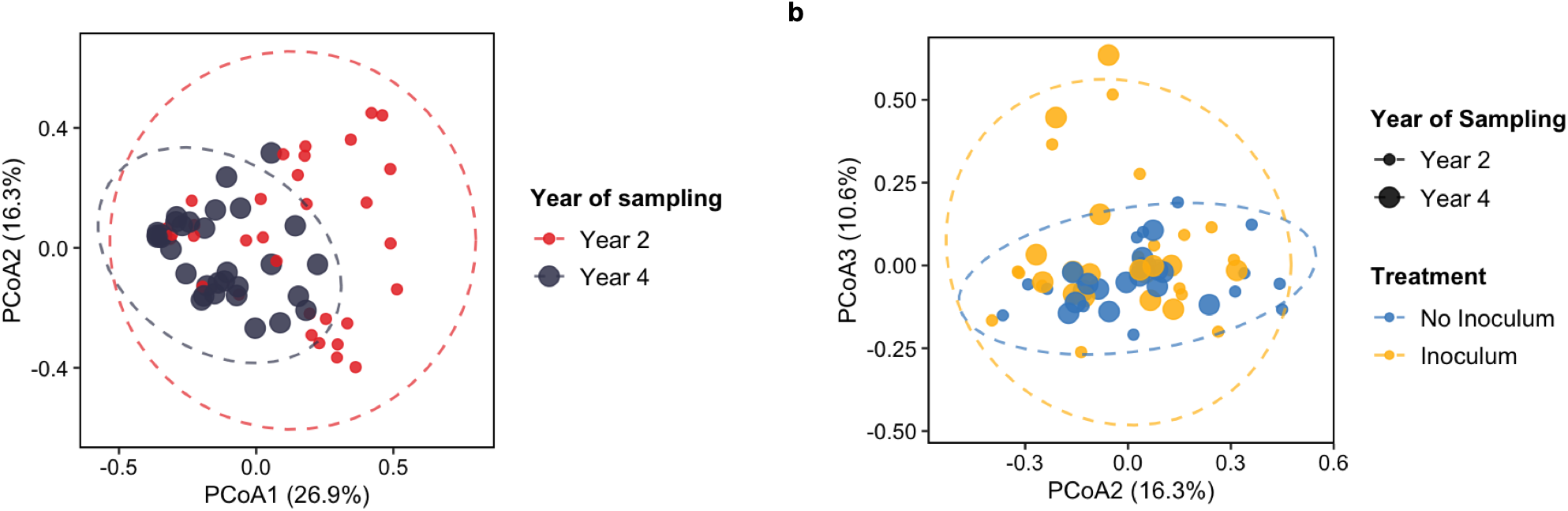
Ectomycorrhizal (EcM) fungal communities in Sitka spruce differ by sampling year and by soil inoculation (Inoculum). Principal coordinate analysis (PCoA) of EcM fungal communities based on Bray-Curtis dissimilarity (n = 64). (a) Axes 1 and 2 explain 26.9% and 16.3% of variation in community composition, respectively, and separate by year of sampling. Point size and colour indicate sampling year. (b) Axes 2 and 3 explain 16.3% and 10.6% of variation, respectively, and separate by inoculum treatment. Point size indicates sampling year, colour indicates inoculum treatment.

In the broadleaf plantation (Fig. 5), soil inoculation has no detectable effect on EcM or AM fungal community composition (PERMANOVA EcM, n = 62, p-F_1,58_ = 0.802, R^2^ = 0.013, *p* = 0.528; AM, n = 60, p-F_1,56_ = 1.291, R^2^ = 0.022, *p* = 0.168; Tables S11 & 13), on diversity (Table S8), or on AM colonisation (GLM *p =* 0.625; Fig. 7; Table S22).

**Figure 5.**
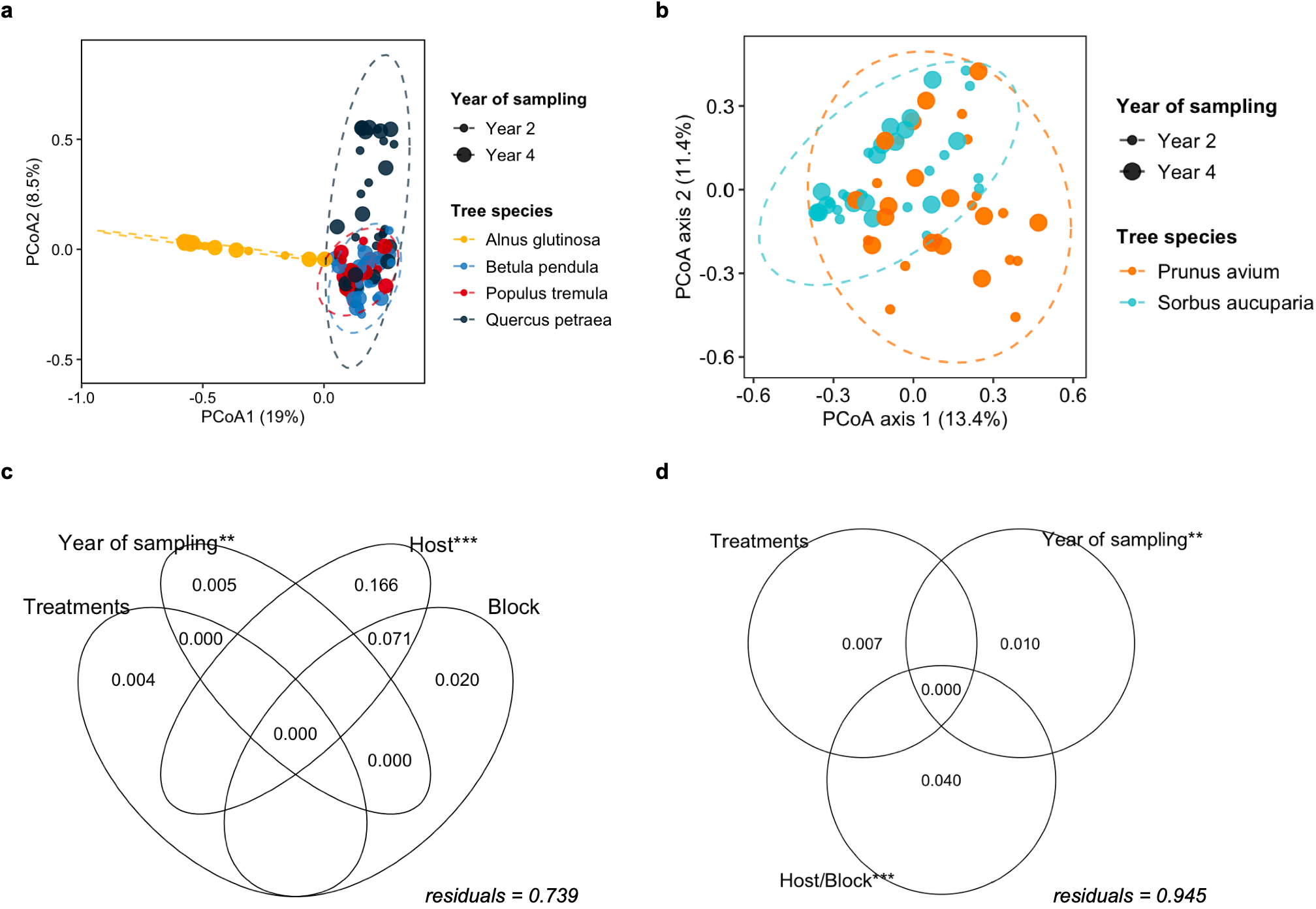
Mycorrhizal community composition in broadleaf plantation is structured by host tree species identity rather than by basalt or inoculum treatments. (a) Principal coordinate analysis (PCoA) of ectomycorrhizal (EcM) communities (of four broadleaf tree species) based on Bray-Curtis dissimilarity (n = 115). (b) PCoA of arbuscular mycorrhizal (AM) communities (of two broadleaf tree species) based on Jaccard dissimilarity (n= 60). (c,d) Variance partitioning of broadleaf EcM (c) and AM (d) community composition among basalt and soil inoculation (Treatments), year of sampling, and host tree species identity (Host), with experimental block included to account for spatial variation (Block). Because AM host species were sampled in non-overlapping block sets, host and block cannot be partitioned for AM communities and are shown as a combined fraction (Host/Block). Values are adjusted R². Asterisks indicate variables that significantly explain variation in community composition (distance-based RDA, p<0.05). Values <0.001 are not shown.

### 3.3 (H2) Tree nursery roots are the primary source of plantation EcM taxa, whereas AM fungi reflected pre-existing pasture

To identify the likely source of mycorrhizal colonisation in the tree plantations, root-associated EcM and AM communities were characterised from roots and bulk soils of potential sources of mycorrhizal colonisation.

For the root-associated EcM dataset, plantation EcM communities shared the greatest proportion of taxa with the tree nursery: 48.4% of broadleaf and 40.7% of spruce plantation EcM taxa also occurred on nursery seedling roots, compared with 2.2% and 9.3% for the corresponding woodland inoculum sources roots (Fig. 6a, c). Because taxon richness in the woodland inoculum sources was comparatively low, we also calculated overlap in the reverse direction: 15.4% of broadleaf and 41.7% of spruce inoculum taxa were detected on plantation roots. Unique OTUs found only in the plantation roots accounted for 33.3% of all OTUs in broadleaf plots, and 33.9% of all OTUs in spruce.

**Figure 6.**
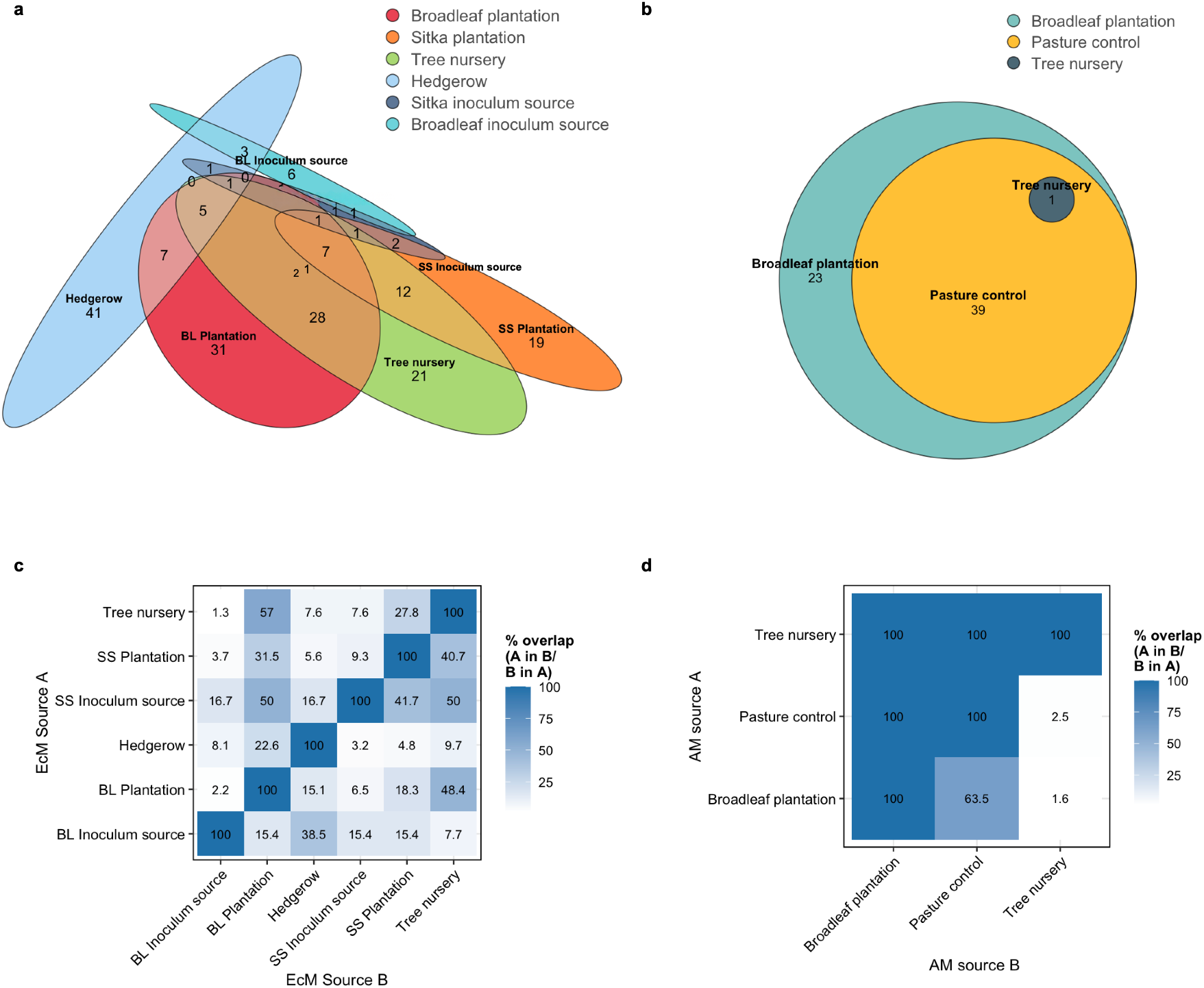
Shared mycorrhizal fungi between spruce and broadleaf plantations, and likely sources of mycorrhizal fungal taxa. Taxa were identified from sequenced colonised ectomycorrhizal (EcM) root tips. (a) Euler diagram showing unique and shared EcM taxa among broadleaf (BL) plantation, Sitka spruce (SS) plantations, the source tree nursery, surrounding hedgerows, and the woodland sites from which soil inoculant were derived (BL inoculum source and SS inoculum source). (b) Euler diagram showing unique and shared arbuscular mycorrhizal (AM) taxa among the BL plantation, tree nursery, and pasture control plots. (c, d) Heatmaps showing asymmetric pairwise comparisons between potential source communities and (c) BL or (d) SS plantations. Values represent the percentage of taxa in source A (rows) also found in source B (columns), and vice versa.

Overlap between soil and root-associated EcM fungal communities was consistently low (Fig. S5). No EcM fungi were shared between plantation roots and inoculum soils. Of the EcM fungi detected in the pasture control soils (a proxy for soil spore bank), 13.3% were also detected on roots of the broadleaf and spruce plantations. Overlap was slightly greater between plantation soils and tree roots, with 16.2% and 21.2% of EcM fungi from broadleaf and spruce plantation soils, respectively, also occurring on roots within the corresponding plantation.

For AM hosts, 63.5% of root associated fungi detected in the broadleaf plantation were also observed in the pasture control roots, and pasture communities were nested entirely within plantation communities (Fig. 6b, d). The single AM fungus detected on nursery roots also occurred in the planation. In contrast, no AM taxa from pasture control soils or inoculum were detected on plantation roots or soils, whereas 37.5% of AM taxa detected in the plantation soils also occurred on plantation roots (Fig. S6).

### 3.4 (H3) Limited AM and EcM fungal responses to basalt amendment

Basalt increased planation soil pH from 4.58 (4.23-4.97) to 5.21 (4.70-5.58; *p* = <0.001) (Fig. S1), but did not alter EcM or AM communities in either plantation type (PERMANOVA; spruce EcM, n = 64, p-F_1,60_= 0.850, R^2^ =0.012, *p* = 0.488; broadleaf EcM, n = 62, p-F_1,58_= 1.224, R^2^ =0.020, *p* = 0.141; broadleaf AM, n = 60, p-F_1,56_ = 1.155, R^2^ = 0.019, *p =* 0.251; Tables S10-13), alpha diversity (Tables S7-S9) or AM colonisation (GLM *p* = 0.636; Fig. 7). AM richness tended to be higher in the basalt-treated broadleaf plots (*p* = 0.076; Fig. S4), and two AM taxa, *Glomeraceae* sp. 1 (*p* = 0.008) and *Claroideoglomus* sp. 1 (*p* = 0.008) were associated with basalt-treated plots (Fig. 7).

**Figure 7.**
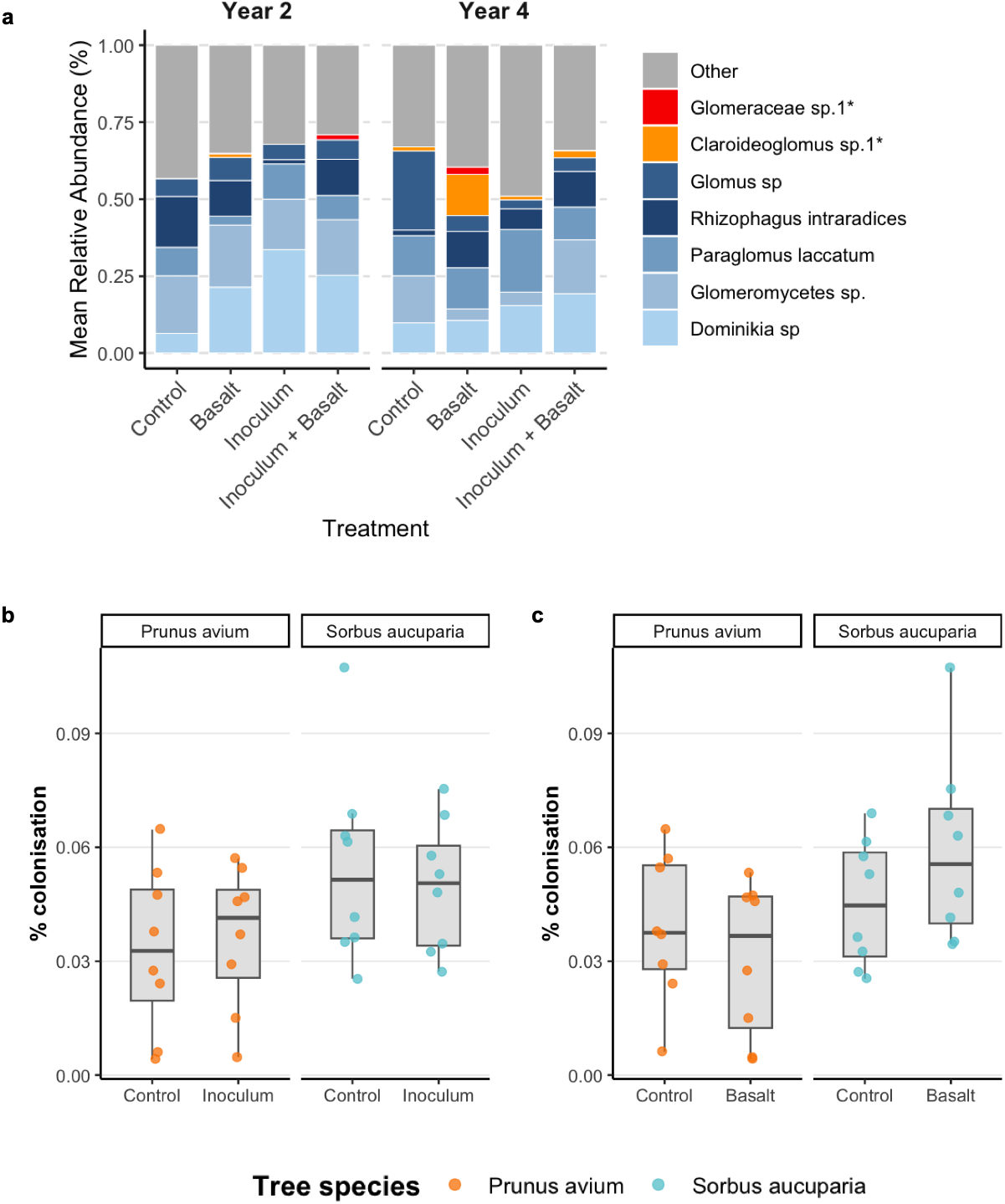
Arbuscular mycorrhizal (AM) fungal community response to basalt and inoculum treatments. (a) Mean relative abundance of AM taxa across treatments, panels separate sampling years. Bars show five most abundant AM taxa, and indicator species of basalt treatment are marked by an asterisk (*). Remaining taxa are grouped as “Other”. (b,c) Percentage of root colonised by AM fungi in cherry (*Prunus avium*) and rowan (*Sorbus aucuparia*) was not changed by either (b) inoculation (inoculum) or (c) basalt treatments. Colonisation included all aseptate fungal structures attributed to Glomeromycota, Mucoromycota (*Endogonales*), and unclassified aseptate fungi from roots collected in sampling year 2 only. Percentage colonisation was calculated for each seedling as the proportion of assessed root sampling units exhibiting AM colonisation relative to the total number of root sampling units examined (n = 32). Boxplots show the median, interquartile range (IQR), and whiskers extending to the minimum and maximum values; points represent individual trees.

### 3.5 (H1 & H3) Host identity determines EcM fungal community composition over treatment in broadleaf plantations

Host identity was the strongest predictor of EcM community composition when EcM fungal communities were aggregated by host rather at the plantation level (PERMANOVA; n = 115, p-F_3,108_ = 12.530, R^2^ = 0.250, *p* = 0.001), uniquely explaining 16.6% of variation (db-RDA; *p* = 0.001; Table S18), with a further 7.1% shared with block (Fig. 5c). Dispersion also differed among species (PERMDISP; *p* = 0.001), but species centroids were clearly separated in the ordination (Fig. 5a) and EcM fungal taxa significantly correlated with the plantations-wide PCoA ordination showed strong host specificity (Fig. 8a).

**Figure 8.**
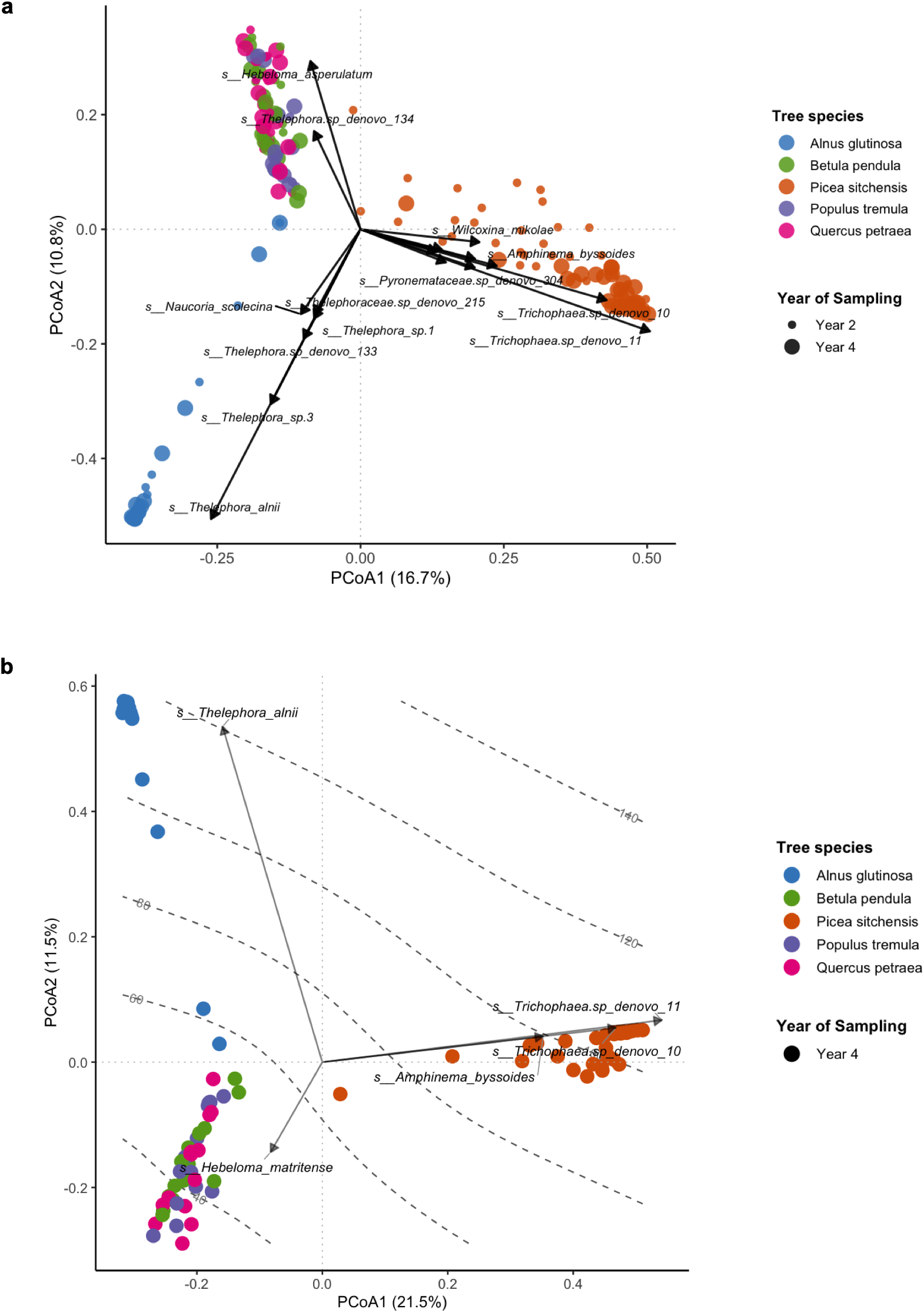
Principal coordinate analysis (PCoA) of ectomycorrhizal (EcM) fungal communities based on Bray-Curtis dissimilarity. (a) The first and second axes explain 16.7% and 10.8% of variation in community composition, respectively. Arrows represent EcM fungi significantly (p < 0.01) correlated with the first two PCoA axes (envfit, 999 permutations). Length of arrow represents strength of correlation. Points are coloured by host tree species identity. (b) PCoA of EcM fungal communities in sampling year 4, to visualize EcM taxon associated with differences in biomass among tree hosts. Grey contour lines show projected tree biomass (kg) across ordination space, fitted using a generalised additive model (GAM; edf = 3.486, adj. R^2^= 0.40, deviance explained = 42.3%, *p* < 0.001) with biomass as a smooth function of the PCoA1 and PCoA2. Arrows indicate associations between individual EcM taxa and biomass, identified using binomial generalised linear models (GLMs) and projected onto the ordination (envfit, 999 permutations, p<0.05).

Host identity was also a significant predictor of AM root colonisation (GLM; n= 32, β = 0.520, *p* = 0.006), whereas the combined host/block fraction explained only 4.0% of AM community variation (db-RDA; *p* = 0.001; Table S19), less than the equivalent fraction for EcM communities (Fig. 5). Cherry and rowan were sampled in non-overlapping blocks, so host and block fractions cannot be separated.

Given the strong influence of host identity on broadleaf EcM communities (Fig. 5), we tested for treatment effects while accounting for host identity. When host identity was included as a random effect, neither inoculation nor basalt affected EcM community composition along PCoA axes 1-3 (Fig. S3; Table S23). Analyses conducted separately for each tree species, also detected no significant treatment effects (analysis of host-specific datasets is not shown).

### 3.6 (H4) Tree species, ectomycorrhizas, treatments and plantation biomass

We modelled square-root transformed tree biomass for each sampling year, using PCoA scored from the plantations-wide EcM ordination (Fig. S7) to assess the relative importance of EcM community composition, host tree species identity, and treatments. Plantation type was the best predictor in both years, with broadleaf plots exhibiting more biomass than spruce (year 2, β = 2.945, *p* = <0.001; year 4, β = 8.120, *p* = <0.001). In year 4, biomass increased with PCoA2 (β = 2.804, *p* = 0.016), whereas PCoA1 showed a marginal negative effect (β = −7.001, *p* = 0.059); neither axis was significant in year 2 (Table S24).

Next, because EcM communities were strongly structured by host identity, we separated the contribution of within-species EcM community composition from among-host variation in year 4 by centering PCoA scores for each tree species. The tree species identity model was far better supported than the (uncentred) EcM model (raw PCoA scores; AIC 320 vs 437; Table S25). There was weak evidence that within-host EcM composition contributed additional information once tree species was accounted for (anova, X² = 5.55, df = 2, *p* = 0.062; AIC 319 vs 320; Table S26), although within-species PCoA2 was positively associated with biomass (β = 2.953, *p* = 0.048). Treatments and their interaction remained non-significant.

We next asked which EcM taxa best characterise biomass differences among the tree species. A two-dimensional GAM fitted to year 4 showed that biomass varied significantly across the EcM ordination (GAM; deviance explained = 42.3%, *p* < 0.001; Fig. 8b), reflecting host-structured differences. Taxa associated with higher biomass (kg; binomial GLM; Table S27) included *Trichophaea* sp. *de novo* 10 (*p* adj. = 0.003), Trichophaea sp. *de novo* 11 (*p* adj. = 0.003) and *Amphinema byssoides* (*p* adj. = 0.036) in spruce, and *Thelephora alnii* in alder (*p* adj. = 0.036), where *Hebeloma matritense was* associated with lower biomass in oak, aspen and birch (*p* adj. = 0.041).

## 4 Discussion

Here, we report results from a full factorial field experiment testing how ERW (through basalt amendment) and soil inoculation affect root-associated mycorrhizas during early woodland establishment. To the best of our knowledge, this study represents the largest and most highly replicated field study examining the impacts of these CDR strategies on mycorrhizal communities.

### 4.1 Host-dependent mycorrhizal responses to soil inoculation limited by established fungi

Soil inoculation resulted in subtle changes to spruce EcM fungal community composition (Fig. 3; Fig.4) but had no detectable effect on broadleaf mycorrhizas (H1) (Fig. 5). Because plantation EcM fungi derived mainly from the nursery rather than inoculum (H2) (Fig. 6), these compositional shifts in spruce are likely to reflect changes in the relative abundance of resident fungi rather than the introduction of new inoculum-derived taxa.

Inoculation may instead have altered ecological filters regulating mycorrhizas, such as competitive interactions, as indicator species analysis suggests. *Trichophaea* sp. 2 was positively associated with inoculum treatment (Fig. 3) but was detected only in tree nursery and plantation root-associated fungal communities, not in inoculum or plantation soils (S5). Species within this genus are common pioneers in tree nurseries and conifer woodlands and produce resistant propagules that facilitate rapid root colonisation in disturbed environments (Gautier et al., 2026; Kipfer et al., 2011; Rosenstock et al., 2018; Tedersoo et al., 2006). Consistent with this strategy, increases in *Trichophaea* abundance following soil inoculation have been also reported in *Pseudotsuga sinensis* (Wen et al., 2018), *Picea asperata* and *Abies fargesii* var. *faxoniana* (Wang et al., 2023). Thus, inoculation appears to favour nursery-derived *Trichophaea* rather than introduce it.

Inoculum may also have altered host-fungal nutrient exchange. Soil inoculation can increase nitrogen availability (Lance et al., 2019), potentially reducing host-investment in EcM fungi with better nutrient-mobilising strategies (Johnson et al., 2013). *Hebeloma hiemale* was negatively associated with inoculated plots and species within *Hebeloma*, among others, have been found to possess enzymatic capacities for soil nitrogen mobilisation, which may incur relatively high carbon costs on its host (Doré et al., 2015; Herinckx et al., 2026; Pena et al., 2023; Tibbett et al., 1999; Zhou et al., 2022). Notably, distinguishing biotic from abiotic inoculum effects (McMahen et al., 2022) would require sterile controls, although sterilisation itself alters soil chemistry (Berns et al., 2008; Endlweber & Scheu, 2006; King et al., 2024).

It is conceivable that the absence of inoculum effects on broadleaf EcM community composition reflected strong host structuring within the mixed-species broadleaf plantation, which masked small, divergent responses among broadleaf tree species (Fig. 5). These findings are consistent with the well-established role of host identity in shaping EcM community assembly (Ferlian et al., 2021; Martínez-García et al., 2015; Tedersoo et al., 2003; van der Linde et al., 2018). Thus, inoculation effects on broadleaf species, if present, may be confined to host-specific taxa and require greater within-species sampling efforts than in this study. These results add to evidence that inoculation outcomes are context- and host-dependent (Dickie et al., 2007; Grove et al., 2019; Lance et al., 2019; McMahen et al., 2022; Parasquive et al., 2024; St-Denis et al., 2017; Taniguchi et al., 2021) and emphasise the need to consider the utility of CDR strategies in mixed vs monoculture forestry.

The mycorrhizal responses reported here are concordant with the biomass responses reported for the same field trial by Waring et al. (2026), where individual tree-level growth responses to soil inoculation were strongest for spruce. Our results show that the same treatment also altered the composition of root-associated mycorrhizas in spruce, extending associations inferred from bulk soil to the symbiotic partners directly mediating host nutrient uptake. This potential functional link is consistent with evidence that shifts in EcM community composition can alter soil carbon cycling dynamics, nutrient-acquisition strategies and functional trait distributions, with consequences for host nutrition and growth even where overall fungal diversity remains unchanged (Kaiser & Anthony, 2025; Pellitier et al., 2021; Yan et al., 2025), although here this link remains correlative.

Finally, nursery-derived fungi greatly contribute to plantation mycorrhizal communities, which may limit the establishment of inoculated taxa. This observed persistence of nursery-associated EcM fungi indicates that the nursery stage represents a more effective point of intervention than inoculation during planting (Fig. 6). Previous studies have also shown that nursery-derived EcM fungi can contribute substantially to mature woodland communities (Flykt et al., 2008; Palfner et al., 2005) and that nursery-applied commercial or single-strain inoculant can persist after out-planting (Franco et al., 2014; Quoreshi et al., 2008). However, evidence assessing the implementation of soil inoculation at the nursery stage is limited (Durgapal et al., 2002; St-Denis et al., 2017; Taniguchi et al., 2021). Conversely, as hypothesised, AM fungal communities appeared to be assembled largely from taxa associated with roots of the previous pasture (Fig. 6).

### 4.2 Limited mycorrhizal responses to basalt across plantations

We report no detectable effects of basalt treatment on EcM or AM community composition, mycorrhizal diversity or AM colonisation (H3), despite a significant increase in soil pH (Fig. S1). This was unexpected because soil pH is a key determinant of both EcM and AM fungal communities (Davison et al., 2021; Kluber et al., 2012; Suz et al., 2014). One explanation for the pattern observed is that the shift in soil pH was large enough to alter total soil fungal communities, as reported at the same site by Waring et al. (2026), but not root-associated mycorrhizal fungi, which may be buffered against environmental change by host filtering (Goldmann et al., 2016).

These contrasting responses may also reflect soil eDNA being an incomplete proxy for root-associated communities, as we found only 16.2% and 21.2% of plantation soil EcM taxa also detected on roots in broadleaf and conifer plantations, respectively (Figs. S5 & S6). Overall, our results indicate established, root-associated mycorrhizas are unchanged under modest, basalt induced changes to soil chemistry in young plantations.

### 4.3 Ectomycorrhizal communities, tree species identity and carbon capture

Host identity was the dominant predictor of plantation biomass, consistent with the established role of tree species identity in determining both above- and belowground carbon sequestration (Mc Kay et al., 2021; Waring et al., 2022). Within-host tree species variation in EcM community composition was positively associated with biomass, but the effect was small and its addition did not significantly improve model support (Table S26).

Several EcM taxa that were strongly host-structured and were associated with biomass differences among tree species (Fig. 8). *Thelephora alnii* was associated with greater biomass in alder, and *Trichophaea spp.* and *Amphinema byssoides* with greater biomass in spruce, whereas *Hebeloma matritense* was associated with lower biomass across broadleaf species. These associations are consistent with previously reported links between fast-growing broadleaf stands and EcM fungi within the Thelephoraceae, and between fast-growing coniferous stands and EcM fungi within the Atheliaceae (Anthony et al., 2022).

Collectively, our data emphasise that fungal community composition and host identity are intertwined components of early plantation performance, with host identity by far the strongest predictor, likely because host traits also govern litter chemistry, nutrient demand, carbon input to rooting system and root architecture, all of which contribute to carbon cycling (Melvin et al., 2015). Resolving their independent contributions and thereby establishing whether inoculation could be optimised to introduce host-specific taxa that enhance growth, will require fine-scale within-host studies under both field and controlled conditions.

### 4.4 Limitations

Sampling year explained more variation in mycorrhizal community composition than treatments (Figs 3 & 5), consistent with previously observed increasing homogeneity of fungal communities as host mature (Zhao et al., 2025), also observed in our study (Fig. 4). Because the strongest treatment effects on tree biomass occurred during the first growing season (Waring et al., 2026), our ability to detect corresponding EcM or AM responses may have been limited by sampling only in the second and fourth growing seasons, by which time inoculation or basalt effects on establishing seedlings may already have attenuated due to these temporal dynamics.

The relatively reduced sampling effort of inoculum-source roots and soils, compared to plantation, may have underestimated the contribution of individual taxa to plantation communities. The apparent overlap between root and soil communities was also likely limited by differences in DNA sequencing approaches, primer sets and bioinformatic pipelines, all of which influence taxon detection and classification. Because sampling was designed at the plantation level, our sampling effort was reduced for individual broadleaf hosts compared to spruce alone. This limited power to detect responses within broadleaf species and confounded our ability to distinguish between host and block effects in broadleaf AM fungal communities.

The quantity and source of soil inoculum may have also influenced the extent to which mycorrhizas respond to inoculation; the difference in species-mixtures between the broadleaf plantation and the broadleaf woodland inoculum source (dominated by oak and beech) may limit the suitability of inoculum taxa, especially due to the strong host specific effect observed even at this early plantation stage.

### 4.5 Conclusions

Early mycorrhizal responses to CDR interventions during afforestation on former pasture were strongly dependent on host identity and pre-existing fungal source pools. Soil inoculation produced only a small shift in spruce EcM only, and basalt application did not substantially disrupt mycorrhizas despite altering soil chemistry, suggesting ERW is unlikely to negatively affect mycorrhizal communities during woodland establishment.

The tree nursery was the main identifiable source of plantation EcM fungi, while AM communities reflected the previous pasture, suggesting that interventions applied after outplanting may be constrained by fungi already established.

For the optimisation of CDR strategies in early afforestation, future work should test approaches tailored to host-identity, existing fungal source pools, and earlier intervention timing to generate more persistent changes in mycorrhizal communities and, ultimately, tree performance.

## Acknowledgements

The authors wish to thank Jess Bode, Elena Arrigoni, Guillaume Delhaye, Robert Barber, Rose Gooda, Angela Biro, Saskia Pearce, Tom Weeks, Kim Arroyo, and all volunteers including Elizabeth Brogan, Annette Rajnak, Freya Jessop, Joshua Dickerson, Khushi Joshi, Leo He, Sean Barry, Sebastian Dohn, Victor Carreras, Zosia Wardynska. This project was partly funded by The Carbon Community and generously supported by a grant from the De Laszlo Foundation.

## Author contributions

Laura M.Suz, Martin Bidartondo, Oliver Linsay and Isabelle Thomson conceived the ideas and designed methodology; Isabelle Thomson, Oliver Lindsay, Laura M.Suz, Martin Bidartondo, Bonnie Waring collected the data; Isabelle Thomson analysed the data; Isabelle Thomson led the writing of the manuscript. All authors contributed critically to the drafts and gave final approval for publication.

## Inclusion statement

Our international author team, including scientists from the host country, was involved in the research design from the start to ensure diverse perspectives.

## 10. Figures

**Figure 9.**
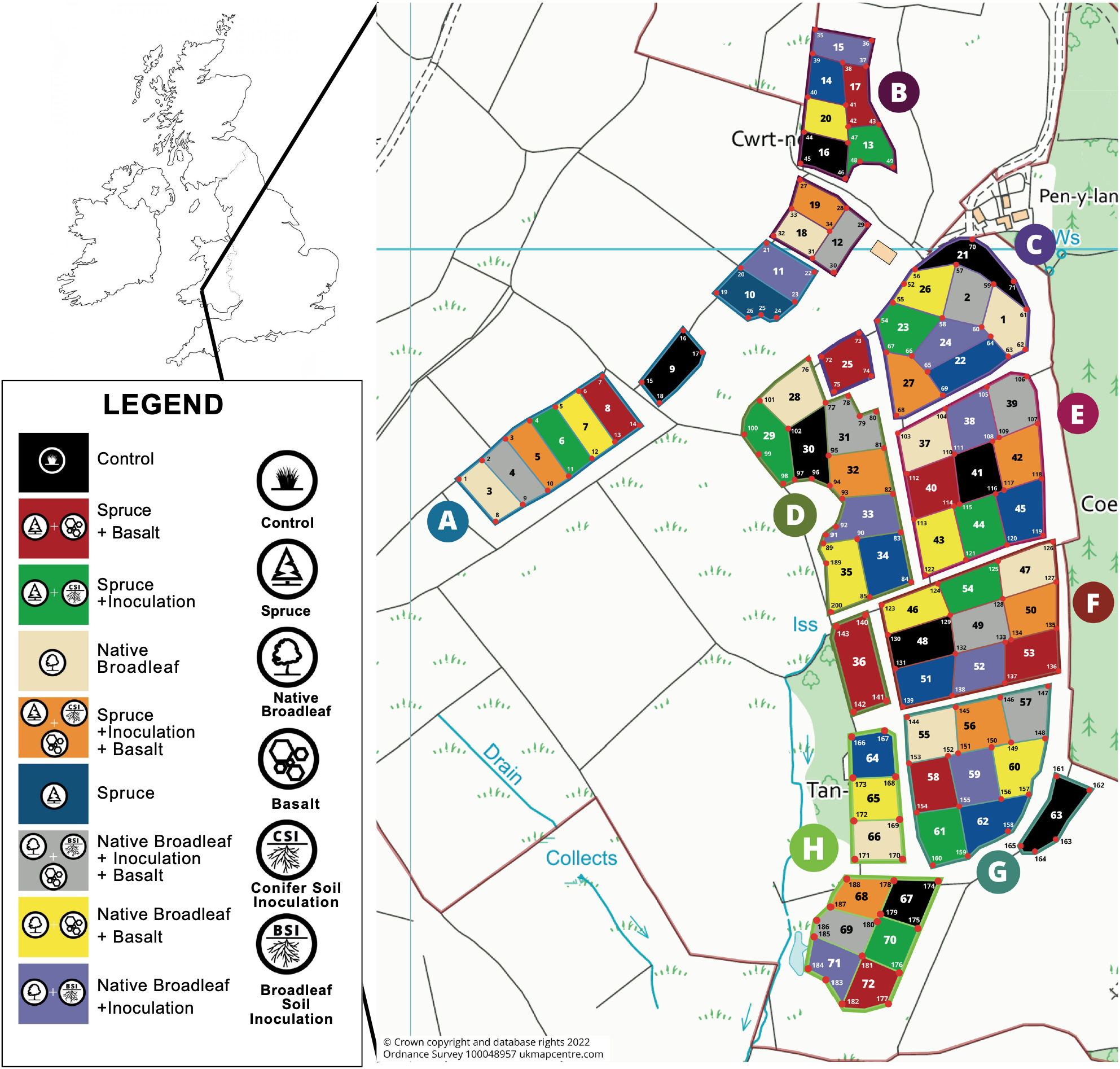
Map of The Carbon Community project experimental field trial, Glandwr Forest, Cynghordy, Wales. This includes the 72 plots and their corresponding treatment (s) (11.52 ha; ∼25,000 trees). Letters indicate blocks, and numbers indicate plot. Adapted from Waring et al., (2026).

**Figure 10.**
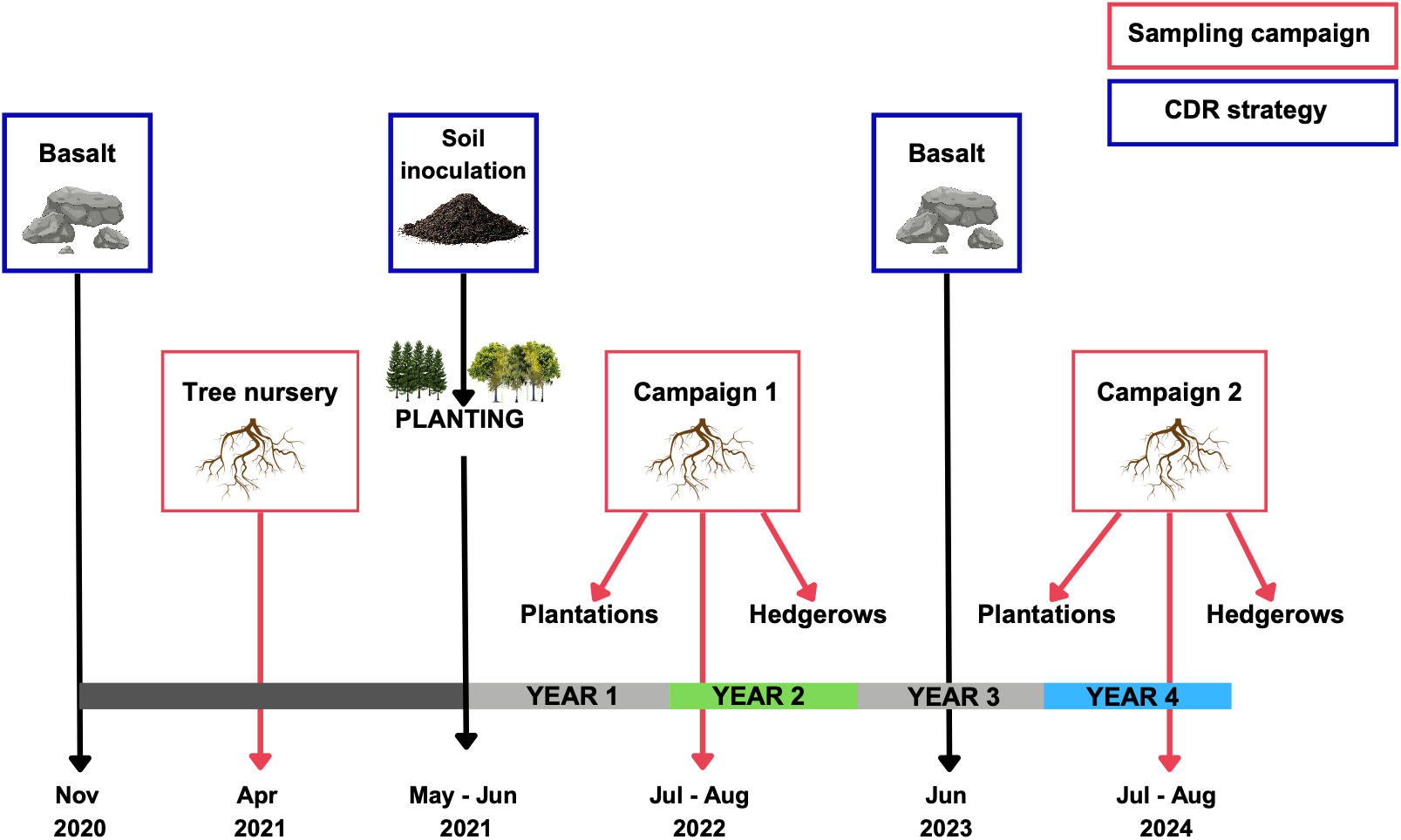
Experimental timeline for The Carbon Community project, illustrating the timing of treatment applications (dark blue) and sampling campaigns (red) throughout the study. Root-associated fungal communities were sampled from the tree nursery, plantation plots, and adjacent hedgerows.

**Figure 11.**
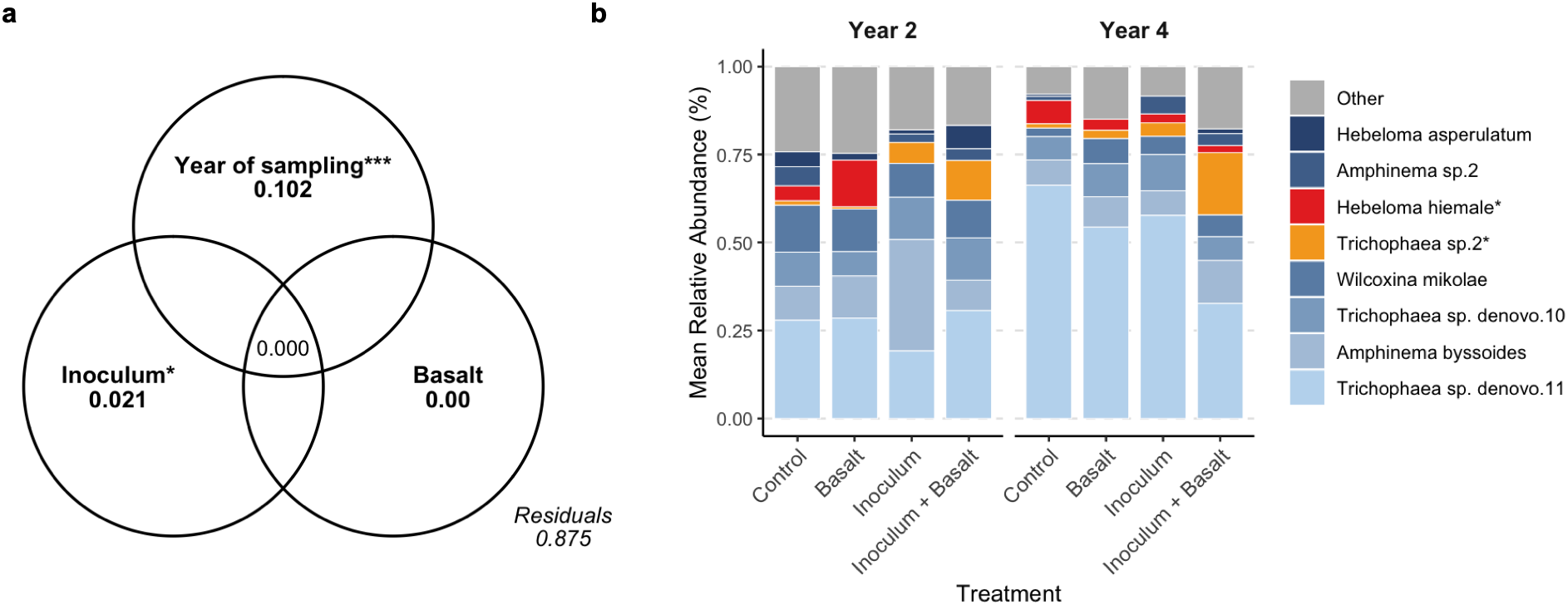
Ectomycorrhizal (EcM) fungal communities in Sitka spruce respond to soil inoculation (Inoculum). (a) Partitioning of variance in EcM fungal community composition explained by basalt, soil inoculation (Inoculum), and year of sampling. Values represent adjusted R^2^. Block effect was included in model but is not shown (R^2^ = 0.02). Asterisks indicate variables that significantly explain variation in community composition (distance-based RDA, *p*<0.05, n = 64). Values <0.001 are not shown. (b) Mean relative abundance of EcM taxa across treatments, panels separate sampling years. Bars show eight most abundant EcM taxa, and indicator species of inoculum are marked by an asterisk (*). Remaining taxa are grouped as “Other”.

**Figure 12.**
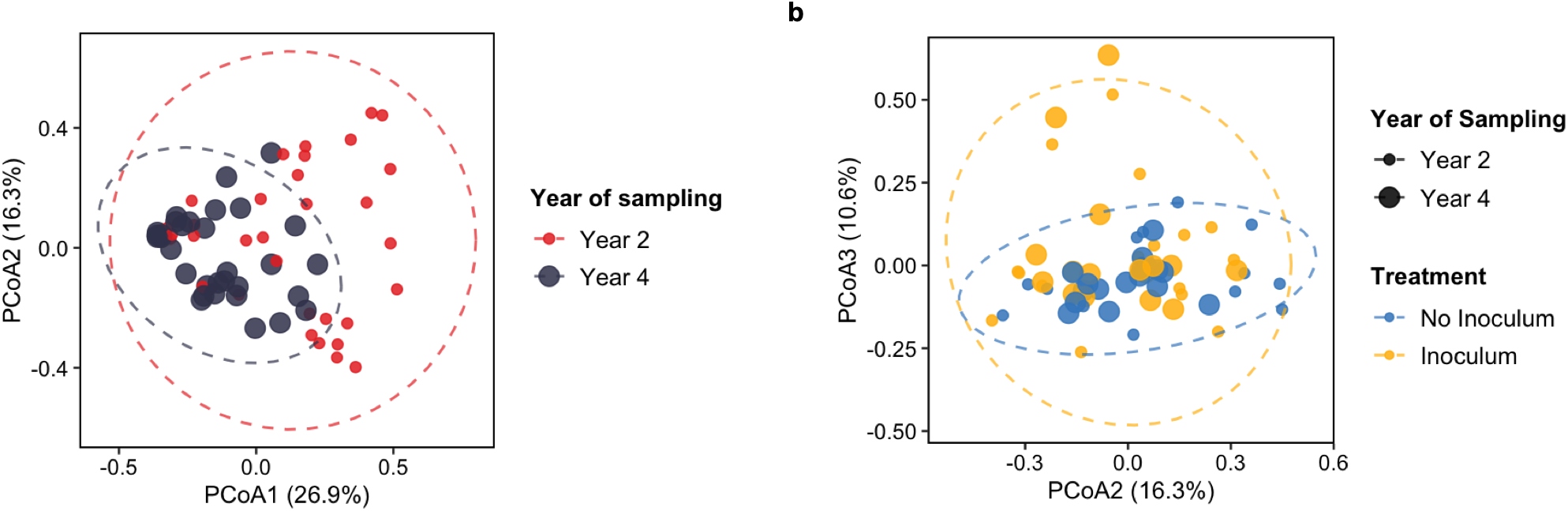
Ectomycorrhizal (EcM) fungal communities in Sitka spruce differ by sampling year and by soil inoculation (Inoculum). Principal coordinate analysis (PCoA) of EcM fungal communities based on Bray-Curtis dissimilarity (n = 64). (a) Axes 1 and 2 explain 26.9% and 16.3% of variation in community composition, respectively, and separate by year of sampling. Point size and colour indicate sampling year. (b) Axes 2 and 3 explain 16.3% and 10.6% of variation, respectively, and separate by inoculum treatment. Point size indicates sampling year, colour indicates inoculum treatment.

**Figure 13.**
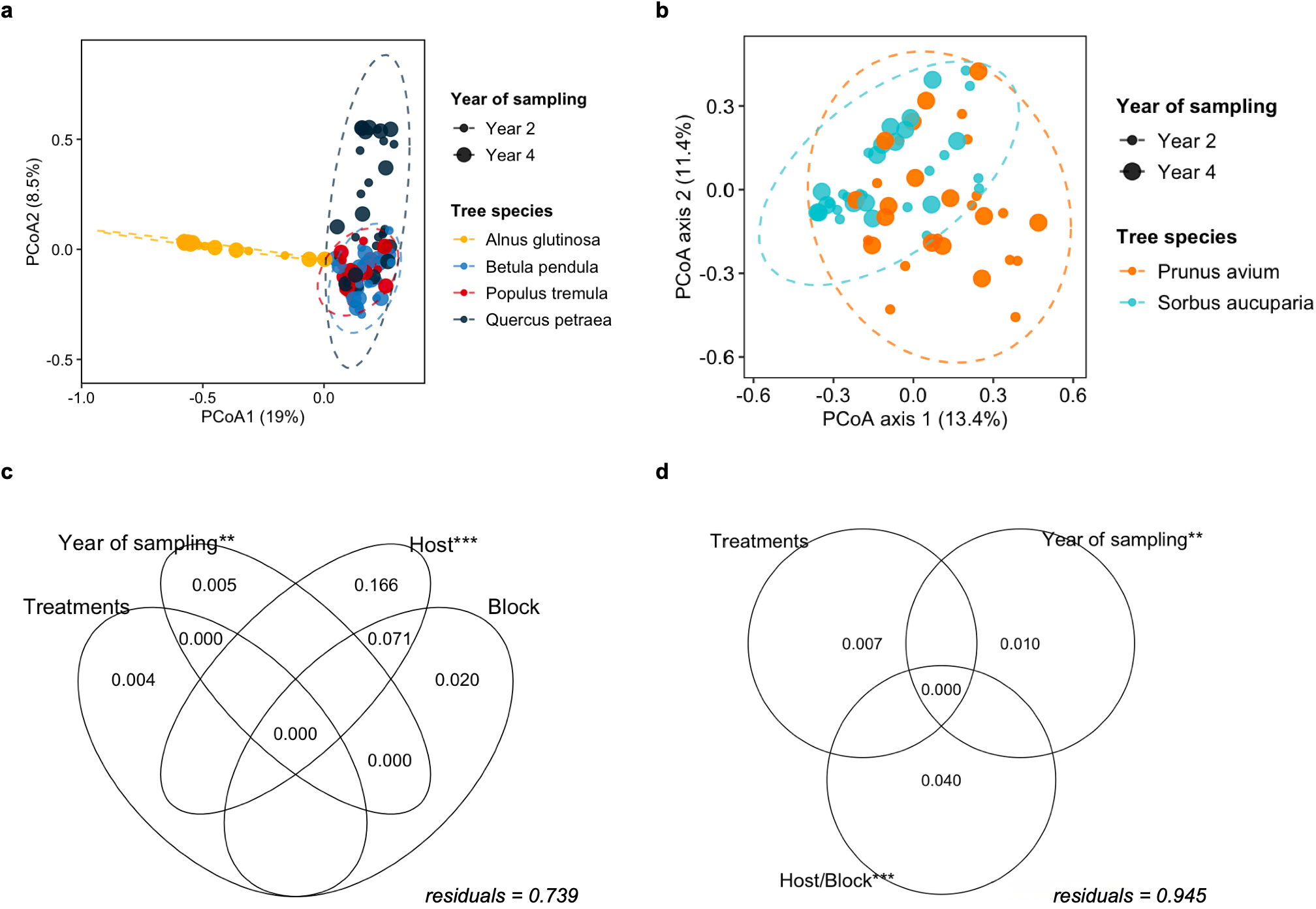
Mycorrhizal community composition in broadleaf plantation is structured by host tree species identity rather than by basalt or inoculum treatments. (a) Principal coordinate analysis (PCoA) of ectomycorrhizal (EcM) communities (of four broadleaf tree species) based on Bray-Curtis dissimilarity (n = 115). (b) PCoA of arbuscular mycorrhizal (AM) communities (of two broadleaf tree species) based on Jaccard dissimilarity (n= 60). (c,d) Variance partitioning of broadleaf EcM (c) and AM (d) community composition among basalt and soil inoculation (Treatments), year of sampling, and host tree species identity (Host), with experimental block included to account for spatial variation (Block). Because AM host species were sampled in non-overlapping block sets, host and block cannot be partitioned for AM communities and are shown as a combined fraction (Host/Block). Values are adjusted R². Asterisks indicate variables that significantly explain variation in community composition (distance-based RDA, p<0.05). Values <0.001 are not shown.

**Figure 14.**
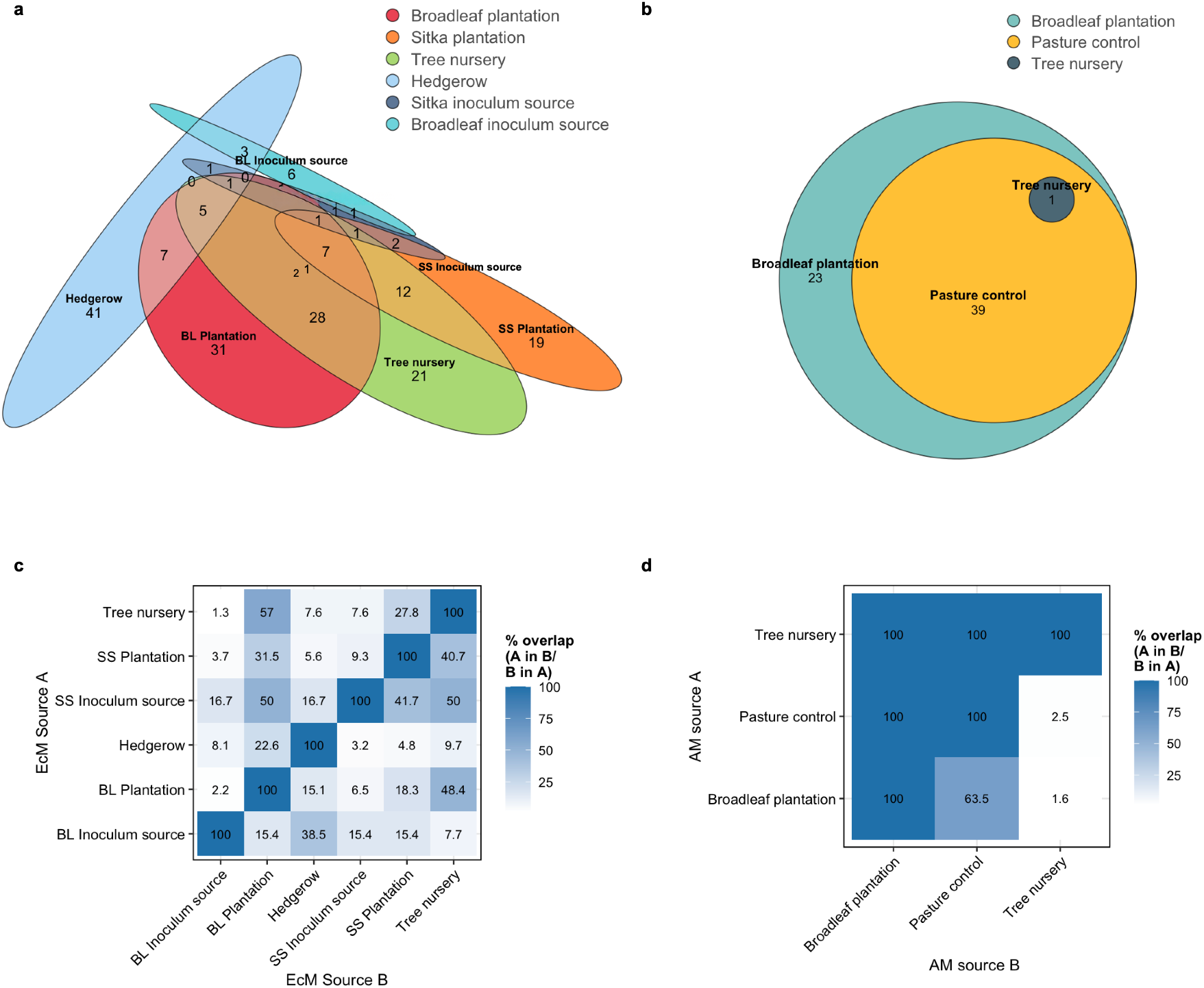
Shared mycorrhizal fungi between spruce and broadleaf plantations, and likely sources of mycorrhizal fungal taxa. Taxa were identified from sequenced colonised ectomycorrhizal (EcM) root tips. (a) Euler diagram showing unique and shared EcM taxa among broadleaf (BL) plantation, Sitka spruce (SS) plantations, the source tree nursery, surrounding hedgerows, and the woodland sites from which soil inoculant were derived (BL inoculum source and SS inoculum source). (b) Euler diagram showing unique and shared arbuscular mycorrhizal (AM) taxa among the BL plantation, tree nursery, and pasture control plots. (c, d) Heatmaps showing asymmetric pairwise comparisons between potential source communities and (c) BL or (d) SS plantations. Values represent the percentage of taxa in source A (rows) also found in source B (columns), and vice versa.

**Figure 15.**
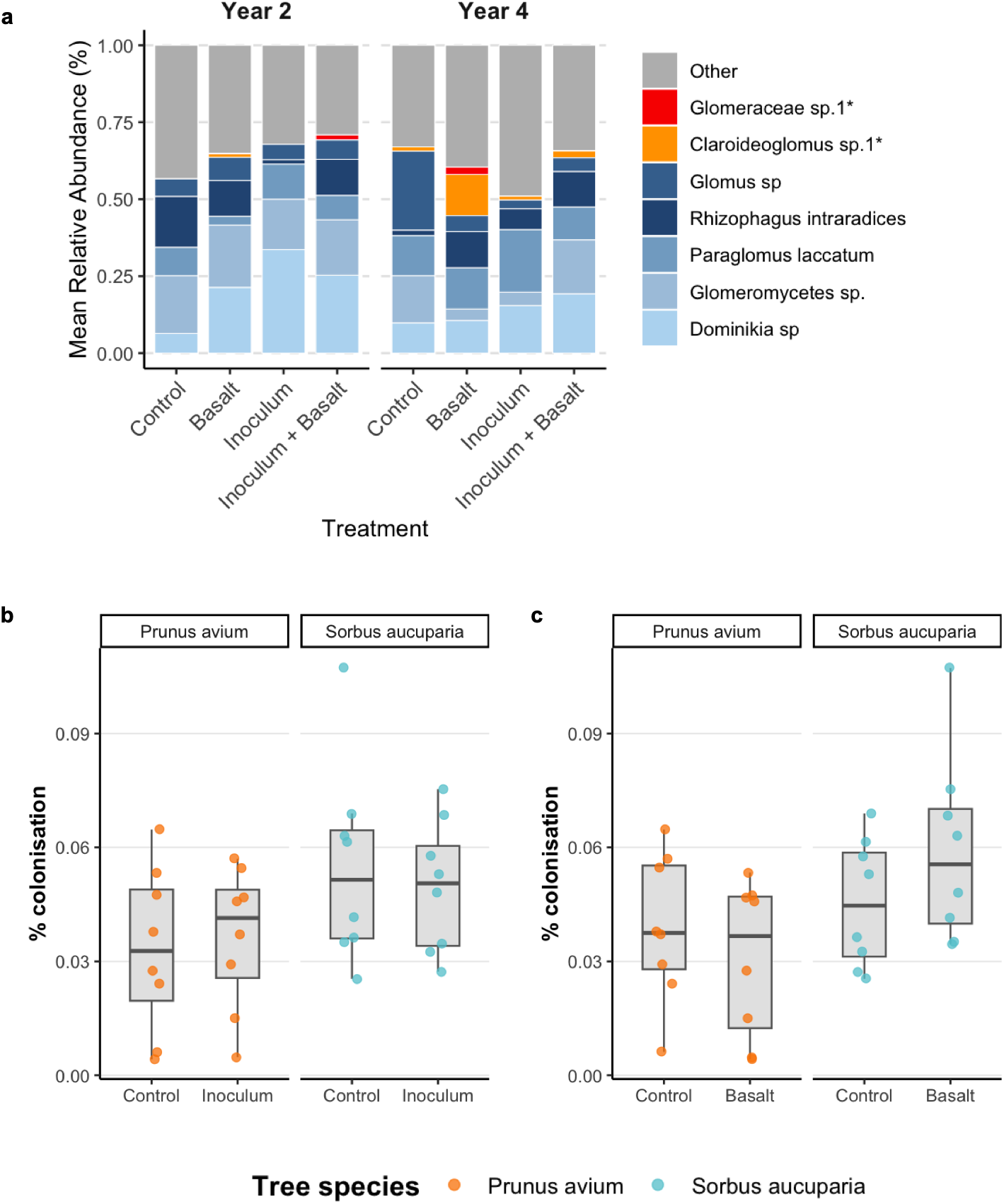
Arbuscular mycorrhizal (AM) fungal community response to basalt and inoculum treatments. (a) Mean relative abundance of AM taxa across treatments, panels separate sampling years. Bars show five most abundant AM taxa, and indicator species of basalt treatment are marked by an asterisk (*). Remaining taxa are grouped as “Other”. (b,c) Percentage of root colonised by AM fungi in cherry (*Prunus avium*) and rowan (*Sorbus aucuparia*) was not changed by either (b) inoculation (inoculum) or (c) basalt treatments. Colonisation included all aseptate fungal structures attributed to Glomeromycota, Mucoromycota (*Endogonales*), and unclassified aseptate fungi from roots collected in sampling year 2 only. Percentage colonisation was calculated for each seedling as the proportion of assessed root sampling units exhibiting AM colonisation relative to the total number of root sampling units examined (n = 32). Boxplots show the median, interquartile range (IQR), and whiskers extending to the minimum and maximum values; points represent individual trees.

**Figure 16.**
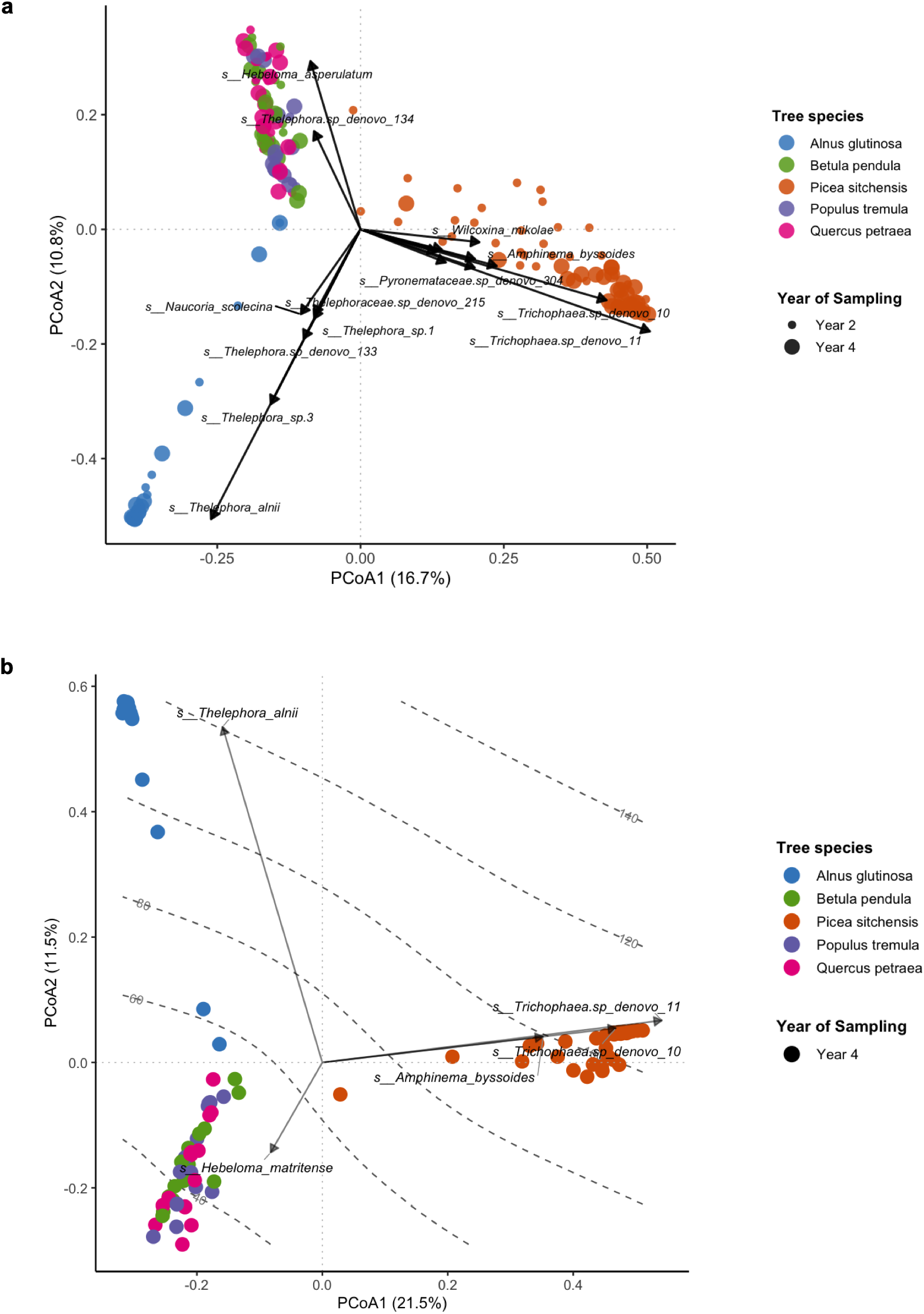
Principal coordinate analysis (PCoA) of ectomycorrhizal (EcM) fungal communities based on Bray-Curtis dissimilarity. (a) The first and second axes explain 16.7% and 10.8% of variation in community composition, respectively. Arrows represent EcM fungi significantly (p < 0.01) correlated with the first two PCoA axes (envfit, 999 permutations). Length of arrow represents strength of correlation. Points are coloured by host tree species identity. (b) PCoA of EcM fungal communities in sampling year 4, to visualize EcM taxon associated with differences in biomass among tree hosts. Grey contour lines show projected tree biomass (kg) across ordination space, fitted using a generalised additive model (GAM; edf = 3.486, adj. R^2^= 0.40, deviance explained = 42.3%, *p* < 0.001) with biomass as a smooth function of the PCoA1 and PCoA2. Arrows indicate associations between individual EcM taxa and biomass, identified using binomial generalised linear models (GLMs) and projected onto the ordination (envfit, 999 permutations, p<0.05).

## References

Abarenkov, K., Nilsson, R. H., Larsson, K. H., Taylor, A. F. S., May, T. W., Frøslev, T. G., Pawlowska, J., Lindahl, B., Põldmaa, K., Truong, C., Vu, D., Hosoya, T., Niskanen, T., Piirmann, T., Ivanov, F., Zirk, A., Peterson, M., Cheeke, T. E., Ishigami, Y., … Kõljalg, U. (2024). The UNITE database for molecular identification and taxonomic communication of fungi and other eukaryotes: sequences, taxa and classifications reconsidered. Nucleic Acids Research, 52(D1), D791–d797. 10.1093/nar/gkad1039

Anda, M., Shamshuddin, J., & Fauziah, C. I. (2015). Improving chemical properties of a highly weathered soil using finely ground basalt rocks. CATENA, 124, 147–161. 10.1016/j.catena.2014.09.012

Anderson, M. J., & Walsh, D. C. (2013). PERMANOVA, ANOSIM, and the Mantel test in the face of heterogeneous dispersions: what null hypothesis are you testing? Ecological Monographs, 83(4), 557–574.

Anthony, M. A. (2025). Does ectomycorrhizal fungal biodiversity affect tree growth? Fungal Ecology, 74, 101413. 10.1016/j.funeco.2025.101413

Anthony, M. A., Crowther, T. W., van der Linde, S., Suz, L. M., Bidartondo, M. I., Cox, F., Schaub, M., Rautio, P., Ferretti, M., Vesterdal, L., De Vos, B., Dettwiler, M., Eickenscheidt, N., Schmitz, A., Meesenburg, H., Andreae, H., Jacob, F., Dietrich, H.-P., Waldner, P., … Averill, C. (2022). Forest tree growth is linked to mycorrhizal fungal composition and function across Europe. The ISME Journal, 16(5), 1327–1336. 10.1038/s41396-021-01159-7

Anthony, M. A., Tedersoo, L., De Vos, B., Croisé, L., Meesenburg, H., Wagner, M., Andreae, H., Jacob, F., Lech, P., Kowalska, A., Greve, M., Popova, G., Frey, B., Gessler, A., Schaub, M., Ferretti, M., Waldner, P., Calatayud, V., Canullo, R., … Averill, C. (2024). Fungal community composition predicts forest carbon storage at a continental scale. Nature Communications, 15(1), 2385. 10.1038/s41467-024-46792-w

Augé, R. M., Toler, H. D., & Saxton, A. M. (2014). Arbuscular mycorrhizal symbiosis and osmotic adjustment in response to NaCl stress: a meta-analysis. Frontiers in Plant Science, 5, 562. 10.3389/fpls.2014.00562

Averill, C., Anthony, M. A., Baldrian, P., Finkbeiner, F., van den Hoogen, J., Kiers, T., Kohout, P., Hirt, E., Smith, G. R., & Crowther, T. W. (2022). Defending Earth’s terrestrial microbiome. Nature Microbiology, 7(11), 1717–1725. 10.1038/s41564-022-01228-3

Azevedo, O., Vanguelova, E., van der Linde, S., Ashwood, F., Fuentes-Montemayor, E., Wilson, C., Watts, K., & Park, K. J. (2026). Decades to establish, centuries to assemble: Soil microbial community development following woodland creation on agricultural land. Forest Ecology and Management, 621, 124228. 10.1016/j.foreco.2026.124228.

Banin, L. F., Raine, E. H., Rowland, L. M., Chazdon, R. L., Smith, S. W., Rahman, N. E. B., Butler, A., Philipson, C., Applegate, G. G., Axelsson, E. P., Budiharta, S., Chua, S. C., Cutler, M. E. J., Elliott, S., Gemita, E., Godoong, E., Graham, L. L. B., Hayward, R. M., Hector, A., … Burslem, D. F. R. P. (2022). The road to recovery: a synthesis of outcomes from ecosystem restoration in tropical and sub-tropical Asian forests. Philosophical Transactions of the Royal Society B: Biological Sciences, 378(1867). 10.1098/rstb.2021.0090

Barber, R. A., Underwood, B. C., Clarkson, J. J., Brekke, T. D., Weeks, T. L., Wilkinson, T., Wilkes, P., Egan, G., Lee, M. A., Openshaw, I., Castro, G., Moat, J., Bidartondo, M. I., Suz, L. M., & Kowal, J. (2026). Ectomycorrhizal community composition and extramatrical hyphal proportion predict soil carbon stocks at the landscape scale. Functional Ecology, 40(8). 10.1111/1365-2435.70369

Bárcena, T., Kiær, L., Vesterdal, L., Stefánsdóttir, H., Gundersen, P., & Sigurdsson, B. (2014). Soil carbon stock change following afforestation in Northern Europe: A meta-analysis. Global Change Biology, 20(8), 2393–2405. 10.1111/gcb.12576

Bashian-Victoroff, C. N., Burns, J. H., Carrino-Kyker, S. R., Hausman, C. E., Jahn, L. V., Lance, A. C., & Burke, D. J. (2026). Outcomes of commercial and forest soil inoculation on tree growth and mycorrhizal associations in a peri-urban forest restoration experiment. Restoration Ecology, 34(4), e70344. 10.1111/rec.70344

Bates, D., Mächler, M., Bolker, B., & Walker, S. (2015). Fitting linear mixed-effects models using lme4. Journal of Statistical Software, 67(1), 1–48. 10.18637/jss.v067.i01

Beerling, D. J., Kantzas, E. P., Lomas, M. R., Wade, P., Eufrasio, R. M., Renforth, P., Sarkar, B., Andrews, M. G., James, R. H., Pearce, C. R., Mercure, J.-F., Pollitt, H., Holden, P. B., Edwards, N. R., Khanna, M., Koh, L., Quegan, S., Pidgeon, N. F., Janssens, I. A., … Banwart, S. A. (2020). Potential for large-scale CO2 removal via enhanced rock weathering with croplands. Nature, 583(7815), 242–248. 10.1038/s41586-020-2448-9

Berns, A. E., Philipp, H., Narres, H. D., Burauel, P., Vereecken, H., & Tappe, W. (2008). Effect of gamma-sterilization and autoclaving on soil organic matter structure as studied by solid state NMR, UV and fluorescence spectroscopy. European Journal of Soil Science, 59(3), 540–550. 10.1111/j.1365-2389.2008.01016.x

Bonneville, S., Morgan, D. J., Schmalenberger, A., Bray, A., Brown, A., Banwart, S. A., & Benning, L. G. (2011). Tree-mycorrhiza symbiosis accelerate mineral weathering: Evidences from nanometer-scale elemental fluxes at the hypha–mineral interface. Geochimica et Cosmochimica Acta, 75(22), 6988–7005. 10.1016/j.gca.2011.08.041

Brekke, T. D., Weeks, T., Barber, R. A., Thomson, I., Gooda, R., Gargiulo, R., Delhaye, G., Andrew, C., Kowal, J., Bidartondo, M., & Suz, L. M. (2026). MycorrhizaTracer: A bioinformatic pipeline for fungi and plant classification of Sanger DNA sequences. bioRxiv, bioRxiv 10.64898/2026.04.23.720352.

Carrino-Kyker, S. R., Kluber, L. A., Petersen, S. M., Coyle, K. P., Hewins, C. R., DeForest, J. L., Smemo, K. A., & Burke, D. J. (2016). Mycorrhizal fungal communities respond to experimental elevation of soil pH and P availability in temperate hardwood forests. FEMS Microbiology Ecology, 92(3). 10.1093/femsec/fiw024

Clarkson, M. O., Larkin, C. S., Swoboda, P., Reershemius, T., Suhrhoff, T. J., Maesano, C. N., & Campbell, J. S. (2024). A review of measurement for quantification of carbon dioxide removal by enhanced weathering in soil. Frontiers in Climate, 6. 10.3389/fclim.2024.1345224

Conceição, L. T., Silva, G. N., Holsback, H. M. S., Oliveira, C. d. F., Marcante, N. C., Martins, É. d. S., Santos, F. L. d. S., & Santos, E. F. (2022). Potential of basalt dust to improve soil fertility and crop nutrition. Journal of Agriculture and Food Research, 10, 100443. 10.1016/j.jafr.2022.100443

Davison, J., Moora, M., Semchenko, M., Adenan, S. B., Ahmed, T., Akhmetzhanova, A. A., Alatalo, J. M., Al-Quraishy, S., Andriyanova, E., Anslan, S., Bahram, M., Batbaatar, A., Brown, C., Bueno, C. G., Cahill, J., Cantero, J. J., Casper, B. B., Cherosov, M., Chideh, S., … Öpik, M. (2021). Temperature and pH define the realised niche space of arbuscular mycorrhizal fungi. New Phytologist, 231(2), 763–776. 10.1111/nph.17240

De Cáceres, M., & Legendre, P. (2009). Associations between species and groups of sites: indices and statistical inference. Ecology, 90(12), 3566–3574. 10.1890/08-1823.1

Deslippe, J. R., Hartmann, M., Mohn, W. W., & Simard, S. W. (2011). Long-term experimental manipulation of climate alters the ectomycorrhizal community of Betula nana in Arctic tundra. Global Change Biology, 17(4), 1625–1636. 10.1111/j.1365-2486.2010.02318.x

Dickie, I., & Reich, P. (2005). Ectomycorrhizal fungal communities at forest edges. Journal of Ecology, 93(2), 244–255. 10.1111/j.1365-2745.2005.00977.x

Dickie, I. A., Montgomery, R. A., Reich, P. B., & Schnitzer, S. A. (2007). Physiological and phenological responses of oak seedlings to oak forest soil in the absence of trees. Tree Physiology, 27(1), 133–140. 10.1093/treephys/27.1.133

Doré, J., Perraud, M., Dieryckx, C., Kohler, A., Morin, E., Henrissat, B., Lindquist, E., Zimmermann, S. D., Girard, V., Kuo, A., Grigoriev, I. V., Martin, F., Marmeisse, R., & Gay, G. (2015). Comparative genomics, proteomics and transcriptomics give new insight into the exoproteome of the basidiomycete Hebeloma cylindrosporum and its involvement in ectomycorrhizal symbiosis. New Phytologist, 208(4), 1169–1187. 10.1111/nph.13546

Duell, E. B., O’Hare, A., & Wilson, G. W. T. (2023). Inoculation with native soil improves seedling survival and reduces non-native reinvasion in a grassland restoration. Restoration Ecology, 31(1), e13685. 10.1111/rec.13685

Durgapal, A., Pandey, A., & Palni, L. M. S. (2002). The use of rhizosphere soil for improved establishment of conifers at nursery stage for application in plantation programmes. Journal of Sustainable Forestry, 15(3), 57–73. 10.1300/J091v15n03_03

Endlweber, K., & Scheu, S. (2006). Establishing arbuscular mycorrhiza-free soil: A comparison of six methods and their effects on nutrient mobilization. Applied Soil Ecology, 34(2), 276–279. 10.1016/j.apsoil.2006.04.001

Ferlian, O., Goldmann, K., Eisenhauer, N., Tarkka, M. T., Buscot, F., & Heintz-Buschart, A. (2021). Distinct effects of host and neighbour tree identity on arbuscular and ectomycorrhizal fungi along a tree diversity gradient. ISME Communications, 1(1), 40. 10.1038/s43705-021-00042-y

Flykt, E., Timonen, S., & Pennanen, T. (2008). Variation of ectomycorrhizal colonisation in Norway spruce seedlings in Finnish forest nurseries. Silva Fennica, 42, 571. 10.14214/sf.234

Forster, E. J., Healey, J. R., Dymond, C., & Styles, D. (2021). Commercial afforestation can deliver effective climate change mitigation under multiple decarbonisation pathways. Nature Communications, 12(1), 3831. 10.1038/s41467-021-24084-x

Franco, A. R., Sousa, N. R., Ramos, M. A., Oliveira, R. S., & Castro, P. M. L. (2014). Diversity and persistence of ectomycorrhizal fungi and their effect on nursery-inoculated pinus pinaster in a post-fire plantation in northern portugal. Microbial Ecology, 68(4), 761–772. 10.1007/s00248-014-0447-9

Frew, A., Heuck, M. K., & Aguilar-Trigueros, C. A. (2023). Host filtering, not competitive exclusion, may be the main driver of arbuscular mycorrhizal fungal community assembly under high phosphorus. Functional Ecology, 37(7), 1856–1869.

Fu, M., Kang, Y., Zhou, J., Zhu, L., Wang, W., Lyu, T., & Lin, L. (2026). Contrasting elevational patterns of soil and root-associated fungal communities highlight host-driven filtering in *Quercus wutaishansea* forests. Frontiers in Plant Science, 17, 1825787. 10.3389/fpls.2026.1825787

Gardes, M., & Bruns, T. D. (1993). ITS primers with enhanced specificity for basidiomycetes--application to the identification of mycorrhizae and rusts. Molecular Ecology, 2(2), 113–118. 10.1111/j.1365-294x.1993.tb00005.x

Gaucher, Y., Tanaka, K., Johansson, D. J. A., Goll, D. S., & Ciais, P. (2025). Leveraging ecosystems responses to enhanced rock weathering in mitigation scenarios. Nature Communications, 16(1), 3021. 10.1038/s41467-025-58284-6

Gautier, M., Selosse, M.-A., Taschen, E., Bellanger, J.-M., Taudière, A., & Richard, F. (2026). Soil disturbance influences ectomycorrhizal fungal communities: an in-situ experiment under Mediterranean holm oak (Quercus ilex L.). Mycorrhiza, 36(4), 58. 10.1007/s00572-026-01302-2

Goldmann, K., Schröter, K., Pena, R., Schöning, I., Schrumpf, M., Buscot, F., Polle, A., & Wubet, T. (2016). Divergent habitat filtering of root and soil fungal communities in temperate beech forests. Scientific Reports, 6(1), 31439. 10.1038/srep31439

Goll, D. S., Ciais, P., Amann, T., Buermann, W., Chang, J., Eker, S., Hartmann, J., Janssens, I., Li, W., Obersteiner, M., Penuelas, J., Tanaka, K., & Vicca, S. (2021). Potential CO2 removal from enhanced weathering by ecosystem responses to powdered rock. Nature Geoscience, 14(8), 545–549. 10.1038/s41561-021-00798-x

Gomes, S. I. F., Gundersen, P., Bezemer, T. M., Barsotti, D., D’Imperio, L., Georgopoulos, K., Justesen, M. J., Rheault, K., Rosas, Y. M., Schmidt, I. K., Tedersoo, L., Vesterdal, L., Yu, M., Anslan, S., Aslani, F., Byriel, D. B., Christiansen, J., Hansen, S. H., Kasal, N., … Kepfer-Rojas, S. (2025). Soil microbiome inoculation for resilient and multifunctional new forests in post-agricultural landscapes. Global Change Biology, 31(1), e70031. 10.1111/gcb.70031

Gosling, P., Mead, A., Proctor, M., Hammond, J. P., & Bending, G. D. (2013). Contrasting arbuscular mycorrhizal communities colonizing different host plants show a similar response to a soil phosphorus concentration gradient. New Phytologist, 198(2), 546–556. 10.1111/nph.12169

Grove, S., Saarman, N. P., Gilbert, G. S., Faircloth, B., Haubensak, K. A., & Parker, I. M. (2019). Ectomycorrhizas and tree seedling establishment are strongly influenced by forest edge proximity but not soil inoculum. Ecological Applications, 29(3), e01867. 10.1002/eap.1867

Gryndler, M., Larsen, J., Hršelová, H., Řezáčová, V., Gryndlerová, H., & Kubát, J. (2006). Organic and mineral fertilization, respectively, increase and decrease the development of external mycelium of arbuscular mycorrhizal fungi in a long-term field experiment. Mycorrhiza, 16(3), 159–166. 10.1007/s00572-005-0027-4

Han, X., Li, Y., Li, Y., Du, X., Li, B., Li, Q., & Bezemer, T. M. (2022). Soil inoculum identity and rate jointly steer microbiomes and plant communities in the field. ISME Communications, 2(1), 59. 10.1038/s43705-022-00144-1

Hawkins, H.-J., Cargill, R. I., Van Nuland, M. E., Hagen, S. C., Field, K. J., Sheldrake, M., Soudzilovskaia, N. A., & Kiers, E. T. (2023). Mycorrhizal mycelium as a global carbon pool. Current Biology, 33(11), R560–R573. 10.1016/j.cub.2023.02.027.

Herinckx, P., Delhaye, G., Bidartondo, M. I., Gargiulo, R., Ghaffar, E., Ruhmann, C., Ticehurst, M., Andrews, C., Apuhtin, V., Lewis, C., Merilä, P., Vanguelova, E., Verstraeten, A., Wambsganss, J., Drouet, T., & Suz, L. M. (2026). High nitrogen deposition is associated with phosphorus-efficient ectomycorrhizas in Europe’s Scots pine forests. bioRxiv, 10.64898/2026.06.04.730184

Hong, S., Cong, N., Ding, J., Piao, S., Liu, L., Peñuelas, J., Chen, A., Quine, T. A., Zeng, H., & Houlton, B. Z. (2023). Effects of afforestation on soil carbon and nitrogen accumulation depend on initial soil nitrogen status. Global Biogeochemical Cycles, 37(1), e2022GB007490. 10.1029/2022GB007490

Hoysted, G. A., Kowal, J., Jacob, A., Rimington, W. R., Duckett, J. G., Pressel, S., Orchard, S., Ryan, M. H., Field, K. J., & Bidartondo, M. I. (2018). A mycorrhizal revolution. Current Opinion Plant Biology, 44, 1–6. 10.1016/j.pbi.2017.12.004

Hsieh, T. C., Ma, K. H., & Chao, A. (2016). iNEXT: an R package for rarefaction and extrapolation of species diversity (Hill numbers). Methods in Ecology and Evolution, 7(12), 1451–1456. 10.1111/2041-210X.12613

Johansson, E. M., Fransson, P. M. A., Finlay, R. D., & van Hees, P. A. W. (2009). Quantitative analysis of soluble exudates produced by ectomycorrhizal roots as a response to ambient and elevated CO2. Soil Biology and Biochemistry, 41(6), 1111–1116. 10.1016/j.soilbio.2009.02.016

Johnson, N. C., Angelard, C., Sanders, I. R., & Kiers, E. T. (2013). Predicting community and ecosystem outcomes of mycorrhizal responses to global change. Ecology Letters, 16(s1), 140–153. 10.1111/ele.12085

Jung, S. C., Martinez-Medina, A., Lopez-Raez, J. A., & Pozo, M. J. (2012). Mycorrhiza-induced resistance and priming of plant defenses. Journal of Chemical Ecology, 38, 651–664. 10.1007/s10886-012-0134-6

Kafy, A. A., Saha, M., Fattah, M. A., Rahman, M. T., Duti, B. M., Rahaman, Z. A., Bakshi, A., Kalaivani, S., Nafiz Rahaman, S., & Sattar, G. S. (2023). Integrating forest cover change and carbon storage dynamics: Leveraging Google Earth Engine and InVEST model to inform conservation in hilly regions. Ecological Indicators, 152, 110374. 10.1016/j.ecolind.2023.110374

Kaiser, C., & Anthony, M. A. (2025). The role of ectomycorrhizal functional diversity in mediating soil carbon cycling under global change. New Phytologist, 248(4), 1605–1608. 10.1111/nph.70559

King, W. L., Grandinette E. M., Trase, O., Rolon, M. L., Salis H. M., Wood, H., & Bell T. H. (2024). Autoclaving is at least as effective as gamma irradiation for biotic clearing and intentional microbial recolonization of soil. mSphere, 9(7), e00476–00424. 10.1128/msphere.00476-24

King, W. L., Richards, S. C., Kaminsky, L. M., Bradley, B. A., Kaye, J. P., & Bell, T. H. (2023). Leveraging microbiome rediversification for the ecological rescue of soil function. Environmental Microbiome, 18(1), 7. 10.1186/s40793-023-00462-4

Kipfer, T., Moser, B., Egli, S., Wohlgemuth, T., & Ghazoul, J. (2011). Ectomycorrhiza succession patterns in Pinus sylvestris forests after stand-replacing fire in the Central Alps. Oecologia, 167(1), 219–228. 10.1007/s00442-011-1981-5

Kluber, L. A., Carrino-Kyker, S. R., Coyle, K. P., DeForest, J. L., Hewins, C. R., Shaw, A. N., Smemo, K. A., & Burke, D. J. (2012). Mycorrhizal response to experimental pH and P manipulation in acidic hardwood forests. PLoS One, 7(11), e48946. 10.1371/journal.pone.0048946

Kowal, J., Arrigoni, E., & Lane, S. (2020). Acidified Blue Ink-staining Procedure for the Observation of Fungal Structures Inside Roots of Two Disparate Plant Lineages. Bio Protocol, 10(20), e3786. 10.21769/BioProtoc.3786

Laganiere, J., Angers, D. A., & Pare, D. (2010). Carbon accumulation in agricultural soils after afforestation: a meta-analysis. Global Change Biology, 16(1), 439–453.

Lance, A. C., Burke, D. J., Hausman, C. E., & Burns, J. H. (2019). Microbial inoculation influences arbuscular mycorrhizal fungi community structure and nutrient dynamics in temperate tree restoration. Restoration Ecology, 27(5), 1084–1093. 10.1111/rec.12962

Larsson, J. (2024). Area-Proportional Euler and Venn Diagrams with Ellipses [R package eulerr version 7.0. 2]. CRAN: Contributed Packages.

Leff, J. W., Jones, S. E., Prober, S. M., Barberán, A., Borer, E. T., Firn, J. L., Harpole, W. S., Hobbie, S. E., Hofmockel, K. S., Knops, J. M. H., McCulley, R. L., La Pierre, K., Risch, A. C., Seabloom, E. W., Schütz, M., Steenbock, C., Stevens, C. J., & Fierer, N. (2015). Consistent responses of soil microbial communities to elevated nutrient inputs in grasslands across the globe. Proceedings of the National Academy of Sciences, 112(35), 10967–10972. 10.1073/pnas.1508382112

Lewis, A. L., Sarkar, B., Wade, P., Kemp, S. J., Hodson, M. E., Taylor, L. L., Yeong, K. L., Davies, K., Nelson, P. N., Bird, M. I., Kantola, I. B., Masters, M. D., DeLucia, E., Leake, J. R., Banwart, S. A., & Beerling, D. J. (2021). Effects of mineralogy, chemistry and physical properties of basalts on carbon capture potential and plant-nutrient element release via enhanced weathering. Applied Geochemistry, 132, 105023. 10.1016/j.apgeochem.2021.105023

Liu, M., Shen, Y., Li, Q., Xiao, W., & Song, X. (2021). Arbuscular mycorrhizal fungal colonization and soil pH induced by nitrogen and phosphorus additions affects leaf C:N:P stoichiometry in Chinese fir (Cunninghamia lanceolata) forests. Plant and Soil, 461(1), 421–440. 10.1007/s11104-021-04831-1

Lüdecke, D., Ben Shachar, M., Patil, I., Waggoner, P., & Makowski, D. (2021). performance: An R package for assessment, comparison and testing of statistical models. Journal of Open Source Software, 6, 3139. 10.21105/joss.03139

Lybrand, R. A., Qafoku, O., Bowden, M. E., Hochella M. F. Jr., Kovarik, L., Perea, D. E., Qafoku, N. P., Schroeder, P. A., Wirth, M. G., & Zaharescu, D. G. (2022). Fungal hyphae develop where titanomagnetite inclusions reach the surface of basalt grains. Scientific Reports, 12(1), 3407. 10.1038/s41598-021-04157-z

Martínez-García, L. B., Richardson, S. J., Tylianakis, J. M., Peltzer, D. A., & Dickie, I. A. (2015). Host identity is a dominant driver of mycorrhizal fungal community composition during ecosystem development. New Phytologist, 205(4), 1565–1576. 10.1111/nph.13226

Mc Kay, H. M., Harrison, A. J., Connolly, T., Forster, J., Gregory, S., Smith, E., & Clark, J. (2021). Early performance of eight broadleaved species grown as short rotation forest in England. Forestry: An International Journal of Forest Research, 95(1), 13–27. 10.1093/forestry/cpab033

McMahen, K., Anglin, C. D., Lavkulich, L. M., Grayston, S. J., & Simard, S. W. (2022). Small-volume additions of forest topsoil improve root symbiont colonization and seedling growth in mine reclamation. Applied Soil Ecology, 180, 104622. 10.1016/j.apsoil.2022.104622

Melvin, A. M., Mack, M. C., Johnstone, J. F., McGuire, A. D., Genet, H., & Schuur, E. A. G. (2015). Differences in ecosystem carbon distribution and nutrient cycling linked to forest tree species composition in a mid-successional boreal forest. Ecosystems, 18(8), 1472–1488. 10.1007/s10021-015-9912-7

Nguyen, N. H., Song, Z., Bates, S. T., Branco, S., Tedersoo, L., Menke, J., Schilling, J. S., & Kennedy, P. G. (2016). FUNGuild: An open annotation tool for parsing fungal community datasets by ecological guild. Fungal Ecology, 20, 241–248. 10.1016/j.funeco.2015.06.006

Oksanen, J., Kindt, R., Legendre, P., O’Hara, B., Stevens, M. H. H., Oksanen, M. J., & Suggests, M. A. S. S. (2007). The vegan package. Community ecology package, 10(631-637), 719.

Palfner, G., Casanova-Katny, M. A., & Read, D. J. (2005). The mycorrhizal community in a forest chronosequence of Sitka spruce [Picea sitchensis (Bong.) Carr.] in Northern England. Mycorrhiza, 15(8), 571–579. 10.1007/s00572-005-0364-3

Parasquive, V., Brisson, J., Laliberté, E., & Chagnon, P. L. (2024). Limited impact of soil inocula from arbuscular and ectomycorrhizal-dominated sites on root morphology and growth of four tree seedling species from a temperate deciduous forest. Plant and Soil, 511, 1137–1149. 10.1007/s11104-024-07043-5

Pellitier, P. T., Zak, D. R., Argiroff, W. A., & Upchurch, R. A. (2021). Coupled shifts in ectomycorrhizal communities and plant uptake of organic nitrogen along a soil gradient: an isotopic perspective. Ecosystems, 24(8), 1976–1990. 10.1007/s10021-021-00628-6

Pena, R., Bluhm, S. L., Ammerschubert, S., Agüi-Gonzalez, P., Rizzoli, S. O., Scheu, S., & Polle, A. (2023). Mycorrhizal C/N ratio determines plant-derived carbon and nitrogen allocation to symbiosis. Communications Biology, 6(1), 1230. 10.1038/s42003-023-05591-7

Penn, C. J., & Camberato, J. J. (2019). A critical review on soil chemical processes that control how soil ph affects phosphorus availability to plants. Agriculture, 9(6), 120. https://www.mdpi.com/2077-0472/9/6/120

Percival-Alwyn, L., Barnes, I., Clark, M.D., Cockram, J., Coffey, M.P., Jones, S., Kersey, P.J., Kidner, C.A., Kosiol, C., Li, B., Marsh, W.A., Zhou, J., Caccamo, M., Milne, I. (2024).UKCropDiversity-HPC: A collaborative high-performance computing resource approach for sustainable agriculture and biodiversity conservation. *Plants, People*, Planet. 10.1002/ppp3.10607

Psistaki, K., Tsantopoulos, G., & Paschalidou, A. K. (2024). An overview of the role of forests in climate change mitigation. Sustainability, 16(14), 6089. 10.3390/su16146089

Quoreshi, A. M., Piché, Y., & Khasa, D. P. (2008). Field performance of conifer and hardwood species 5 years after nursery inoculation in the Canadian Prairie Provinces. New Forests, 35(3), 235–253. 10.1007/s11056-007-9074-3

R Core Team. (2020). RA language and environment for statistical computing, R Foundation for Statistical. Computing.

Riedo, J., Wettstein, F. E., Rösch, A., Herzog, C., Banerjee, S., Büchi, L., Charles, R., Wächter, D., Martin-Laurent, F., & Bucheli, T. D. (2021). Widespread occurrence of pesticides in organically managed agricultural soils—the ghost of a conventional agricultural past? Environmental Science & Technology, 55(5), 2919–2928.

Rockström, J., Gaffney, O., Rogelj, J., Meinshausen, M., Nakicenovic, N., & Schellnhuber, H. J. (2017). A roadmap for rapid decarbonization. Science, 355(6331), 1269–1271. 10.1126/science.aah3443

Rodriguez-Ramos, J. C., Cale, J. A., Cahill, J. F. Jr., Erbilgin, N., & Karst, J. (2022). Soil transfers from intact to disturbed boreal forests neither alter ectomycorrhizal fungal communities nor improve pine seedling performance. Journal of Applied Ecology, 59(9), 2430–2439. 10.1111/1365-2664.14245

Rognes, T., Flouri, T., Nichols, B., Quince, C., & Mahé, F. (2016). VSEARCH: a versatile open source tool for metagenomics. PeerJ, 4, e2584. 10.7717/peerj.2584

Rosenstock, N., Ellstrom, M., Oddsdottir, E., Sigurdsson, B., & Wallander, H. (2018). Carbon sequestration and community composition of ectomycorrhizal fungi across a geothermal warming gradient in an Icelandic spruce forest. Fungal Ecology, 30, 1e11. 10.1016/j.funeco.2018.05.010.

Sayers, E. W., Beck, J., Bolton, E. E., Brister, J. R., Chan, J., Connor, R., Feldgarden, M., Fine, A. M., Funk, K., Hoffman, J., Kannan, S., Kelly, C., Klimke, W., Kim, S., Lathrop, S., Marchler-Bauer, A., Murphy, T. D., O’Sullivan, C., Schmieder, E., … Pruitt, K. D. (2025). Database resources of the National Center for Biotechnology Information in 2025. Nucleic Acids Res, 53(D1), D20–d29. 10.1093/nar/gkae979

Smits, M. M., Bonneville, S., Benning, L. G., Banwart, S. A., & Leake, J. R. (2012). Plant-driven weathering of apatite – the role of an ectomycorrhizal fungus. Geobiology, 10(5), 445–456. 10.1111/j.1472-4669.2012.00331.x

St-Denis, A., Kneeshaw, D., Bélanger, N., Simard, S., Laforest-Lapointe, I., & Messier, C. (2017). Species-specific responses to forest soil inoculum in planted trees in an abandoned agricultural field. Applied Soil Ecology, 112, 1–10. 10.1016/j.apsoil.2016.12.008

Straub, D., Blackwell, N., Langarica-Fuentes, A., Peltzer, A., Nahnsen, S., & Kleindienst, S. (2020). Interpretations of environmental microbial community studies are biased by the selected 16s rRNA (gene) amplicon sequencing pipeline. Frontiers in Microbiology, 11, 550420. 10.3389/fmicb.2020.550420

Suz, L. M., Barsoum, N., Benham, S., Dietrich, H. P., Fetzer, K. D., Fischer, R., García, P., Gehrman, J., Kristöfel, F., Manninger, M., Neagu, S., Nicolas, M., Oldenburger, J., Raspe, S., Sánchez, G., Schröck, H. W., Schubert, A., Verheyen, K., Verstraeten, A., & Bidartondo, M. I. (2014). Environmental drivers of ectomycorrhizal communities in Europe’s temperate oak forests. Molecular Ecology, 23(22), 5628–5644. 10.1111/mec.12947

Taniguchi, T., Yuzawa, T., HuiPing, M., Yamamoto, F., & Yamanaka, N. (2021). Plantation soil inoculation combined with straw checkerboard barriers enhances ectomycorrhizal colonization and subsequent growth of nursery grown Pinus tabulaeformis seedlings in a dryland. Ecological Engineering, 163, 106191. 10.1016/j.ecoleng.2021.106191

Tedersoo, L., & Anslan, S. (2019). Towards PacBio-based pan-eukaryote metabarcoding using full-length ITS sequences. Environmental Microbiology Reports, 11(5), 659–668. 10.1111/1758-2229.12776

Tedersoo, L., & Bahram, M. (2019). Mycorrhizal types differ in ecophysiology and alter plant nutrition and soil processes. Biological Reviews, 94(5), 1857–1880. 10.1111/brv.12538

Tedersoo, L., Hansen, K., Perry, B. A., & Kjøller, R. (2006). Molecular and morphological diversity of pezizalean ectomycorrhiza. New Phytologist, 170(3), 581–596. 10.1111/j.1469-8137.2006.01678.x

Tedersoo, L., Kõljalg, U., Hallenberg, N., & Larsson, K. H. (2003). Fine scale distribution of ectomycorrhizal fungi and roots across substrate layers including coarse woody debris in a mixed forest. New Phytologist, 159(1), 153–165. 10.1046/j.1469-8137.2003.00792.x

Tedersoo, L., & Lindahl, B. (2016). Fungal identification biases in microbiome projects. Environ Microbiol Rep, 8(5), 774–779. 10.1111/1758-2229.12438

Teste, F. P., Simard, S. W., & Durall, D. M. (2009). Role of mycorrhizal networks and tree proximity in ectomycorrhizal colonization of planted seedlings. Fungal Ecology, 2(1), 21–30. 10.1016/j.funeco.2008.11.003

Tibbett, M., Sanders, F. E., Cairney, J. W. G., & Leake, J. R. (1999). Temperature regulation of extracellular proteases in ectomycorrhizal fungi (Hebeloma spp.) grown in axenic culture. Mycological Research, 103(6), 707–714.

Tourville, J. C., Horton, T. R., & Dovciak, M. (2024). Mycorrhizal fungi as critical biotic filters for tree seedling establishment during species range expansions. Ecological Monographs, 94(4), e1634. 10.1002/ecm.1634

Truog, E. (1947). Soil reaction influence on availability of plant nutrients. Soil Science Society of America Journal, 11, 305–308. 10.2136/sssaj1947.036159950011000C0057x

van der Linde, S., Suz, L. M., Orme, C. D. L., Cox, F., Andreae, H., Asi, E., Atkinson, B., Benham, S., Carroll, C., Cools, N., De Vos, B., Dietrich, H. P., Eichhorn, J., Gehrmann, J., Grebenc, T., Gweon, H. S., Hansen, K., Jacob, F., Kristöfel, F., … Bidartondo, M. I. (2018). Environment and host as large-scale controls of ectomycorrhizal fungi. Nature, 558(7709), 243–248. 10.1038/s41586-018-0189-9

Wallander, H., Hagerberg, D., & Åberg, G. (2006). Uptake of 87Sr from microcline and biotite by ectomycorrhizal fungi in a Norway spruce forest. Soil Biology and Biochemistry, 38(8), 2487–2490. 10.1016/j.soilbio.2006.02.002

Wang, X., Kou, Y., Liu, J., Zhao, W., & Liu, Q. (2023). Soil microbial legacy determines mycorrhizal colonization and root traits of conifer seedlings during subalpine forest succession. Plant and Soil, 485(1), 361–375. 10.1007/s11104-022-05835-1

Waring, B. G., Averill, C., Bidartondo, M., Suz, L. M., Beerling, D. J., Crowther, T. W., Jones, G., Lancastle, L., Epihov, D. Z., Clayton, K., Gobelius, L., Lindsay, O., Steidinger, B., Allen, H., & Nicholls, C. (2026). Microbiome manipulation and enhanced weathering influence tree growth in reforestation. Communications Sustainability, 1(1), 102. 10.1038/s44458-026-00103-0

Waring, B. G., Smith, K. R., Belluau, M., Khlifa, R., Messier, C., Munson, A., & Paquette, A. (2022). Soil carbon pools are affected by species identity and productivity in a tree common garden experiment. Frontiers in Forests and Global Change, 5, 1032321. 10.3389/ffgc.2022.1032321

Wasyliw, J., Fellrath, E. G., Pec, G. J., Cale, J. A., Franklin, J., Thomasson, C., Erbilgin, N., & Karst, J. (2023). Soil inoculation of lodgepole pine seedlings alters root-associated fungal communities but does not improve seedling performance in beetle-killed pine stands. Restoration Ecology, 31(1), e13663. 10.1111/rec.13663

Wen, Z., Shi, L., Tang, Y., Hong, L., Xue, J., Xing, J., Chen, Y., & Nara, K. (2018). Soil spore bank communities of ectomycorrhizal fungi in endangered Chinese Douglas-fir forests. Mycorrhiza, 28(1), 49–58. 10.1007/s00572-017-0800-1

White, T. J., Bruns, T. D., Lee, S. B., & Taylor, J. W. (1990). Amplificationand direct sequencing of fungal ribosomal RNA genes for phyloge-netics. In M. A. Innis, D. H. Gelfand, J. J. Sninsky, & T. J. White (Eds.),PCR protocols: A guide to methods and applications (pp. 315–322). NewYork, NY: Academic Press

Wubs, E. R., van der Putten, W. H., Bosch, M., & Bezemer, T. M. (2016). Soil inoculation steers restoration of terrestrial ecosystems. Nature Plants, 2, 16107. 10.1038/nplants.2016.107

Yan, X., Zhou, L., Chen, Y., Liu, R., Guo, L., Li, N., Kang, A., Zhai, K., Zhou, G., & Zhou, X. (2025). Ectomycorrhizal fungi explain more variation in rhizosphere nutrient availability than root traits in temperate forests. Applied Soil Ecology, 207, 105923. 10.1016/j.apsoil.2025.105923

Yuan, X.-L., Zhu, X.-T., Shi, Y., Miao, Y., Zhang, R.-G., Li, P., & Shen, C. (2025). Soil microbiomes in degraded grasslands: Assembly, function, and application. Grassland Research, 4(4), 352–365. 10.1002/glr2.70023

Zang, Z., Li, Y., Wang, Y., Zhang, Y., Deng, S., Guo, X., Yang, K., & Zhao, W. (2024). Contrasting roles of plant, bacterial, and fungal diversity in soil organic carbon accrual during ecosystem restoration: A meta-analysis. Science of the Total Environment, 930, 172767. 10.1016/j.scitotenv.2024.172767

Zhao, D.-X., Bai, Z., Yuan, Y.-W., Li, S.-A., Wei, Y.-L., & Yuan, H.-S. (2025). Ectomycorrhizal fungal community varies across broadleaf species and developmental stages. Scientific Reports, 15(1), 6955. 10.1038/s41598-025-91411-3

Zhou, J., Gube, M., Holz, M., Song, B., Shan, I., Shi, L., Kuzyakov, Y., Dippold, M. A., & Pausch, J. (2022). Ectomycorrhizal and non-mycorrhizal rhizosphere fungi increase root-derived C input to soil and modify enzyme activities: A 14C pulse labelling of Picea abies seedlings. Plant, Cell & Environment, 45(10), 3122–3133. 10.1111/pce.14413

