## Supplementary_1-4 for "Host identity and fungal source pools shape mycorrhizal assembly under carbon removal interventions in large-scale afforestation"

**S1 Written methods**

***S1.1 Plantation management***

Plantation vegetation was managed by applying glyphosate (Gallup Biograde 360; maximum application rate 10 L $ha^{-1}$) between 2022 (year 2) and 2024 (year 4) to control competing grasses and weeds. Plots are irregularly shaped to accommodate landscape features, and treatments were randomly assigned to plots within each block

***S1.2 Basalt treatment***

The basalt amendment consisted of crushed basaltic andesite sourced as a quarry by-product from Builth Wells Quarry, ca. 35 km from the experimental site. The material contained 7.2% Mg, 6.0% Ca, 0.4% K and 0.3% P by mass. For basalt application, first deployment in year 2 was across the whole plot prior to planting; the second in year 3 was inter-tree row (due to the challenging nature of a second deployment access after trees had been planted and had grown).

##### *S1.3 Broadleaf sampling*

Broadleaf sampling groups (Table S1): As broadleaf plantations comprised six tree species, species were divided between two groups of four blocks: alder (EcM), birch (EcM) and cherry (AM) were sampled in blocks A, B, E and G, and oak (EcM), aspen (EcM) and rowan (AM) in blocks C, D, F and H. Consequently, each broadleaf species was represented in four blocks per treatment.

##### *S1.4 Aspen mycorrhizal status*

Aspen was initially processed as a dual-mycorrhizal host, sampled for both EcM and AM colonisation. After Year 2 sampling, in which only EcM colonisation was detected, Aspen was treated as an EcM host. The reduced sample number in Year 2 therefore reflects root tips that were processed as AM samples but showed no AM/EcM colonisation.

##### *S1.5 Root sampling of potential sources of mycorrhizal fungi colonising plantation roots*

To identify potential sources of mycorrhizal fungi colonising plantation roots, we sampled roots from the source Maelor Nursery in April 2021 (nursery-grown seedlings), woodland inoculum sources in November 2021 (sites from which donor soil was obtained), surrounding hedgerows and non-forested pasture controls.

Prior to planting, 30 saplings per tree species were randomly sampled from Maelor Nursery in April 2021, matching the per-species sampling effort of the plantation sampling campaigns. Woodland inoculum sources were sampled in November 2021, with 12 soil cores collected along two transects between neighbouring EcM host trees at each site, covering the area where the soil inoculum was taken. Hedgerows and pasture controls were sampled during year 2 and year 4 field campaigns. Eight soil cores were collected at 1 m intervals along a 7 m transect from six hedgerows adjacent to the plantation (5 - 10 m away). For the pasture controls, four evenly spaced cores were collected along the diagonal of the plots. Sample size reflected maximum resource and operational efforts for representative sampling across all blocks.

***S1.6 Sampling of the woodland inoculum sources***

Only EcM trees were sampled, the woodland inoculum source was dominated by beech and oak. We also we expected the inoculum to primarily influence the EcM fungal communities in the plantation, given that ECM fungi were likely less dominant in the pasture while AM fungi predominated and broadleaf woodland inoculum sources primary contains EcM hosts.

At the spruce woodland inoculum source, sampling was conducted in the donor woodland, where trees had recently been felled.

***S1.7 Mycorrhizal assessment***

Soil cores were stored at 1-5°C for up to two weeks before processing. Soil was washed over a 500 μm sieve with tap water and roots collected under a dissecting microscope for 10 min and transferred to deionised water. For plantation samples, the four cores collected per tree were pooled before processing. Twelve roots were randomly selected from each sample using a grid-based approach and processed according to mycorrhizal type.

For EcM hosts, one colonised root tip (mycorrhiza) was selected from each of the twelve roots and cleared of surface debris. Tree host identity was confirmed by root morphology. Ectomycorrhizal mantle colour, hydrophobicity and morphological traits associated with soil exploration type, including the presence or absence of hyphae and rhizomorphs (Agerer, 2001) were recorded. Each ectomycorrhiza was transferred to a 96-well plate containing 10 μL Extract-N-Amp extraction buffer (Sigma-Aldrich) for further DNA analysis.

For AM plantation tree hosts and pasture controls (mostly grasses), roots at least 2 cm in length were selected and divided into two 1 cm segments. One segment was preserved in 2% CTAB for molecular analysis, with up to 12 roots (1 cm segments) per sample pooled into one tube containing CTAB prior to DNA extraction. The other segment was placed in 70% ethanol for root staining, following methods from Kowal et al. (2020), and subsequent root colonisation assessment.

Nursery roots were processed separately. For EcM hosts, mycorrhizas were selected according to distinct morphotypes, with the number analysed per seedling depending on the number of morphotypes present. For AM hosts, one root was collected per seedling and pooled by tree species for DNA extraction. Sample totals are in Table S2.

***S1.8 Molecular analysis of ectomycorrhizal fungi***

Genomic DNA was extracted from individual ectomycorrhizas using the Extract-N-Amp Plant PCR Kit (Sigma-Aldrich) following the manufacturer's instructions. The fungal internal transcribed spacer (ITS) region of the ribosomal DNA was amplified using the ITS1F (5'-

CTTGGTCATTTAGAGGAAGTAA) and ITS4 (5'-TCCTCCGCTTATTGATATGC) primer pair (White, 1990; Gardes & Bruns, 1993). The PCR products were cleaned using Exo1 and FAST-AP enzymes and sequenced bidirectionally on an Applied Biosystems 3730xl DNA Analyzer (Azenta Life Sciences).

The subsequent bioinformatics and computational analyses were performed on Crop Diversity HPC, described by Percival-Alwyn et al. (2024). Sanger chromatograms obtained from each individual ectomycorrhiza across samples were processed using the MycorrhizaTracer pipeline (v2.3.0; Brekke et al. (2026), *bioRxiv 10.64898/2026.04.23.720352*) with a minimum Phred quality score of 25, a standard deviation threshold of 0.1, and a minimum salvage read length of 100 bp. MycorrhizaTracer uses VSEARCH (Rognes et al., 2016) to match high-quality sequences to the reference sequence database UNITE v10 (Abarenkov et al., 2024). Sequences matching ≥97% to a sequence in the database were assigned the corresponding SH (species hypothesis) number. The remaining sequences were subsequently clustered into operational taxonomic units (OTUs) at 97% sequence similarity and considered as *de novo*.

Lower-quality sequences were manually inspected and trimmed where necessary and retained if they shared ≥97% sequence similarity with an existing OTU in the dataset. Taxonomic assignments were based on sequence similarity to reference databases using BLASTN (Sayers et al., 2025), with assignments made to species (≥97% identity), genus (94.0-96.9%), family (91.0-93.9%), and order or higher (<91.0%) (Deslippe et al., 2011). For each OTU, the taxonomic identity of the centroid sequence was assigned to all sequences within the cluster. Low-quality singleton and doubleton sequences with <90% sequence identity to reference databases were excluded from further analyses. This removed all low-quality, poorly represented sequences from the dataset. Mycorrhizal guild assignments were determined using FUNGuild (vGuilds_v1.1) (Nguyen et al., 2016) and non-EcM fungi were excluded from the final dataset. The resultant dataset consists of one EcM taxon assigned to each mycorrhiza.

##### *S1.9 Molecular analysis of arbuscular mycorrhizal fungi*

Genomic DNA was extracted from root samples using the DNeasy 96 Plant Kit (QIAGEN) according to the manufacturer's instructions. The full ITS region and part of the SSU were amplified using the eukaryotic specific primers NS1B1ngs (5'-CCTNGTTGATYCTGCCAGT; Tedersoo & Anslan, 2019) and ITS4ngsUni (5′-CCTSCSCTTANTDATATGC; Tedersoo & Lindahl, 2016). The PCR products were purified using AMPure XP beads. AM roots were sequenced on a PacBio Revio platform (Azenta Life Sciences) to enable simultaneal detection of multiple potential AM species per root and the plant host.

PacBio circular consensus sequencing reads were demultiplexed using lima, PacBio's demultiplexing software (available at lima.how) and then processed using the nf-core/ampliseq pipeline (v2.18.0) (Straub et al., 2020). The parameters for Ampliseq are available in Suppl.1.3. Primer sequences were removed using Cutadapt, allowing a minimum overlap of three bases and a maximum error rate of 10%; reads lacking both primer sequences were discarded. Quality filtering retained reads with a minimum quality score of Q25, a maximum expected error rate of five, and lengths between 1,900 and 3,000 bp. Samples containing fewer than 1,000 reads after filtering were excluded from downstream analyses.

Quality-filtered sequences were clustered into OTUs at 97% sequence similarity using VSEARCH (Rognes et al., 2016). The OTUs represented by fewer than 10 reads, occurring or in fewer than two samples. Taxonomic assignments were performed against the UNITE reference database (v10.0), with SH assigned if available. The resulting OTU table and taxonomic assignments were imported into R (v2023.12.0) for downstream analyses using the phyloseq (McMurdie & Holmes, 2013). To retain only AM fungi, the final dataset was filtered to include taxa belonging to Glomeromycota and Endogonales (Hoysted et al., 2018).

##### *S1.10 Soil mycorrhizal fungal communities*

Raw sequence data from DNA extractions of the plantation soils, soil inoculum, and pasture control soils were obtained from Waring et al. (2026). Soils had been collected from the rooting zone of trees in year 2. Total DNA was extracted, and the ITS region (primers ITS9munngs, GTACACACCGCCCGTCG, and ITS4ng, CGCCTSCSCTTANTDATATGC) was amplified and sequenced using circular consensus sequencing on a PacBio platform. We demultiplexed the sequences using the Cutadapt (v4.5) yielding 4.4 million reads that once processed resulted in 1.7 million reads after filtering. After quality filtering this led to an average of 3335 reads per sample (1-16,430). To identify mycorrhizal fungi in these soils, sequences were processed using the nf-core/ampliseq pipeline, as previously described.

##### *S1.11 Soil pH*

In year 4, three soil cores were taken per plot, across blocks A-D (not all blocks were sampled for pH due to equipment constraints), at regular intervals across a diagonal transect of each plot. The pH was subsequently measured with a pH probe (HI-99121, Hanna instruments UK) using a 5g soil slurry following manufacturer's instructors with the pH recorded three times per core.

##### *S1.12 Plantation pH and biomass measurements*

**Soil pH:** In year 4, three soil cores were taken per plot, across blocks A-D, at regular intervals across a diagonal transect of each plot. The pH was subsequently measured with a probe (HI-99121, Hanna instruments, UK) using a 5 g soil slurry following manufacturer's instructors.
**Plantation biomass:** Biomass data were retrieved from Waring et al. (2026). Briefly, tree growth measurements were recorded annually from 100 trees within each plot across all experimental blocks by researchers and trained citizen science volunteers. Because individual biomass measurements could not be matched to the trees sampled for mycorrhizal analyses, biomass was averaged for each tree species within each plot before analysis.

##### *S1.13 Additional detail on community assessment and statistical analyses*

***Software.*** Analyses used vegan v2.7.1 (Oksanen et al., 2013), lme4 v1.1.37 (Bates et al., 2015), glmmTMB v1.1.13, iNEXT v3.0.1 (Hsieh et al., 2016), indicspecies (De Cáceres & Legendre, 2009), eulerr v7.0.2 (Larsson, 2024), performance v0.15.0 (Lüdecke et al., 2021) and mgcv v1.9.1. PCoA was performed with cmdscale.

***Basalt x inoculum interaction:*** Basalt x inoculum interactions were tested in all models; as none were detected and none were hypothesised *a priori*, additive models are also reported.

***Community data and sample sizes.*** At plantation level, 64 plots (32 spruce, 32 broadleaf) were sampled in each of year 2 and year 4. Sample sizes at tree-species level were: oak, n = 32; birch, n = 30; alder, n = 29; aspen, n = 24; rowan, n = 32; cherry, n = 30; Sitka spruce, n = 64 (Table S3). EcM data were converted to relative abundance using decostand (method = "total"). For broadleaf matrices incorporating host identity, the two EcM seedlings per plot row were split to one row per species per plot, and for AM seedlings one row per species per plot. Metadata comprised treatment, and means of continuous variables, and the total number of mycorrhizas sampled. As different AM species were sampled in different blocks, the host/block effect cannot be partitioned for AM communities, therefore host effects (separate from block) are not reported in AM analyses.

***Community composition H1 & H3*** Where missing samples unbalanced the design, samples were omitted to restore balance before PERMANOVA. Basalt × inoculum interactions were tested in all models; because none were detected and none were hypothesised a priori, additive models are also reported. Variance partitioning used varpart, and unique fractions were tested by db-RDA with the permutation schemes described in S1.14. Principal coordinates analysis (PCoA; cmdscale v4.3) was performed on dissimilarity matrices (R Core

Team, 2020), separately by plantation type, mycorrhizal type and sampling year, and across both plantations combined.

Species vectors were fitted using envfit (999 permutations) to identify taxa significantly associated with ordination space. PCoA axis scores were used as response variables in linear and linear mixed-effects models using the lmer function (lme4 v1.1.37) (Bates et al., 2025), testing treatment effects, and as predictors of biomass.

Alpha diversity was estimated using the iNEXT framework (v3.0.1) using Hill numbers (Hsieh et al., 2016): species richness ($q$ = 0), Shannon diversity ($q$ = 1), and Simpson diversity ($q$ = 2), with asymptotic species richness and associated 95% confidence intervals calculated per assemblage. Treatment differences were tested by one-way ANOVA or Kruskal-Wallis, according to normality assessed by Shapiro-Wilk.

Indicator species analysis was performed using the indicspecies package (v1.8.0) (De Cáceres & Legendre, 2009). Indicator species are presented as stacked bar charts, showing the most abundant OTUs overall (ranked by summed relative abundance across all plots), and all remaining OTUs pooled into a single "Other" category within each plot. Plot-level relative abundances were then averaged across replicate plots within each combination of treatment (control, basalt, inoculum, inoculum + basalt) and sampling year (dplyr, forcats v1.0.1) and ggplot2 (v3.5.2).

***Biomass models H4:*** For PCoA scores centred within each tree species, used to represent within-host EcM variation in models, the centred score for sample $i$ of species $s$ was calculated as $V_i - \bar{V}_s$, where $\bar{V}_s$ is the mean PCoA score of all samples of that species. Centred scores are orthogonal to host identity, so their coefficients are estimated solely from variation among conspecific samples, while among-host differences in composition are absorbed by the host species term. Model assumptions were checked using *performance* (v0.15.0).

Biomass gradients across fungal community composition were visualised by fitting biomass as a two-dimensional smooth surface over the EcM ordination (generalised additive model; GAM; mgcv pacakge v1.9-1; k=10). Biomass (kg) was not square-root transformed, to allow visualisation of real biomass gradients.

To identify EcM taxa associated with biomass, plot-level OTU data were converted to presence-absence and a binomial generalised linear model (GLM) fitted for each OTU with presence as the response and biomass as predictor (glm; stats v3.6.2). OTUs occurring in fewer than five plots were excluded. For each dataset (split by sampling year) $p$-values were adjusted (Benjamini-Hochberg FDR). These OTUs were subsequently projected onto the ordination using species vectors fitted with envfit (999 permutations), allowing biomass-associated taxa to be visualised.

##### *S1.14 Permutation schemes*

Permutation schemes used for PERMANOVA and distance-based redundancy (db-RDA) analyses had to be altered depending on hypothesis, plantation type, according to sampling design. Permutation schemes were constructed in permute (v0.9-8) and matched to the level at which each factor varied. Basalt amendment and soil inoculation, randomised at plot level, were tested on plot-aggregated data by permuting whole plots within blocks with rows held fixed. Sampling year, repeated within plots, was tested on the same data with plots held in place and rows permuted within them.

For the broadleaf EcM fungal dataset, as tree-species varied within plot, hosts were permuted within plot x year combinations to test host identity. As sampling was conducted across two sampling groups, with distinct EcM hosts (two per block), we then validated plantation-wide results by testing host-identity in datasets split by sampling group. Both tests are reported (Tables S12, S18). For the broadleaf AM fungal dataset, *Prunus avium* and *Sorbus aucuparia* were sampled in non-overlapping block sets, so host identity is fully confounded with block and was not tested by PERMANOVA. Variance partition of host identity is reported as host/block predictor, as host effects cannot be distinguished from that of block,

All tests used 999 permutations and marginal (type III) terms; the same schemes were applied to PERMANOVA (adonis2; vegan v2.7.1) and to distance-based redundancy analysis of unique variance fractions (db-RDA: 999 permutations). To fit these schemes datasets had to be filtered to create even sampling efforts as needed (Broadleaf EcM, n=62; Broadleaf AM, n= 60; Sitka spruce n=64). For individual species broadleaf species n=115: Oak, n= 32; Birch, n= 30; Alder, n= 29; Aspen, n= 24; Rowan, n=32; Cherry, n= 30.

#### S2 Methods tables

***Table S 1*** Composition of broadleaf and Sitka spruce plantations at The Carbon Community Project large-scale field trial. Sampling of broadleaf plantation was divided into two sampling groups, each sampled in four blocks. Conifer (Sitka spruce) plantation was sampled in every block. In total, for each block, two ectomycorrhizal (EcM) species were sampled of both broadleaf and Sikta spruce, and one Arbuscular mycorrhizal (AM) tree species was sampled per block.

| Plantation type | Sampling group | Species | Mycorrhizal type | % of Plantation | Individuals sampled per treatment | Block Sampled |
|---|---|---|---|---|---|---|
| Broadleaf | Broadleaf group 1 | Alder (*Alnus glutinosa*) | EcM | 22% | 4 | A, B, E, G |
| | | Downy birch (*Betula pubescens*) | EcM | 35% | 4 | |

| | | | | | | |
|---|---|---|---|---|---|---|
| | | Wild cherry (*Prunus avium*) | AM | 19% | 4 | |
| | Broadleaf group 2 | Sessile oak (*Quercus petraea*) | EcM | 13% | 4 | C, D, F, H |
| | | Aspen (*Populus tremula*) | EcM | 5% | 4 | |
| | | Rowan (*Sorbus aucuparia*) | AM | 6% | 4 | |
| Conifer | Sitka spruce | Sitka spruce (*Picea sitchensis*) | EcM | 100% | 16 | A, B, C, D, E, F, G, H |

***Table S 2*** Mycorrhizal root samples taken from broadleaf and Sitka spruce plantations during the two main sampling campaigns, the first in year 2 and the second in year 4. Roots were also sampled from the tree nursery, woodland inoculum sources, the surrounding hedgerows, and the unforested pasture controls. EcM; Ectomycorrhizal fungi. AM; Arbuscular mycorrhizal fungi.

| **Source of root samples** | **Number of EcM root tips sequenced** | **Number of tips assigned to a EcM taxon** | **Number of AM roots collected** | **Number of pooled AM samples for sequencing** | **Pools found with AM taxa** |
|---|---|---|---|---|---|
| Broadleaves year 2 | 684 | 553 | 253 | 32 | 31 |
| Broadleaves year 4 | 761 | 667 | 373 | 32 | 32 |
| Tree nursery | 939 | 303 | 60 | 2 | 2 |
| Hedgerow | 336 | 261 | NA | NA | NA |
| Spruce year 2 | 768 | 640 | NA | NA | NA |
| Spruce year 4 | 770 | 596 | NA | NA | NA |
| Spruce woodland inoculum source | 62 | 31 | NA | NA | NA |
| Broadleaf woodland inoculum source | 70 | 21 | NA | NA | NA |
| Pasture control year 2 | NA | NA | 64 | 8 | 6 |
| Pasture control year 4 | NA | NA | 93 | 8 | 6 |

***Table S 3*** Mycorrhizal sampling coverage by ectomycorrhizal (EcM) or arbuscular mycorrhizal (AM) tree species. Values give the number of individual trees yielding mycorrhizal data that meets quality thresholds; trees/samples were retained for EcM analyses where at least five root tips returned successful EcM sequences; for AM analyses, where pooled root samples returned at least ten reads assigned to AM fungal taxa. Under complete sampling, 32 trees per broadleaf species and 64 Sitka spruce would be represented across sampling years 2 and 4; departures from these totals reflect sequencing failures/absence of colonisation.

| | | **By plantation** | | | **By tree species** | | | | | | |
|---|---|---|---|---|---|---|---|---|---|---|---|
| **Year of sampling** | | **Spruce** | **Broadleaf** | **Broadleaf** | **Spruce** | **Alder** | **Birch** | **Aspen** | **Oak** | **Rowan** | **Cherry** |
| | **Block** | **EcM** | **EcM** | **AM** | **EcM** | **EcM** | **EcM** | **EcM** | **EcM** | **AM** | **AM** |
| **Year 2** | A | 4 | 4 | 4 | 4 | 4 | 4 | NA | NA | NA | 4 |
| | B | 4 | 3 | 4 | 4 | 3 | 2 | NA | NA | NA | 4 |
| | E | 4 | 4 | 4 | 4 | 4 | 4 | NA | NA | NA | 4 |
| | G | 4 | 4 | 3 | 4 | 4 | 4 | NA | NA | NA | 3 |
| | C | 4 | 4 | 4 | 4 | NA | NA | 3 | 4 | 4 | NA |
| | D | 4 | 4 | 4 | 4 | NA | NA | 0 | 4 | 4 | NA |
| | F | 4 | 4 | 4 | 4 | NA | NA | 2 | 4 | 4 | NA |
| | H | 4 | 4 | 4 | 4 | NA | NA | 3 | 4 | 4 | NA |
| **Year 4** | A | 4 | 4 | 4 | 4 | 2 | 4 | NA | NA | NA | 4 |
| | B | 4 | 4 | 4 | 4 | 4 | 4 | NA | NA | NA | 4 |
| | E | 4 | 4 | 3 | 4 | 4 | 4 | NA | NA | NA | 3 |
| | G | 4 | 4 | 4 | 4 | 4 | 4 | NA | NA | NA | 4 |
| | C | 4 | 4 | 4 | 4 | NA | NA | 4 | 4 | 4 | NA |
| | D | 4 | 4 | 4 | 4 | NA | NA | 4 | 4 | 4 | NA |
| | F | 4 | 4 | 4 | 4 | NA | NA | 4 | 4 | 4 | NA |
| | H | 4 | 4 | 4 | 4 | NA | NA | 4 | 4 | 4 | NA |
| **TOTAL (n)** | | **64** | **63** | **62** | **64** | **29** | **30** | **24** | **32** | **32** | **30** |

***Table S 4*** Primer sequences.

| Primer | Sequence | Reference |
|---|---|---|
| ITS1F | 5’-CTTGGTCATTTAGAGGAAGTAA | White, 1990 |
| ITS4 | 5’-TCCTCCGCTTATTGATATGC | Gardes & Bruns, 1993 |
| NS1B1ngs | 5’-CCTNGTTGATYCTGCCAGT | Tedersoo & Anslan, 2019 |
| ITS4ngsUni | 5′-CCTSCSCTTANTDATATGC | Tedersoo & Lindahl, 2016 |
| ITS9munngs | GTACACACCGCCCGTCG | Tedersoo & Anslan, 2019 |
| ITS4ng | CGCCTSCSCTTANTDATATGC | Tedersoo & Lindahl, 2016 |

### S3 Supplementary results

#### *S3.1 Taxonomic descriptions mycorrhizal communities*

**EcM communities from sources of colonisation**

In the tree nursery, a total of 77 EcM OTUs were detected (Table S4), primarily *Hebeloma* (24%), *Thelephora* (18%) and *Peziza* (14%). Ectomycorrhizas from the surrounding hedgerows yielded 62 EcM OTUs, including *Lactarius* (22%), *Russula* (19%), *Laccaria* (10%) and *Xerocomellus* (10%). In the woodland sources of soil inoculum we identified 23 EcM OTUs, 12 of these were from the Sitka spruce woodland inoculum source, dominated by Amphinema (39%) and Thelephora (16%) and 13 from the broadleaf inoculum source, dominated by Russula (29%) and Lactarius (29%).

**AM communities from sources of colonisation**

Pasture control plots were compositionally similar to plantation AM communities. The 40 AM OTUs detected (Table S5) comprised 75% Glomeromycetes, 10% Archaeosporomycetes and 3% Paraglomeromycetes, with *Dominikia* (16%) and *Glomus* (15%) again representing the most frequent genera. In the tree nursery, the only detected AM taxon was *Rhizophagus intraradices*.

***Soil mycorrhizal communities***

We obtained 761,572 sequences from the soil eDNA sequencing after filtering. Of these, 46,257 sequences were identified as EcM fungi, assigned to 186 unique EcM OTUs and 116 unique SHs. Of the EcM fungi, 16% were Ascomycota and 84% Basidiomycota. The most common genera were *Tomentella* (12%), *Entoloma* (10%) and *Cortinarius* (9%). Among the AM fungi, 8,024 DNA sequences were assigned to 34 unique AM OTUs, nine of these assigned to a unique SH. The most common were *Archaeospora* (22%), *Glomus* (18%) and *Dominikia* (13%).

#### *S3.2 Results figures*

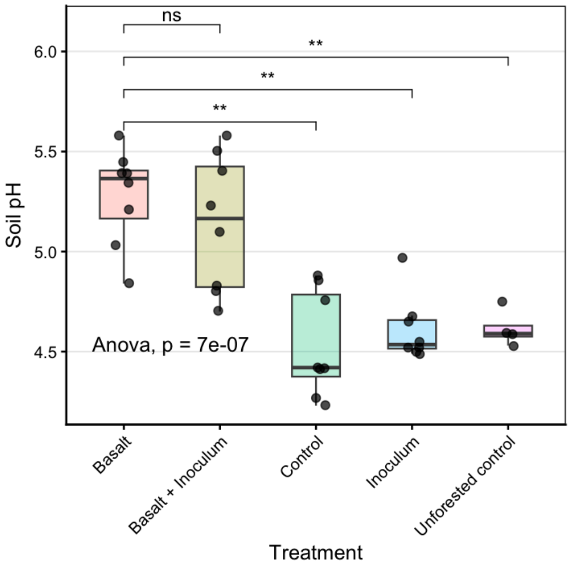

***Figure S 1*** Basalt treatment significantly (p<0.05) increased soil pH across plantation plots. Boxplots show the median and interquartile range. Overall treatment effects were assessed using ANOVA, with p-values reported; pairwise comparisons were conducted using Mann-Whitney U tests.

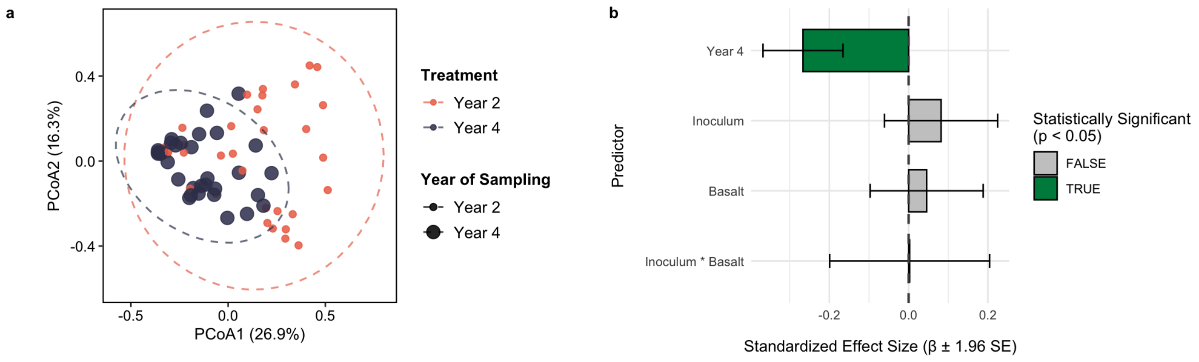

***Figure S 2*** Year of sampling predicts variation in Sitka spruce ectomycorrhizal (EcM) fungal community composition. (a) Principal coordinate analysis (PCoA) of EcM communities based on Bray-Curtis dissimilarity. The first two axes explain 26.9% and 16.3% of variation in community composition, respectively. (b) Linear mixed-effects model estimates (β coefficients ± 1.96 SE) showing the standardized effect size of predictors: Basalt treatment, soil inoculation (Inoculum) and sampling year (Year) on PCoA1. Significant predictors (p<0.05) are indicated by bar colour. PCoA1 reflects the primary gradient in EcM community structure shown in panel (a). Tree species and block were included as random intercepts.

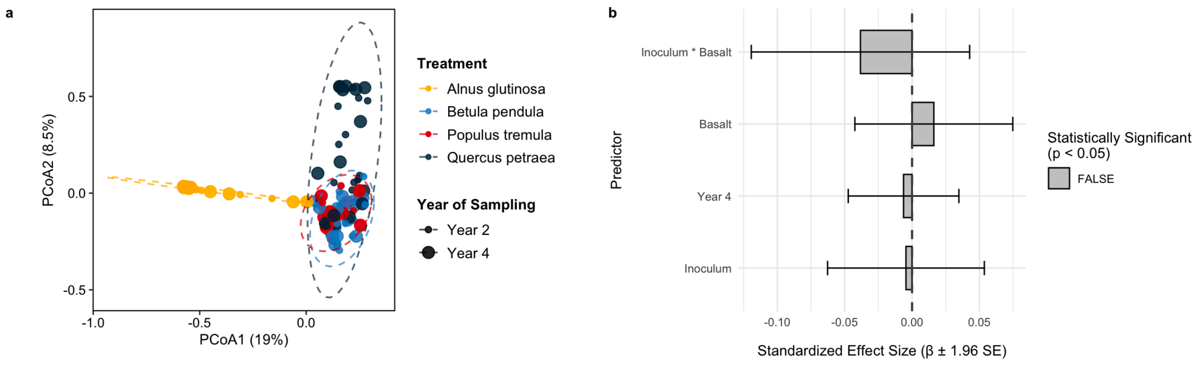

***Figure S 3*** Neither soil inoculation nor basalt treatment resulted in significant changes to ectomycorrhizal (EcM) fungal community composition in four broadleaf tree species. (a) Principal coordinate analysis (PCoA) of EcM communities based on Bray-Curtis dissimilarity. The first two axes explain 19% and 8.5% of variation in community composition, respectively. (b) Linear mixed-effects model estimates (β coefficients ± 1.96 SE) showing the standardized effect size of predictors; basalt treatment, soil inoculation (inoculum) and sampling year (Year) on PCoA1. Significant predictors ($p<0.05$) are indicated by bar colour. Tree species and block were included as random intercepts.

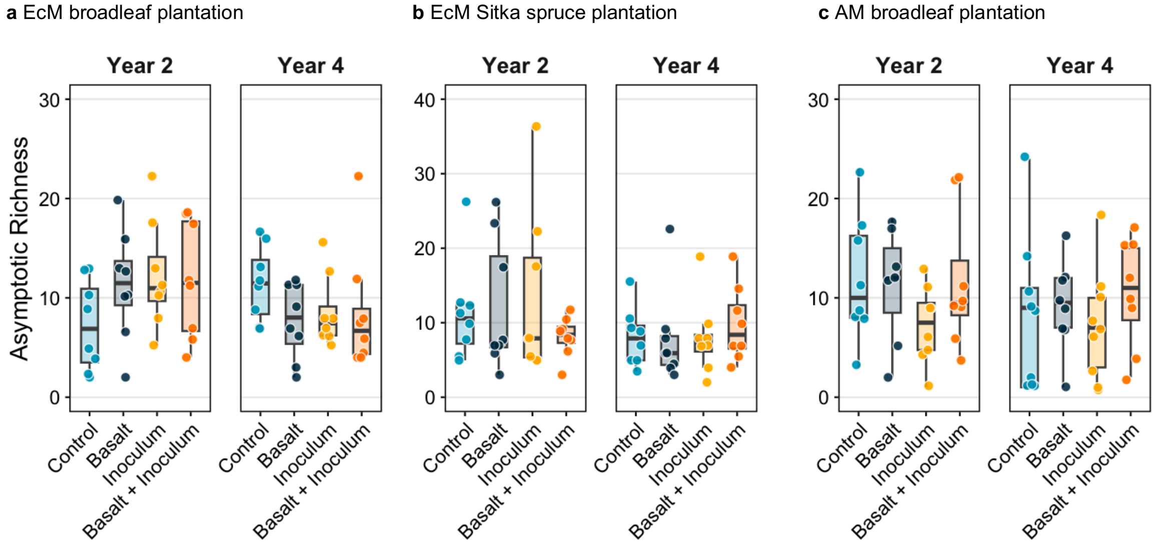

***Figure S 4*** Estimated asymptotic fungal species richness (Hill number *q=0*) calculated using iNEXT for ectomycorrhizal (EcM) fungal communities in (a) the broadleaf plantation and (b) the Sitka spruce plantation, and for arbuscular mycorrhizal (AM) fungal communities in (c) the broadleaf plantation. Species richness is shown for each carbon-capture enhancement intervention (control, basalt amendment, and soil inoculation) and sampling year. Boxplots show the median, interquartile range (IQR), and whiskers extending to the minimum and maximum values.

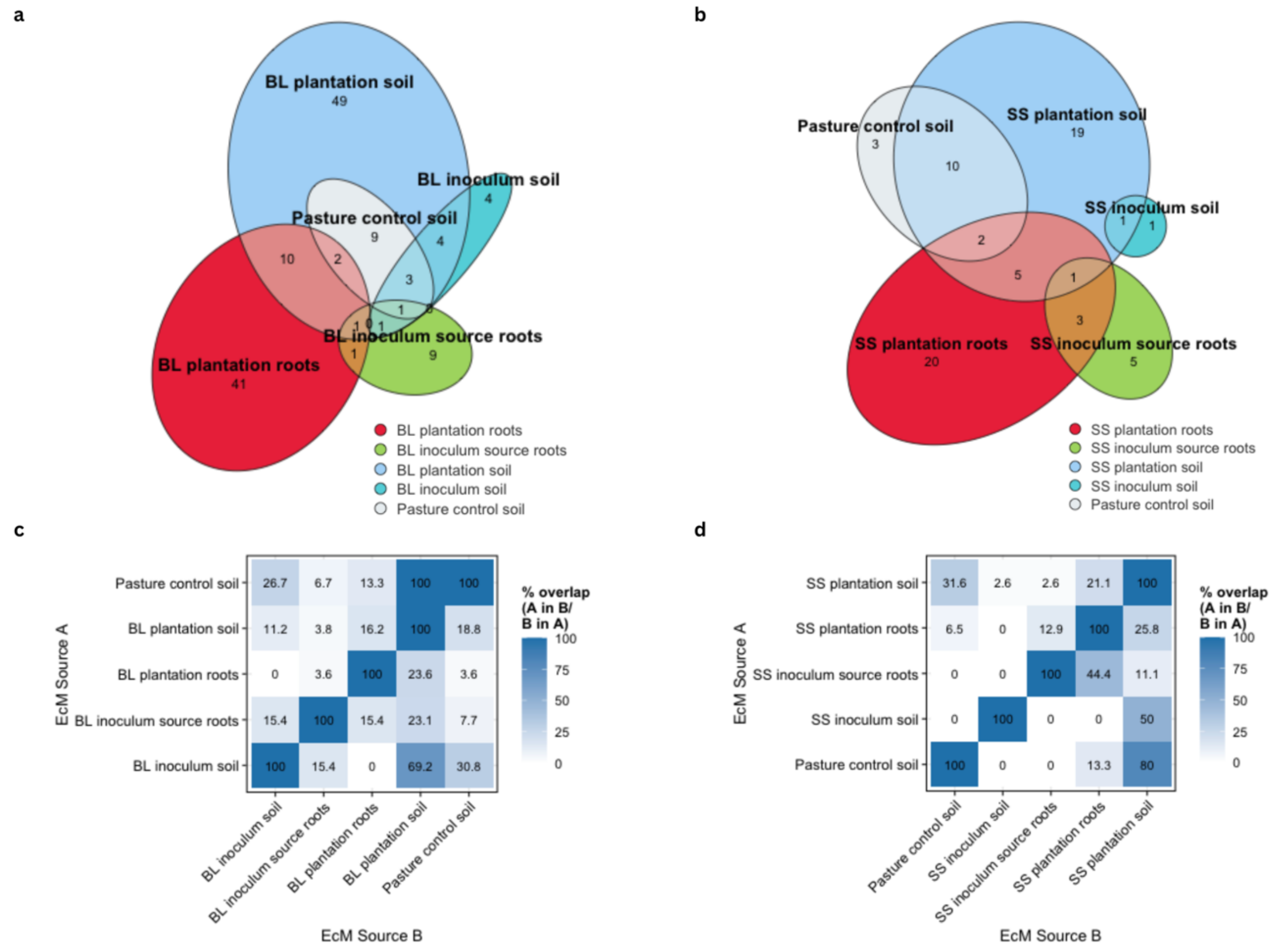

***Figure S 5*** Shared ectomycorrhizal (EcM) fungal taxa between broadleaf (BL) or Sitka spruce (SS) plantations and sources of potential EcM colonisation. Fungal taxa were identified from colonised sequenced root-tips (roots), or from soil environmental DNA samples (soil). (a, b) Euler diagrams illustrating the unique and shared EcM taxa among root-associated and soil associated sources. (c, d) Heatmaps showing asymmetric pairwise comparisons of EcM taxa shared among colonisation sources for (c) the BL plantation, (d) the SS plantation. Values represent the percentage of taxa in source A (rows) also found in source B (columns), and vice versa. (a-d) Represent only the taxa assigned to a species hypothesis (SH; UNITE v10;), to allow for root and soil comparison.

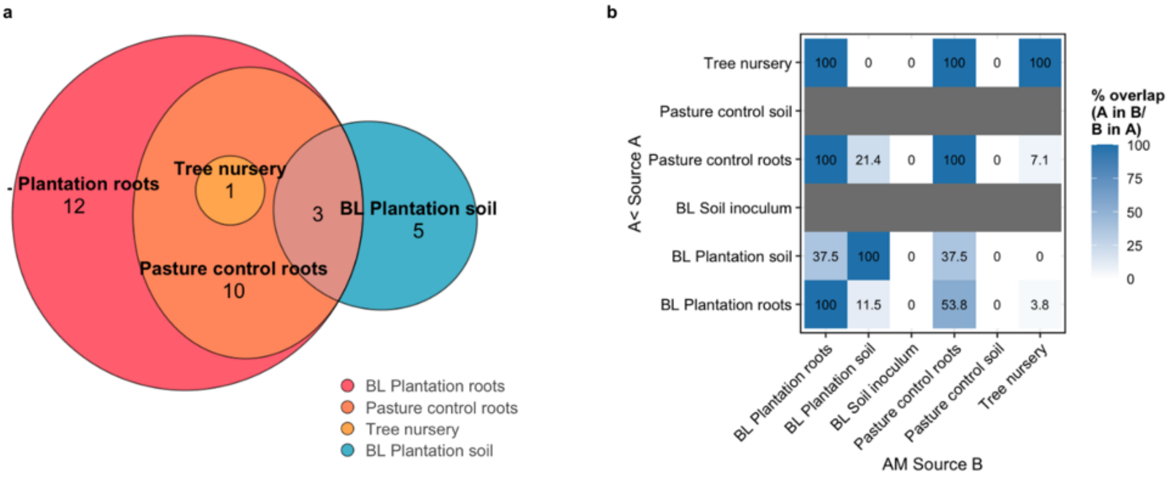

***Figure S 6*** Shared arbuscular mycorrhizal (AM) fungal taxa between a broadleaf (BL) plantation and sources of AM fungal colonisation. Taxa were identified from sequences derived from root-associated fungi (root), and from soil environmental DNA samples (soil). (a) Euler diagram illustrating the unique and shared AM taxa among root-associated and soil sources. a) Heatmap showing asymmetric pairwise comparisons of AM taxa. Values represent the percentage of taxa in source A (rows) also found in source B (columns), and vice versa. (a,b) Represent only the taxa assigned to a species hypothesis (SH; UNTIE v10), to allow for root and soil comparison.

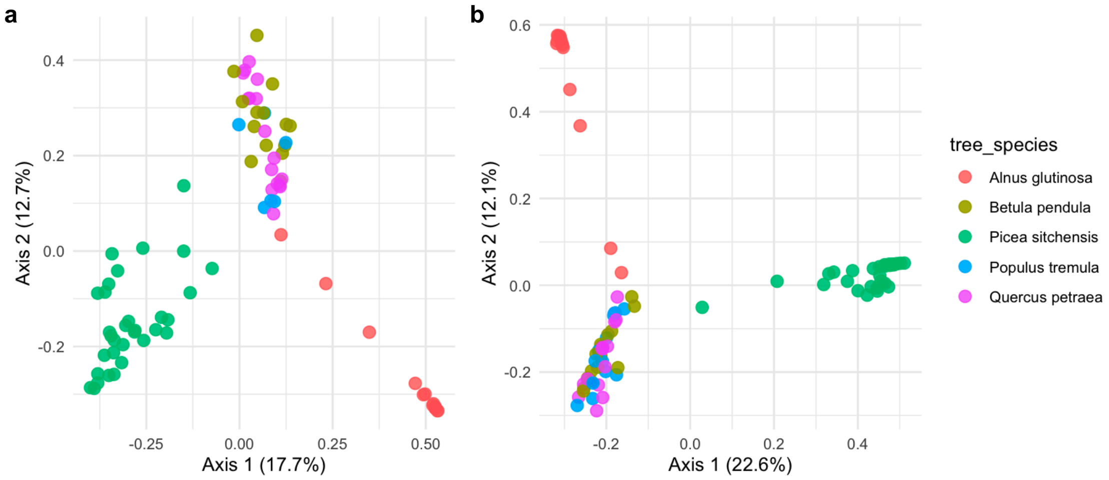

***Figure S 7*** Principal coordinate analysis (PCoA) of ectomycorrhizal (EcM) fungal communities based on Bray-Curtis dissimilarity across five EcM host tree species for (a) sampling year 2, and (b) sampling year 4.

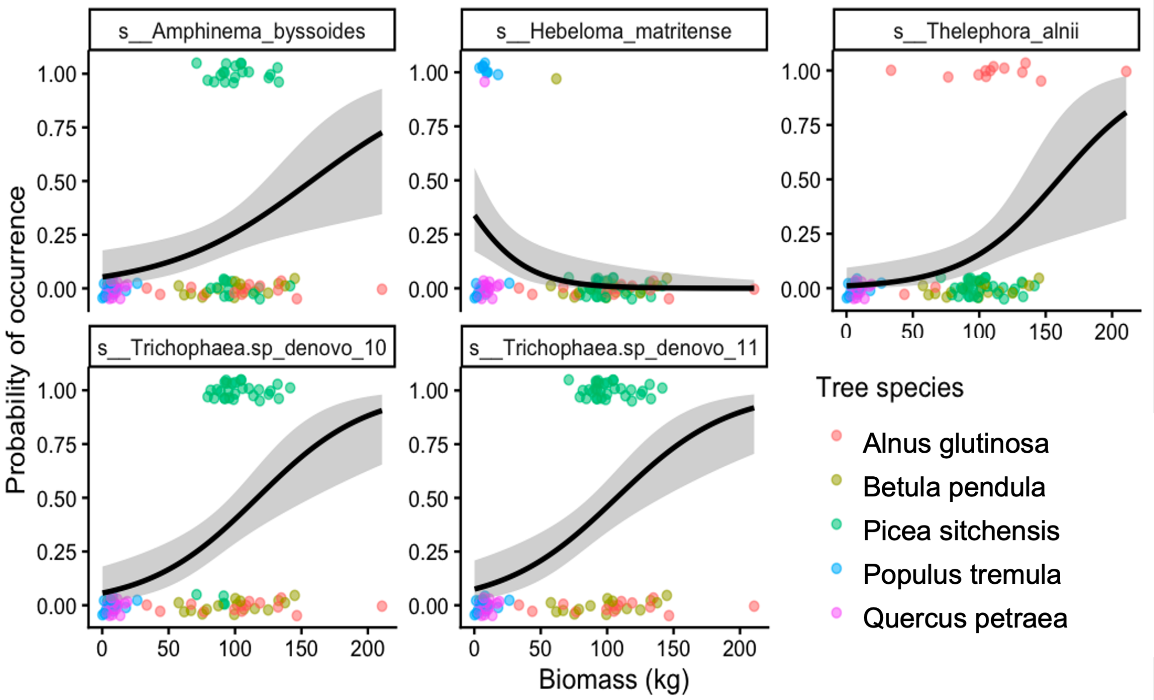

***Figure S 8*** Ectomycorrhizal taxa significantly associated with variation in tree biomass in sampling year 4. Points show presence/absence of species. Fitted lines show the overall relationship between biomass and probability of the species occurrence, based on a binomial GLM ($p<0.05$).

#### S3.3 Results tables

##### S3.3.1 Sequencing success

***Table S 5*** Number of operational taxonomic units (OTUs) detected across colonised root tips, collected from broadleaf (BL) and Sitka spruce (SS) plantations, from corresponding woodland inoculum sources, and from other potential sources of ectomycorrhizal (EcM) fungal colonisation. Values give the total number of OTUs detected and the subset classified as EcM taxa. Values are shown for sampling year 2 and 4 where appropriate.

| Source | All OTUs | EcM OTUS | % |
|---|---|---|---|
| Broadleaf plantation (Year 2) | 83 | 68 | 81.9 |
| Broadleaf plantation (Year 4) | 75 | 65 | 86.7 |
| Sitka spruce plantation (Year 2) | 60 | 44 | 73.3 |
| Sitka spruce plantation (Year 4) | 47 | 29 | 61.7 |
| Tree nursery | 331 | 77 | 23.9 |
| Hedgerow | 78 | 62 | 79.5 |
| Woodland inoculum source SS | 21 | 12 | 57.1 |
| Woodland inoculum source BL | 17 | 13 | 76.5 |

***Table S 6*** Number of operational taxonomic units (OTUs) detected across roots collected from broadleaf plantations and from pasture control plots. Values give the total number of OTUs detected and the subset classified as arbuscular mycorrhizal (AM) taxa. Values are shown for sampling year 2 and 4 where appropriate.

| Source | All OTUS | AM OTUs | % AM |
|---|---|---|---|
| Control plots | 417 | 40 | 9.6 |
| Cherry | 595 | 44 | 7.4 |
| Rowan | 754 | 62 | 8.2 |
| Broadleaf plantation (Year 2) | 466 | 58 | 12.4 |
| Broadleaf plantation (Year 4) | 756 | 57 | 7.5 |

##### *S3.3.2 Effects of treatments on mycorrhizal diversity*

***Table S 7*** Effects of basalt treatment and soil inoculation on ectomycorrhizal fungal community diversity indices of a Sitka spruce plantation. Species richness (q = 0), Shannon diversity (q = 1), and Simpson diversity (q = 2) were compared between treatment and control groups using ANOVA or Kruskal-Wallis tests as appropriate. Test statistics (test), degrees of freedom (df), and *p*-values (*p*) are reported.

| Treatment contrast | Estimated diversity metric | Test | Statistic | df | *p* |
|---|---|---|---|---|---|
| Basalt vs no basalt | Species richness (q=0) | Kruskal-Wallis | 0.157 | 1.000 | 0.692 |
| Inoculum vs no inoculum | Species richness (q=0) | Kruskal-Wallis | 0.031 | 1.000 | 0.861 |
| Basalt vs no basalt | Shannon diversity (q=1) | Kruskal-Wallis | 0.478 | 1.000 | 0.489 |
| Inoculum vs no inoculum | Shannon diversity (q=1) | Kruskal-Wallis | 0.247 | 1.000 | 0.619 |
| Basalt vs no basalt | Simpson diversity (q=2) | Kruskal-Wallis | 1.894 | 1.000 | 0.169 |
| Inoculum vs no inoculum | Simpson diversity (q=2) | Kruskal-Wallis | 0.234 | 1.000 | 0.629 |

***Table S 8*** Effects of basalt and soil inoculation treatments on ectomycorrhizal fungal community diversity indices of a broadleaf plantation. Species richness (q = 0), Shannon diversity (q = 1), and Simpson diversity (q = 2) were compared between treatment and control groups using ANOVA or Kruskal-Wallis tests as appropriate. Test statistics (test), degrees of freedom (df), and *p*-values (*p*) are reported.

| Treatment contrast | Estimated diversity metric | Test | Statistic | df | *p* |
|---|---|---|---|---|---|
| Basalt vs no basalt | Species richness (q=0) | ANOVA (F) | 0.003 | 1, 62 | 0.959 |
| Inoculum vs no inoculum | Species richness (q=0) | Kruskal-Wallis | 0.069 | 1 | 0.793 |
| Basalt vs no basalt | Shannon diversity (q=1) | Kruskal-Wallis | 0.058 | 1 | 0.783 |
| Inoculum vs no inoculum | Shannon diversity (q=1) | Kruskal-Wallis | 0.883 | 1 | 0.347 |
| Basalt vs no basalt | Simpson diversity (q=2) | Kruskal-Wallis | 0.122 | 1 | 0.727 |
| Inoculum vs no inoculum | Simpson diversity (q=2) | Kruskal-Wallis | 1.412 | 1 | 0.235 |

***Table S 9*** Effects of basalt and inoculum treatments on arbuscular mycorrhizal fungal community diversity indices of a broadleaf plantation. Species richness (q = 0), Shannon diversity (q = 1), and Simpson diversity (q = 2) were compared between treatment and control groups using ANOVA or Kruskal-Wallis tests as appropriate. Test statistics (test), degrees of freedom (df), and *p*-values (*p*) are reported.

| Treatment contrast | Estimated diversity metric | Test | Statistic | df | *p* |
|---|---|---|---|---|---|
| Inoculum vs no inoculum | Species richness (q=0) | ANOVA (F) | 0.386 | 1,63 | 0.537 |
| Basalt vs no basalt | Species richness (q=0) | Kruskal-Wallis | 3.138 | 1 | 0.076 |
| Inoculum vs no inoculum | Shannon diversity (q=1) | ANOVA (F) | 1.145 | 1,63 | 0.289 |
| Basalt vs no basalt | Shannon diversity (q=1) | Kruskal-Wallis | 2.167 | 1 | 0.141 |
| Inoculum vs no inoculum | Simpson diversity (q=2) | ANOVA (F) | 1.263 | 1,63 | 0.265 |
| Basalt vs no basalt | Simpson diversity (q=2) | Kruskal-Wallis | 2.128 | 1 | 0.145 |

##### *S3.3.3 Effects of treatments on mycorrhizal fungal community composition by PERMANOVA*

***Table S 10*** PERMANOVA test of 999 permutations tested on Bray-Curtis dissimilarity of ectomycorrhizal (EcM) fungal community composition of a Sitka spruce plantation. Marginal tests of each term are given from an additive model; the inoculum × basalt interaction was tested separately and not supported. Permutations were restricted according to the experimental design; inoculum and basalt were applied to plots nested within eight blocks, and each plot was sampled in both sampling year (year 2 and year 4). Plots were permuted freely within blocks for inoculum and basalt, and sampling years permuted within plots for year. $R^2$ gives the proportion of total variation in composition attributable to each term. Degrees of freedom (df), sum of squares (SS), F value (F), $R^2$ and p-values (p) are reported.

| Model | Predictor | df | SS | $R^2$ | F | *p* |
|---|---|---|---|---|---|---|
| Full model | Inoculum x Basalt | 1 | 0.164 | 0.012 | 0.855 | 0.502 |
| | Residual | 59 | | | | |
| Additive model | Inoculum | 1 | 0.422 | 0.032 | 2.213 | 0.009 |
| | Basalt | 1 | 0.162 | 0.012 | 0.850 | 0.488 |
| | Year of sampling | 1 | 1.362 | 0.102 | 7.137 | 0.001 |
| | Residual | 60 | 11.450 | 0.855 | | |
| | Total | 63 | 13.396 | 1.000 | | |
| | Year x Inoculum | 1 | 0.103 | 0.008 | 0.536 | 0.800 |

***Table S 11*** PERMANOVA test of 999 permutations tested on Bray-Curtis dissimilarity of ectomycorrhizal fungal community composition of a broadleaf plantation. Marginal tests of each term are given from an additive model; the inoculum × basalt interaction was tested separately and not supported. Permutations were restricted according to the experimental design; inoculum and basalt were applied to plots nested within eight blocks, and each plot was sampled in both sampling years (year 2 and year 4). Plots were permuted freely within blocks for inoculum and basalt, and sampling years permuted within plots for year. $R^2$ gives the proportion of total variation in composition attributable to each term. Degrees of freedom (df), sum of squares (SS), F value (F), $R^2$ and p-values (p) are reported.

| Model | Predictor | df | SS | $R^2$ | F | *p* |
|---|---|---|---|---|---|---|
| Full model | Inoculum x Basalt | 1 | 0.333 | 0.014 | 0.882 | 0.416 |
| | Residual | 57 | | | | |
| Additive model | Inoculum | 1 | 0.302 | 0.013 | 0.802 | 0.528 |
| | Basalt | 1 | 0.462 | 0.020 | 1.224 | 0.141 |
| | Year of sampling | 1 | 0.595 | 0.026 | 1.577 | 0.001 |
| | Residual | 58 | 21.881 | 0.942 | | |
| | Total | 61 | 23.237 | 1.000 | | |

***Table S 12*** PERMANOVA test of 999 permutations tested on Bray-Curtis dissimilarity of ectomycorrhizal fungal community composition of four broadleaf tree species within a plantation. Permutations were restricted according to the experimental design; inoculum and basalt were applied to plots nested within eight blocks, and each plot was sampled in both year 2 and year 4. Plots were permuted freely within blocks for inoculum and basalt, and sampling years permuted within plots for year. Host identity varied within plots; permutations within plot × year combinations were used to test host identity. Because each term was tested under a different scheme, residual and total values are given as subscripts alongside the corresponding predictor. Marginal tests ("by = margin") were performed. $R^2$ gives the proportion of total variation in composition attributable to each term. Degrees of freedom (df), sum of squares (SS), F value (F), $R^2$ and p-values (p) are reported.

| **Predictor** $_{\text{residual, total}}$ | **df** | **SS** | $R^2$ | **F** | **p** |
|---|---|---|---|---|---|
| Inoculum $_{58,61}$ | 1 | 0.394 | 0.008 | 1.189 | 0.235 |
| Basalt $_{58,61}$ | 1 | 0.442 | 0.009 | 1.332 | 0.167 |
| Year of sampling $_{58,61}$ | 1 | 0.597 | 0.012 | 1.799 | 0.001 |
| Host identity $_{108,114}$ | 3 | 12.473 | 0.250 | 12.530 | 0.001 |
| Host identity (Oak vs Aspen) $_{51,55}$ | 1 | 1.578 | 0.070 | 4.100 | 0.001 |
| Host identity (Alder vs birch) $_{54,58}$ | 1 | 6.725 | 0.295 | 23.651 | 0.001 |

***Table S 13*** PERMANOVA test of 999 permutations tested on Jaccard dissimilarity of arbuscular mycorrhizal fungal community composition of two broadleaf tree species within a plantation. Marginal tests of each term are given from an additive model; the inoculum ×

basalt interaction was tested separately and not supported. Permutations were restricted according to the experimental design; inoculum and basalt were applied to plots nested within eight blocks, and each plit was sampled in both sampling years (year 2 and year 4). Plots were permuted freely within blocks for inoculum and basalt, and sampling years permuted within plots for year. Marginal tests ("by = margin") were performed. Degrees of freedom (df), sum of squares (SS), F value (F), $R^2$ and *p*-values (*p*) are reported.

| **Model** | **Predictor** | **df** | **SS** | **$R^2$** | **F** | ***p*** |
|---|---|---|---|---|---|---|
| Full model | Inoculum x Basalt | 1 | 0.293 | 0.015 | 0.916 | 0.543 |
| | Residual | 57 | | | | |
| Additive model | Inoculum | 1 | 0.412 | 0.022 | 1.291 | 0.168 |
| | Basalt | 1 | 0.369 | 0.019 | 1.155 | 0.251 |
| | Year of sampling | 1 | 0.461 | 0.024 | 1.443 | 0.031 |
| | Residual | 56 | 17.870 | 0.936 | | |
| | Total | 59 | 19.098 | 1.000 | | |

##### *S3.3.4 Permutation test for homogeneity of multivariate dispersions*

**Table S 14** PERMDISP test of 999 permutations tested on Bray-Curtis dissimilarity of ectomycorrhizal (EcM) fungal community composition of a Sitka spruce plantation. Degrees of freedom (df), sum of squares (SS), F value (F), and p-values (*p*) are reported.

| | | **df** | **SS** | **MS** | **F** | ***p*** |
|---|---|---|---|---|---|---|
| Inoculum | Groups | 1 | 0.020 | 0.020 | 0.747 | 0.202 |
| | Residuals | 62 | 1.632 | 0.026 | | |
| Basalt | Groups | 1 | 0.014 | 0.014 | 0.550 | 0.263 |
| | Residuals | 62 | 1.633 | 0.026 | | |
| Year | Groups | 1 | 0.421 | 0.421 | 26.451 | 0.001 |
| | Residuals | 62 | 0.986 | 0.016 | | |

***Table S 15*** PERMDISP test of 999 permutations tested on Bray-Curtis dissimilarity of ectomycorrhizal (EcM) fungal community composition of a broadleaf plantation. Test of homogeneity of multivariate dispersion for each factor. Degrees of freedom (df), sum of squares (SS), mean square (MS), F value (F), and *p*-values (*p*) are reported.

| | | **df** | **SS** | **MS** | **F** | ***p*** |
|---|---|---|---|---|---|---|
| Inoculum | Groups | 1 | 0.000 | 0.000 | 0.001 | 0.974 |
| | Residuals | 60 | 0.389 | 0.006 | | |
| Basalt | Groups | 1 | 0.001 | 0.001 | 0.092 | 0.783 |
| | Residuals | 60 | 0.367 | 0.006 | | |

| | | | | | | |
|---|---|---|---|---|---|---|
| Year | Groups | 1 | 0.000 | 0.000 | 0.003 | 0.952 |
| | Residuals | 60 | 0.415 | 0.007 | | |
| Host | Groups | 3 | 1.699 | 0.566 | 22.367 | 0.001 |
| | Residuals | 111 | 2.810 | 0.025 | | |

***Table S 16*** PERMDISP test of 999 permutations tested on Bray-Curtis dissimilarity of ectomycorrhizal (AM) fungal community composition of a broadleaf plantation. Test of homogeneity of multivariate dispersion for each factor. Degrees of freedom (df), sum of squares (SS), mean square (MS), F value (F), and *p*-values (*p*) are reported.

| **Factor** | | **df** | **SS** | **MS** | **F** | ***p*** |
|---|---|---|---|---|---|---|
| Inoculum | Groups | 1 | 0.002 | 0.002 | 0.277 | 0.561 |
| | Residuals | 58 | 0.520 | 0.009 | | |
| Basalt | Groups | 1 | 0.025 | 0.025 | 2.998 | 0.056 |
| | Residuals | 58 | 0.486 | 0.008 | | |
| Year | Groups | 1 | 0.011 | 0.011 | 1.216 | 0.223 |
| | Residuals | 58 | 0.511 | 0.009 | | |

***S3.3.5 Variance partitioning of mycorrhizal communities***

***Table S 17*** Results of distance-based redundancy analyses (db-RDA) used to partition variation in Sitka spruce ectomycorrhizal (ECM) fungal community composition explained by either soil inoculation (inoculum), basalt amendment (basalt) or year of sampling (year 2 or year 4; n= 64). Each model tested the unique contribution of one predictor while conditioning on the remaining predictors. Significance was assessed using permutation tests, and permutation were restricted according to experimental design: inoculum and basalt were applied to plots nested within eight blocks, and each plot was sampled in both year 2 and year 4. Plots permuted freely within blocks for inoculum and basalt, and sampling years permuted within plots for year. Degrees of freedom (df), F value (F), and *p*-values (*p*) are reported.

| **Predictor** | **Df** | **F** | ***p*** |
|---|---|---|---|
| Inoculum | 1 | 2.213 | 0.015 |
| Basalt | 1 | 0.850 | 0.432 |
| Year of sampling | 1 | 7.137 | <0.001 |
| Residual | 60 | | |

***Table S 18*** Results of distance-based redundancy analyses (db-RDA) used to partition variation in broadleaf ectomycorrhizal (ECM) fungal community composition explained by either soil inoculation (inoculum), basalt amendment (basalt), year of sampling (year 2 or year 4) or host tree species identity. Each model tested the unique contribution of one predictor while conditioning on the remaining predictors. Significance was assessed using permutation tests, and permutation were restricted according to experimental design; inoculum and basalt were applied to plots nested within eight blocks, and each plot was sampled in both year 2 and year 4. Plots permuted freely within blocks for inoculum and basalt, and sampling years permuted within plots for year. Host identity varied within plots, permutations within plot × year combinations were used to test host identity. Because each term was tested under a different scheme, residual values are given as subscripts alongside the corresponding predictor. Degrees of freedom (df), F value (F), and *p*-values (*p*) are reported.

| **Predictor** $_{residual}$ | **df** | **F** | ***p*** |
|---|---|---|---|
| Inoculum $_{58}$ | 1 | 0.839 | 0.307 |
| Basalt $_{58}$ | 1 | 1.164 | 0.068 |
| Year of sampling $_{58}$ | 1 | 1.6492 | 0.003 |
| Host identity $_{108}$ | 3 | 12.530 | 0.001 |
| Host identity (Oak vs Aspen) $_{51}$ | 1 | 4.100 | 0.001 |
| Host identity (Alder vs Birch) $_{54}$ | 1 | 23.651 | 0.001 |

***Table S 19*** Results of distance-based redundancy analyses (db-RDA) used to partition variation in broadleaf arbuscular mycorrhizal (AM) fungal community composition explained by either soil inoculation (inoculum), basalt amendment (basalt), year of sampling or host tree species identity/block. Each model tested the unique contribution of one predictor while conditioning on the remaining predictors. Significance was assessed using permutation tests, with permutation restricted according to experimental design; inoculum and basalt were applied to plots nested within eight blocks, and each plot was sampled in both year 2 and year 4. Plots permuted freely within blocks for inoculum and basalt, and sampling years permuted within plots for year. Variance attributable to host identity and block cannot be partitioned as AM tree species host covaried with block. Degrees of freedom (df), F value (F), and *p*-values (*p*) are reported.

| **Predictor** | **df** | **F** | ***p*** |
|---|---|---|---|
| Inoculum | 1 | 1.306 | 0.142 |

| | | | |
|---|---|---|---|
| Basalt | 1 | 1.124 | 0.283 |
| Year of sampling | 1 | 1.594 | 0.018 |
| Host identity/Block* | 1 | 2.961 | 0.001 |
| Residual | 55 | | |

* 1 Host and block cannot be partitioned.

##### *S3.3.6 Indicator species analysis*

***Table S 20*** Results of indicator species analysis for ectomycorrhizal fungi, identifying taxa associated with soil inoculation (inoculum vs no inoculum). The r.g statistic represents the strength and direction of the relationship between species occurrence and treatment. *p* values are based on 999 permutations.

| Ectomycorrhizal fungi | Group | Statistic | Method | *p* |
|---|---|---|---|---|
| *Hebeloma hiemale* (SH1375983.10FU) | No inoculum | 0.303 | r.g | 0.008 |
| *Trichophaea sp.2* (SH1093534.10FU) | Inoculum | 0.266 | r.g | 0.042 |

***Table S 21*** Results of indicator species analysis for arbuscular mycorrhizal (AM) fungi, identifying taxa associated with basalt treatment (basalt vs no basalt). The r.g statistic represents the strength and direction of the relationship between species occurrence and treatment. *p* values are based on 999 permutations**.**

| Arbuscular mycorrhizal fungi | Group | Statistic | Statistic | *p* |
|---|---|---|---|---|
| *Glomeraceae sp.1* *(OTU: bcb020928fd61ab8def23fedecb31574)* | Basalt | 0.363 | r.g | 0.008 |
| *Claroideoglomus sp.1* *(OTU: d82fd15859ea2062925a318a8b2ae31a)* | Basalt | 0.355 | r.g | 0.008 |

##### *S3.3.7 Colonisation assessment*

***Table S 22*** Results of a beta-binomial generalized linear model examining the effects of inoculum treatment, basalt amendment, and host tree species identity on the proportion of root colonised by arbuscular mycorrhizal (AM) fungi in *Prunus avium* (Pa) and *Sorbus acuparia* (Sa) roots from a broadleaf plantations. Root colonisation was calculated as the proportion of assessed root units exhibiting colonisation relative to the total number of assessed root units (colonised + non-colonised) for each seedling. Data were aggregated at the individual tree level, and the total number of assessed root units per tree was included as model weights. Detailed methods for colonisation assessment can be found in S3. Beta binomial was fitted due to overdispersion (19.65827) of the binomial model.

| | Predictor | Estimate | Standard Error | z value | *p* |
|---|---|---|---|---|---|
| Model 1 | Intercept | -3.451 | 0.224 | -15.400 | <0.001 |
| | Inoculum | 0.127 | 0.261 | 0.488 | 0.625 |
| | Basalt | 0.123 | 0.261 | 0.474 | 0.636 |
| | Tree species (Sa vs Pa) | 0.520 | 0.190 | 2.745 | 0.006 |
| | Inoculum x Basalt | -0.224 | 0.367 | -0.609 | 0.542 |
| Model 2 | Intercept | -3.235 | 0.203 | -15.899 | <0.001 |
| | Inoculum | 0.035 | 0.178 | 0.199 | 0.842 |
| | Basalt | -0.356 | 0.286 | -1.246 | 0.213 |
| | Tree species (Sa vs Pa) | 0.227 | 0.253 | 0.894 | 0.371 |
| | Basalt x Tree species | 0.605 | 0.367 | 1.650 | 0.099 |

***S3.3.8 Modelling effect of treatments on mycorrhizal fungal community composition***

***Table S 23*** Linear mixed-effects model estimates for the effects of basalt treatment (+ Basalt), soil inoculation (+ Inoculum), and sampling year (+Year) on ectomycorrhizal fungal community composition within a broadleaf plantation. Community structure was constructed using principal coordinates analysis (PCoA) based on Bray-Curtis dissimilarity. Model included tree species identity as a random intercept. Fixed-effect estimates are shown as regression coefficients (β), standard errors (SE), t-values (t), *p* values (*p*), and degrees of freedom (df). Model fit is reported using AIC and $R^2$ values.

| Model term | β | SE | t | *p* | df |
|---|---|---|---|---|---|
| Intercept | 0.012 | 0.160 | 0.078 | 0.943 | 3 |
| +Inoculum | -0.005 | 0.030 | -0.153 | 0.879 | 103 |
| +Basalt | 0.016 | 0.030 | 0.541 | 0.590 | 103 |
| +Year (Year 4) | -0.006 | 0.021 | -0.301 | 0.764 | 103 |
| Inoculum x Basalt | -0.038 | 0.041 | -0.926 | 0.357 | 103 |
| Model | PCoA1 ~ Inoculum x Basalt + year + (1 \| tree species): Conditional $R^2$: 0.895, Marginal $R^2$ = 0.002 | | | | |

* 1 Block and plot were initially included as a random effect to account for the experimental design; however, the model exhibited a singular fit, indicating negligible variance attributable to block. Consequently, block was excluded from the final model.

##### *S3.3.9 Ectomycorrhizal community composition and tree biomass*

***Table S 24*** Linear mixed-effects model estimates for the effects of ectomycorrhizal fungal community composition (+ principal coordinates analysis; PCoA1 and PCoA2), plantation type (+ Plantation type), basalt treatment (+ Basalt) and soil inoculation (+ Inoculum) on plantation biomass (kg) for sampling year 2 and 4. Biomass was square-root transformed (sqrt). Community structure was constructed using PCoA based on Bray-Curtis dissimilarity. Models were fitted with block and plot as a random intercept. Plantation type described Sitka spruce plantation or broadleaf plantation. Fixed-effect estimates are shown as regression coefficients (β), standard errors (SE), t-values (t), *p* values (*p*), and degrees of freedom (df). Model fit is reported using AIC and $R^2$ values.

| Model term | | | Intercept | + PCoA1 | + PCoA2 | + Plantation type (Sitka spruce vs broadleaf) | + Inoculum | + Basalt | B x I | AIC | $R^2$ |
|---|---|---|---|---|---|---|---|---|---|---|---|
| Model | A | β | 2.141 | 1.620 | 0.700 | 2.945 | 0.132 | 0.041 | -0.246 | 234.5353 | Conditional $R^2$: 0.707 Marginal $R^2$ = 0.440 |
| | | SE | 0.435 | 1.048 | 0.793 | 0.690 | 0.262 | 0.259 | 0.360 | | |
| | | t | 4.927 | 1.546 | 0.883 | 4.265 | 0.504 | 0.157 | -0.684 | | |
| | | *p* | 0.000 | 0.127 | 0.380 | <0.001 | 0.616 | 0.876 | 0.496 | | |
| | | df | 29.244 | 71.246 | 70.387 | 70.899 | 68.869 | 68.951 | 68.854 | | |
| | B | β | 4.848 | -7.001 | 2.804 | 8.120 | 0.132 | 0.157 | -0.624 | 419.0483 | Conditional $R^2$: 0.657 Marginal $R^2$: 0.307 |
| | | SE | 1.208 | 3.660 | 1.134 | 2.449 | 0.621 | 0.629 | 0.894 | | |
| | | t | 4.012 | -1.913 | 2.472 | 3.316 | 0.212 | 0.250 | -0.699 | | |
| | | *p* | 0.000 | 0.059 | 0.016 | <0.001 | 0.833 | 0.803 | 0.487 | | |
| | | df | 32.863 | 78.632 | 80.731 | 78.647 | 77.839 | 77.827 | 77.888 | | |
| | A | Plantation biomass Year 2 (kg)) ~ PCoA1 + PCoA2 + Basalt x Inoculum + Plantation type + (1\|block) + (1\|plot) | | | | | | | | | |
| | B | Plantation biomass Year 4 (kg)) ~ PCoA1 + PCoA2 + Basalt x Inoculum + Plantation type + (1\|block) + (1\| plot) | | | | | | | | | |

***Table S 25*** Linear mixed-effects model estimates for the effects of ectomycorrhizal fungal (EcM) community composition (+ principal coordinates analysis; PCoA1 and PCoA2), within-host tree species EcM community composition (+ PCoA1 within, + PCoA2 within), basalt treatment (+ Basalt), soil inoculation (+ Inoculum), and host tree species identity (+ Tree species; Birch/Sitka/Aspen/Oak/Alder) on plantation biomass (kg) in year 4 (2024). Biomass was square-root transformed (sqrt). Community structure was constructed using PCoA based on Bray-Curtis dissimilarity and centered for each tree species to represent for within species PCoAs. Model selection was based on AIC. Estimates are shown as regression (β), the unstandardized coefficient on sqrt-transformed biomass scale. Standard error (SE); residual degrees of freedom (df); t-values (t); p-values (p).

***Models***

| **Models** | **EcM** | | | | | **Tree species** | | | | | **Tree species + EcM within** | | | | |
|---|---|---|---|---|---|---|---|---|---|---|---|---|---|---|---|
| **Fixed effects** | **β** | **SE** | **df** | **t** | ***p*** | **β** | **SE** | **df** | **t** | ***p*** | **β** | **SE** | **df** | **t** | ***p*** |
| Alder (intercept) | 7.613 | 0.811 | 13.434 | 9.384 | 0.000 | 10.058 | 0.408 | 92 | 24.635 | <0.001 | 10.042 | 0.396 | 92 | 25.329 | <0.001 |
| Birch | | | | | | -0.207 | 0.454 | 92 | -0.457 | 0.649 | -0.199 | 0.441 | 92 | -0.452 | 0.652 |
| Sitka spruce | | | | | | -0.052 | 0.398 | 92 | -0.131 | 0.896 | -0.044 | 0.386 | 92 | -0.113 | 0.910 |
| Apsen | | | | | | -7.596 | 0.469 | 92 | -16.188 | <0.001 | -7.586 | 0.455 | 92 | -16.660 | <0.001 |
| Oak | | | | | | -7.110 | 0.454 | 92 | -15.660 | <0.001 | -7.102 | 0.441 | 92 | -16.121 | <0.001 |
| Inoculum | 0.123 | 0.642 | 83.690 | 0.192 | 0.848 | 0.110 | 0.366 | 92 | 0.300 | 0.765 | 0.201 | 0.358 | 92 | 0.563 | 0.575 |
| Basalt | 0.023 | 0.648 | 83.660 | 0.035 | 0.972 | 0.081 | 0.370 | 92 | 0.219 | 0.827 | 0.121 | 0.360 | 92 | 0.336 | 0.738 |
| Inoculum x basalt | -0.083 | 0.907 | 83.696 | -0.092 | 0.927 | -0.215 | 0.518 | 92 | -0.415 | 0.679 | -0.444 | 0.515 | 92 | -0.862 | 0.391 |
| PCoA1 | 4.914 | 0.714 | 83.711 | 6.886 | 0.000 | | | | | | | | | | |
| PCoA2 | 4.369 | 1.077 | 87.538 | 4.058 | 0.000 | | | | | | | | | | |
| PCoA1 within | | | | | | | | | | | -2.861 | 2.078 | 92 | -1.377 | 0.172 |
| PCoA2 within | | | | | | | | | | | 2.953 | 1.471 | 92 | 2.008 | 0.048 |

| | |
|---|---|
| **EcM model** | Plantation biomass (year 4) ~ PCoA1 + PCoA2 + Inoculum x Basalt + (1 \| block): AIC 437.6723, Conditional $R^2$=0.600, Marginal $R^2$ = 0.298 |
| **Tree species model** | Plantation biomass (year 4) ~ Tree species + Inoculum x Basalt + (1 \| block): AIC: 320.603, Conditional $R^2$= 0, Marginal $R^2$ = 0.884 |
| **Tree species + EcM within model** | Plantation biomass (year 4) ~ Tree species + PCoA1 within + PCoA1 within + Inoculum x Basalt + (1 \| block): AIC: 319.0546, Conditional $R^2$= 0, Marginal $R^2$= 0.884 |

*** 2**Plot was initially included as a random effect to account for the experimental design; however, the model exhibited a singular fit, indicating negligible variance attributable to plot.

Consequently, plot was excluded from the final model. Tree species effects are relative to Alder, *Alnus glutinosa* (reference category).

***Table S 26*** Model comparison, with anova() stats (v3.6.2).

| Model | npar | AIC | BIC | logLik | -2*log(L) C | Chisq | df | p |
|---|---|---|---|---|---|---|---|---|
| Tree species model | 10 | 320.6 | 345.82 | -150.3 | 300.6 | | | |
| Tree species + EcM within model | 12 | 319.05 | 349.32 | -147.53 | 295.06 | 5.549 | 2 | 0.0624 |

***Table S 27*** Ectomycorrhizal (EcM) taxon significantly associated with host-dependent variation in plantation tree biomass (kg). Plot-level OTU abundance data were converted to presence-absence, and a separate binomial logistic regression was fitted for each OTU with biomass as the predictor. OTUs occurring in fewer than five plots were excluded. p-values were adjusted for multiple testing using the Benjamini-Hochberg false discovery rate (FDR), and OTUs with adjusted p ≤ 0.05 were considered significantly associated with biomass.

| EcM taxon | Estimate | SE | statistic | p | p adj |
|---|---|---|---|---|---|
| *Thelephora alnii* | 0.0285 | 0.0103 | 2.7529 | 0.0059 | 0.0355 |
| *Amphinema byssoides* | 0.0182 | 0.0067 | 2.7386 | 0.0062 | 0.0355 |
| *Hebeloma matritense* | -0.0397 | 0.0152 | -2.6172 | 0.0089 | 0.0408 |
| *Trichophaea*. sp. denovo 10 | 0.024 | 0.007 | 3.62o | <0.001 | 0.003 |
| *Trichophaea.* sp. denovo 11 | 0.0234 | 0.0062 | 3.7781 | 0.0002 | 0.0034 |

#### S4 Arbuscular mycorrhizal colonisation assessment

Stained root sections were scanned using an Epredia 3DHISTECH Pannoramic MIDI II Rx brightfield digital slide scanner with a 37× objective (0.27 µm pixel−1). Digital images were analysed in QuPath (v0.6.0) (Bankhead et al., 2017). Individual roots were manually delineated using the annotation tool and assigned the root classification. A custom QuPath script was then used to randomly distribute 55 assessment units within each annotated root section. Each assessment unit consisted of six neighbouring rectangular assessment regions (50 × 20 µm), each with a central classification point (Fig. S9). Each assessment region was examined for the presence or absence of arbuscular mycorrhizal (AM) colonisation, identified by aseptate fungal hyphae and associated structures, including arbuscules. The central classification point was then assigned to a predefined annotation category according to the scoring criteria described in Supplementary Table S28.

Assessment regions were excluded if ≥50% of the assessment region fell outside the root tissue, intersected vascular tissue, or could not be reliably assessed because of poor focus, excessive staining, debris, or other image artefacts. These assessment units were classified as unreadable and excluded from subsequent analyses.

Classification data were exported from QuPath for each root section. For statistical analyses, data from up to 12 root sections were pooled at the seedling level. The proportion of AM colonisation for each seedling was calculated as the number of classification points scored as colonised divided by the total number of readable classification points assessed.

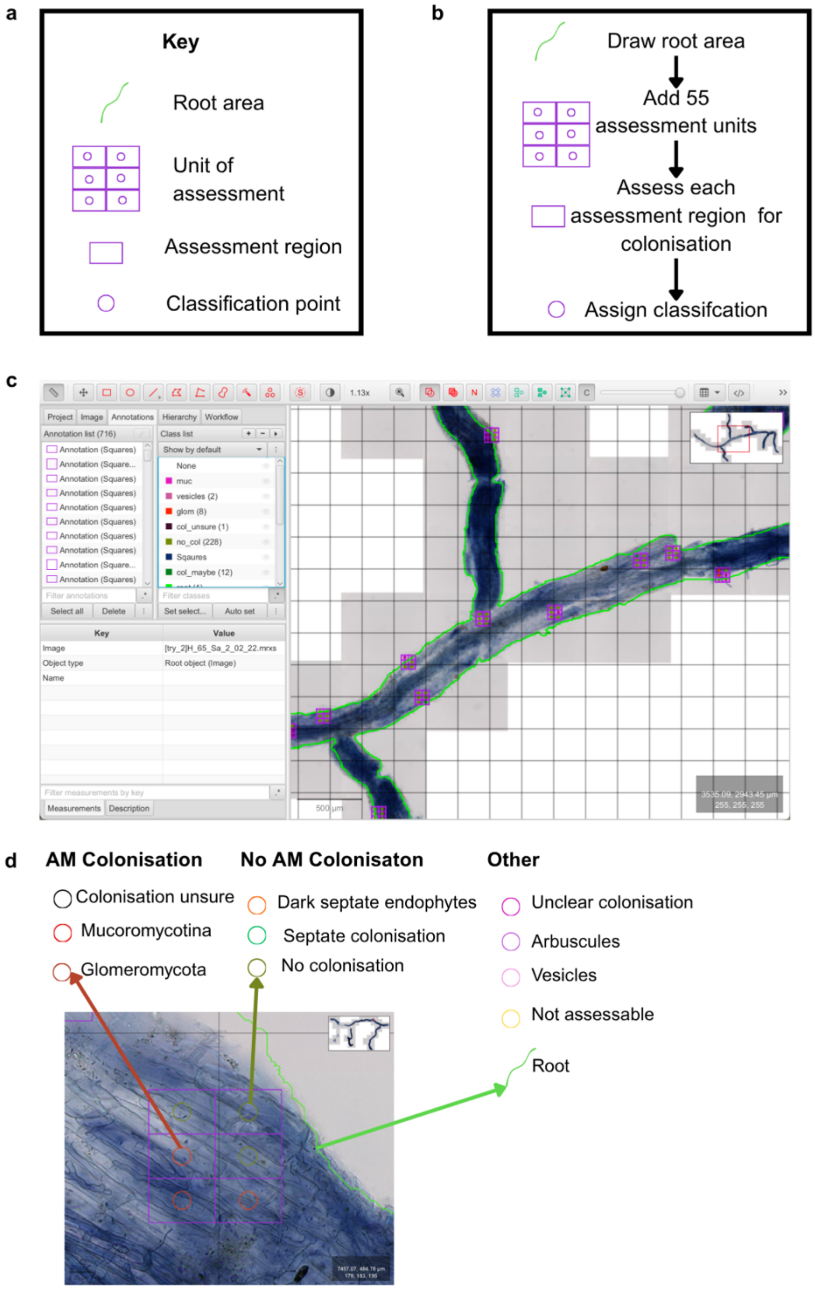

***Figure S 9*** Workflow for assessing arbuscular mycorrhizal (AM) colonisation of stained roots in QuPath (v0.6.0). (a) Annotation key showing the classification categories used during colonisation assessment. (b) Overview of the assessment workflow. Each root section is first delineated and assigned the “Root” classification. A custom QuPath script then randomly distributes 55 assessment units within the annotated “Root”. Each assessment unit comprises six assessment regions (50 × 20 µm), which are examined for the presence of AM colonisation structures. The central classification point is then assigned to a predefined annotation category based on the observations. (c) Screenshot of the QuPath interface showing an annotated root with randomly distributed assessment units. (d) Example of an individual assessment unit, illustrating assessment regions surrounding the central classification point, with examples of classifications assigned as Glomeromycota or No colonisation. Additional annotation categories are shown in panel (a).

***Table S 28*** Predefined annotation categories assigned to regions of stained roots during colonisation assessment. Colours of classification category match the colour of classification circles within the image examples.

| **Classification** | **Example** | **AM fungal colonisation** | **Features** |
|---|---|---|---|
| Glomeromycota | 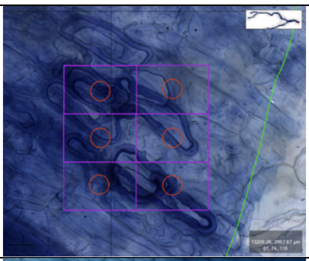 | Yes | Thick tubular aseptate hyphae. These often form coils inside/around cells. |
| Mucoromycota (Endogonales) | 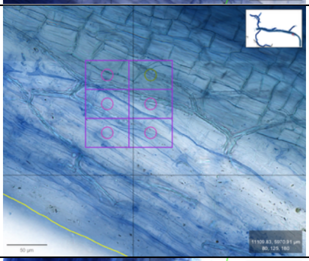 | Yes | Thin networks of aseptate hyphae, can often be highly branched. |
| Arbuscules | 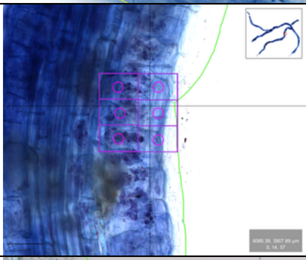 | Yes | Arbuscules appear as clusters of stain (often blotchy) within cells. Defined, “tree-like” structures inside cells are not often observed due to image-based assessment. These were often defined only when aseptate hyphae were observed nearby. |
| Vesicles | 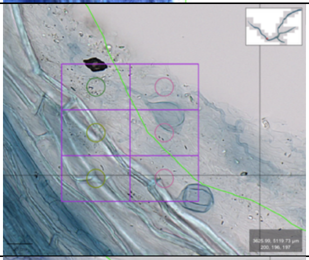 | Yes | Vesicles were classified as AM vesicles if they were attached to aseptate hyphae. |
| Potential colonisation | 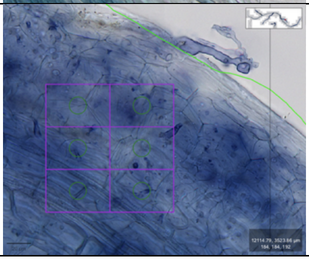 | No (added in some models) | Hyphae are present; however, image does not show clearing the presence/absence of septa or staining is not defined enough to be sure of colonisation. |

| | | | |
|---|---|---|---|
| Dark septate endophytes (DSE) | 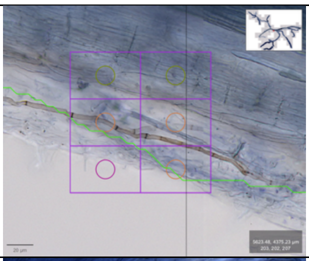 | No | These do not stain blue, instead appear brown due to melanisation. Clear septate hyphae are observed. |
| Septate colonisation | 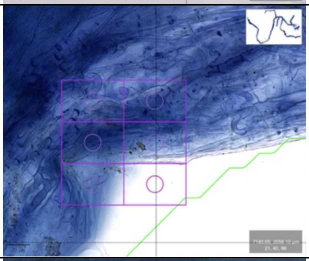 | No | Septate hyphae visible, therefore not AM. |
| Uncolonized | 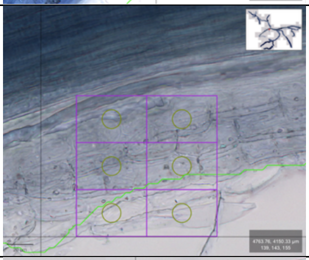 | No | No hyphae, or observable stained structures. |
| Not assessable | 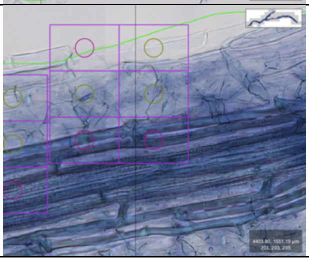 | No | Root region was not assessed if image was out of focus, too dark, dirty, or if assessment region was outside of root area. |

#### References

1. Bankhead, P., Loughrey, M. B., Fernández, J. A., Dombrowski, Y., McArt, D. G., Dunne, P. D., McQuaid, S., Gray, R. T., Murray, L. J., Coleman, H. G., James, J. A., Salto-Tellez, M., & Hamilton, P. W. (2017). QuPath: Open source software for digital pathology image analysis. Scientific Reports, 7(1), 16878. https://doi.org/10.1038/s41598-017-17204-5
2. White, T. J., Bruns, T. D., Lee, S. B., & Taylor, J. W. (1990). Amplificationand direct sequencing of fungal ribosomal RNA genes for phyloge-netics. In M. A. Innis, D. H. Gelfand, J. J. Sninsky, & T. J. White (Eds.),PCR protocols: A guide to methods and applications (pp. 315–322). NewYork, NY: Academic Press
3. Tedersoo, L., & Anslan, S. (2019). Towards PacBio-based pan-eukaryote metabarcoding using full-length ITS sequences. Environmental Microbiology Reports, 11(5), 659-668. https://doi.org/10.1111/1758-2229.12776

4. Tedersoo, L., & Lindahl, B. (2016). Fungal identification biases in microbiome projects. Environ Microbiol Rep, 8(5), 774-779. https://doi.org/10.1111/1758-2229.12438

5. Percival-Alwyn, L., Barnes, I., Clark, M.D., Cockram, J., Coffey, M.P., Jones, S., Kersey, P.J., Kidner, C.A., Kosiol, C., Li, B., Marsh, W.A., Zhou, J., Caccamo, M., Milne, I. (2024). UKCropDiversity-HPC: A collaborative high-performance computing resource approach for sustainable agriculture and biodiversity conservation. Plants, People, Planet. DOI: 10.1002/ppp3.10607
